# Characterisation of an *Escherichia coli* line that completely lacks ribonucleotide reduction yields insights into the evolution of obligate intracellularity

**DOI:** 10.1101/2022.09.28.509999

**Authors:** Samantha D. M. Arras, Nellie Sibaeva, Ryan J. Catchpole, Nobuyuki Horinouchi, Dayong Si, Alannah M. Rickerby, Kengo Deguchi, Makoto Hibi, Koichi Tanaka, Michiki Takeuchi, Jun Ogawa, Anthony M. Poole

**Author notes:** RJC: Department of Biochemistry and Molecular Biology, University of Georgia, Athens, GA, USA NH: Amano Enzyme Inc., Nagoya, Aichi, Japan DS: State Key Laboratory of Animal Nutrition, Laboratory of Feed Biotechnology, College of Animal Science and Technology, China Agricultural University, Beijing, China MH: Department of Biotechnology, Biotechnology Research Center, Toyama Prefectural University, Toyama, Japan KT: Department of Nutritional Science, Okayama Prefectural University, Soja, Okayama, Japan MT: Industrial Microbiology, Graduate School of Agriculture, Kyoto University, Kyoto, Japan.

## Abstract

All life requires ribonucleotide reduction for *de novo* synthesis of deoxyribonucleotides. A handful of obligate intracellular species are known to lack ribonucleotide reduction and are instead dependent on their host for deoxyribonucleotide synthesis. As ribonucleotide reduction has on occasion been lost in obligate intracellular parasites and endosymbionts, we reasoned that it should in principle be possible to knock this process out entirely under conditions where deoxyribonucleotides are present in the growth media. We report here the creation of a strain of *E. coli* where all three ribonucleotide reductase operons have been fully deleted. Our strain is able to grow in the presence of deoxyribonucleosides and shows slowed but substantial growth. Under limiting deoxyribonucleoside levels, we observe a distinctive filamentous cell morphology, where cells grow but do not appear to divide regularly. Finally, we examined whether our lines are able to adapt to limited supplies of deoxyribonucleosides, as might occur in the evolutionary switch from *de novo* synthesis to dependence on host production during the evolution of parasitism or endosymbiosis. Over the course of an evolution experiment, we observe a 25-fold reduction in the minimum concentration of exogenous deoxyribonucleosides necessary for growth. Genome analysis of replicate lines reveals that several lines carry mutations in *deoB* and *cdd. deoB* codes for phosphopentomutase, a key part of the deoxyriboaldolase pathway, which has been hypothesised as an alternative to ribonucleotide reduction for deoxyribonucleotide synthesis. Rather than synthesis via this pathway complementing the loss of ribonucleotide reduction, our experiments reveal that mutations appear that reduce or eliminate the capacity for this pathway to catabolise deoxyribonucleotides, thus preventing their loss via central metabolism. Mutational inactivation of both *deoB* and *cdd* is also observed in a number of obligate intracellular bacteria that have lost ribonucleotide reduction. We conclude that our experiments recapitulate key evolutionary steps in the adaptation to intracellular life without ribonucleotide reduction.

## Introduction

All life on our planet requires ribonucleotide reduction for *de novo* synthesis of deoxyribonucleotides, from ribonucleotides^1,2^. The emergence of ribonucleotide reduction likely drove the transition from RNA to DNA early in the evolution of life^3-6^ and its centrality is reflected in its near ubiquity in organismal genomes^2^. Structural analyses indicate that the catalytic cores of ribonucleotide reductases (RNRs) are structurally homologous^3,7^, despite sequence similarities between them being extremely limited^8^. RNRs catalyse deoxyribonucleotide synthesis via a common mechanism wherein a cysteinyl free radical is generated in the active site. While this similarity in catalysis underscores a common evolutionary origin, RNRs have diverse mechanisms for radical generation^1^. Three broad, evolutionarily distinct, families exist which are also divided on the basis of their radical generation mechanism^1,3,9^. This divergence likely coincides with adaptation to varied environments^9,10^: class I enzymes use a radical generation mechanism based on an oxygen-containing metal ion centre and are strictly aerobic, while class III enzymes generate a stable glycine radical via cleavage of S-adenosyl methionine, and are strictly anaerobic. Class II enzymes generate a 5**’**-deoxyadenosyl radical through cleavage of adenosylcobalamin and operate irrespective of oxygen presence or absence.

To date, there is only one type of environment known where ribonucleotide reduction is dispensable, albeit indirectly: some obligately intracellular species lack genes for ribonucleotide reduction and instead rely on their hosts for deoxyribonucleotides^2^. Examples include a strain of *Buchnera aphidicola str. Cc* from the Cedar bark aphid, *Cinara cedri*^11^. *Buchnera aphidicola* are maternally inherited aphid endosymbionts that synthesise amino acids in short supply in the aphid diet and appear to be on the path to transitioning to an organelle^12,13^. Two obligate intracellular pathogens—*Ureaplasma urealyticum*^14^ and *Borrelia burgdorferi*^15^*—* have also lost genes for ribonucleotide reduction^2^.

While ribonucleotide reduction is essential, we reasoned that it should be possible to dispense with genes for ribonucleotide reduction if deoxyribonucleotides or their precursors are available *via* growth media. We report the successful elimination of all three RNR operons from the bacterium *E. coli*. The resulting strain is dependent on deoxyribonucleosides in the growth medium but cannot grow if the media are supplemented with deoxyribose plus the four nucleobases (A, G, C, T). To understand the impact of limited deoxyribonucleoside availability, we subjected our knockout line to experimental evolution followed by genome sequencing. We were interested in establishing whether, under such conditions, our lines would utilise the reverse deoxyriboaldolase pathway for deoxyribonucleotide synthesis. This pathway is reversible *in vitro*^16,17^, and has long been considered a plausible alternative to ribonucleotide reduction^18,19^ for dNTP synthesis, with some considering it to be a plausible ancestral route for the origin of DNA^20,21^. Our evolution experiments reveal that this pathway is not coopted for deoxyribonucleotide synthesis following loss of ribonucleotide reduction. Rather, it is a liability under conditions of limited deoxyribonucleotide availability; we observe loss-of-function mutations in *deoB*, which are predicted to prevent catabolism of 2-deoxy-D-ribose-1-phosphate. Available genome sequence data from obligate intracellular bacteria that lack ribonucleotide reduction indicate that *deoB* has in fact been lost on multiple occasions, suggesting that recycling of deoxyribose is disadvantageous under conditions where this sugar is in limited supply; deletion of *deoB* would enable this sugar to be rerouted for deoxyribonucleotide synthesis but would also preclude *de novo* synthesis via the reverse deoxyriboaldolase reaction. Our results thus illuminate a key adaptive step taken by obligate intracellular species to mitigate the loss of ribonucleotide reduction.

## Methods

### Strains and growth conditions

All *E. coli* strains used in this study are listed in **Supplementary Table 1**. *E. coli* B strain REL606^22^ was used as the wild type strain for all experiments. Strains were grown in LB (1% tryptone, 1% NaCl and 0.5% yeast extract +/-2% agar) with the addition of 1 mg/mL each of the four deoxyribonucleosides (dNS—deoxyguanine (dG), deoxyadenine (dA), deoxycytosine (dC) and deoxythymine (dT) (AK Scientific)) for mutant strains, unless stated otherwise. For evolution experiments, strains were grown in 1x MOPS media^23^ supplemented with dNS, as specified. All experiments were conducted in the presence of 100 µg/mL streptomycin.

### Creation of a ribonucleotide reductase mutant

Deletion of RNR operons was carried out using a scarless genome engineering method^24^ in series (**Supplementary Text 1**): the *nrdDG* operon (class III RNR) was deleted first, followed by *nrdHIEF* (class IB) then *nrdAB* (class IA). For each operon, 500 bp upstream and downstream from the start and stop codon was PCR amplified. A second round of overlap PCR combined fragments together and the three resulting product was cloned into pST76a. The resulting construct was transformed into wild type at 30°C, then with genome integration induced by increasing temperature to 42°C. Helper plasmid (pSTKST) was transformed into the resulting strain, and deletion was induced with chlortetracycline. Successful deletion was confirmed using PCR across the operon (Primers used in this study are listed in **Supplementary Table 2**). The resulting strain, which we call ΔRNR, was confirmed by whole genome sequencing to lack all three RNR operons (Δ*nrdAB*, Δ*nrdHIEF*, Δ*nrdDG*).

### Total RNA extraction and RT-PCR

To confirm absence of ribonucleotide reductase expression, strains were grown overnight in LB and diluted to 1:100 in 10 mL fresh media after reaching stationary phase. The cultures were then grown for ∼3 hours and total RNA was isolated from mid-log phase cultures using TRIzol Max Bacterial Isolation kit (ThermoFisher). RNA was quantified using a Qubit 4.0 fluorometer. Purified RNA was diluted to 300 ng/mL and treated with TURBO DNaseI (Ambion). This DNA-free RNA was then subjected to RT-PCR using the SuperScript III One-Step RT-PCR system with Platinum Taq DNA Polymerase kit (Invitrogen) with primers specific to the gene of interest (**Supplementary Table 2**).

### Growth assays

Bacterial strains were retrieved from -80°C stocks and grown overnight in LB (1 mg/ml dNS was added to media for growth of ΔRNR lines). Cultures were washed twice in 1x PBS and 10 µL equivalent was added to fresh 24-well plates containing 1 mL of 1x MOPS + 1% glucose. Differing concentrations of dNS were added from a 20 mg/mL stock solution to each individual well. Growth was monitored for 24-48 hours, taking measurements (OD_595_) every 6 minutes using a FLUOstar Omega Microplate Reader (BMG Labtech). All experiments were performed in triplicate.

### Microscopy

Overnight cultures were grown at 37°C in MOPS media (with dNS added as required). Cultures were twice spun down and washed in 1x PBS. 10 µl of culture was aliquoted onto a microscope slide. Brightfield images were taken using a LEICA ICC50 W microscope (Leica, v.3.2.0) and imported to Photoshop (v 22.4.2) for cell-length measurement. Cell length measurements were determined for each strain by averaging from 20 observations (length of the first 5 cells from top left to right were counted from each of 4 images).

To visualise DNA, we used a modified version of a FITC/DRAQ-5 double-staining protocol (Silva et al., 2010), instead using DAPI in place of DRAQ-5. Coverslips were coated in poly-D-lysine and placed at 37°C overnight. Coverslips were washed twice with water, dried and stored at 4°C until ready for use. Overnight cultures were washed in 1x PBS, resuspended, and placed on a poly-D-lysine coated coverslip at 37°C for 1 hour. Cells were washed with PBS, and fixed using 4% paraformaldehyde at room temperature for 10 minutes. Coverslips were washed with 1x PBS, followed by 1% Triton X-100 for 5 minutes. Following a further wash with 1x PBS, coverslips were then incubated in 1x PBS containing 6 µg/mL FITC for 30 minutes at 37°C. The coverslips were washed again with 1x PBS, then suspended in 1x PBS containing 5 µg/mL DAPI and placed in the dark for 10 minutes. Coverslips were lastly washed twice more in 1x PBS, then were mounted on microscope slides. Strains were visualised on a Nikon Ni-E Fluorescence microscope using fluorescent filter cubes for DAPI and FITC, and a 100x oil objective lens. Images were overlaid using Nikon NIS-Elements software.

### Experimental Evolution

All evolution experiments were performed at 37 °C. Original bacterial strains were retrieved from -80 °C stocks and grown overnight in LB (1mg/ml dNS was added to media for growth of ΔRNR lines). Cultures were then washed twice in 1x PBS and 50 µL equivalent was added to fresh 6-well plates containing 5 mL 1x MOPS media (1% glucose) supplemented with the dNS concentration required for each condition (1 mg/ml, 0.25 mg /ml, 0.01 mg/mL). 5 lineages of ΔRNR and 3 for WT (*E. coli* REL606^22,25^) were passaged for each condition until transfer 30. Line 1 of ΔRNR grown at 0.25 mg/ml (ΔRNR_T30_250_1) was serially passaged as ten independent lines for an additional 10 transfers in MOPS+1% glucose and 0.01 mg/ml dNS (ΔRNR_T40_10_1-10). Plates were left to reach stationary with agitation at 120 rpm. Approximately 10^7^ – 10^8^ of wash cells were transferred to a fresh well. A glycerol stock was created for each line every 5 transfers. PCR contamination checks were performed every 5 transfers.

### Sequencing and genome assembly

All strains and lineages required for sequencing were streaked for single colonies on LB media, with a single colony being used to inoculate an overnight culture. Genomic material was isolated using 20µg genomic tips (Qiagen) as per manufacturer**’**s instructions and DNA quantified using a Qubit 4.0 fluorometer (ThermoFisher). Libraries were generated using the NEXTFLEX rapid XP DNA-seq kit (PerkinElmer) and NEXTFLEX UDI Barcodes. Strains were sequenced to approximately 50-fold coverage at Auckland Genomics Facility using an Illumina MiSeq Platform with 2x 150 bp paired-end reads. Reads were trimmed using the BBDuk^26^ 1.0 plug-in for Geneious Prime 2021.1.1 (https://www.geneious.com). Mapping to the *E. coli* REL606 genome reference was performed using Bowtie2 (v2.3.2)^27^ plug-in (v7.2.1) and visualised using default settings in Geneious. All genome data have been deposited to the SRA database and are available via Bioproject accession number PRJNA882995 (https://www.ncbi.nlm.nih.gov/sra/PRJNA882995).

### Sequence analyses

For creation of the phosphopentomutase alignment, we ran BLASTP with default settings (E-value threshold = 0.1, BLOSUM62, Filtering: none, Gapped: yes, Hits: 1000) against the Uniprot reference genomes plus Swiss-Prot database (https://www.uniprot.org/blast/) using the deoB protein sequence from *E. coli* (REL606) as query. To create our alignment dataset, we filtered results for Swiss-Prot sequences and removed duplicates. We then added the *E. coli* REL606 query sequence and the phosphopentomutase from *Bacillus cereus*, the crystal structure of which has been determined^28^. We generated our alignment with Muscle^29^ using the Geneious plugin (v. 3.8.425) and default settings.

To screen for presence/absence of ribonucleotide reduction, phosphopentomutase and cytidine deaminases in *Buchera, Borrelia* and *Ureaplasma*, we ran blastp searches against the nr_protein database restricted to these taxa using query sequences listed in **Supplementary Table 10**. Where data indicated gene absence, we confirmed this by examining the genome sequences for assembly issues, tblastn searches, and manual inspection of predicted ORFs from the genome assembly files (https://www.ncbi.nlm.nih.gov/genome/browse/#!/overview/). For cases where gene absence was clearly due to the assembly, these were excluded from further analysis. Assembly accession numbers and taxids are provided in **Supplementary Text 2**.

## Results

### Creation of an *E. coli* line lacking ribonuclease reduction

*E. coli* carries genes for three RNRs: aerobic class Ia (encoded by the *nrdAB* operon), class Ib (*nrdHIEF*), and anaerobic class III (*nrdDG*) (**Figure 1**). Under aerobic growth, ribonucleotide reduction is primarily performed by the class Ia enzyme, with the role of the Ib enzyme being less clear^30^.

**Figure 1.**
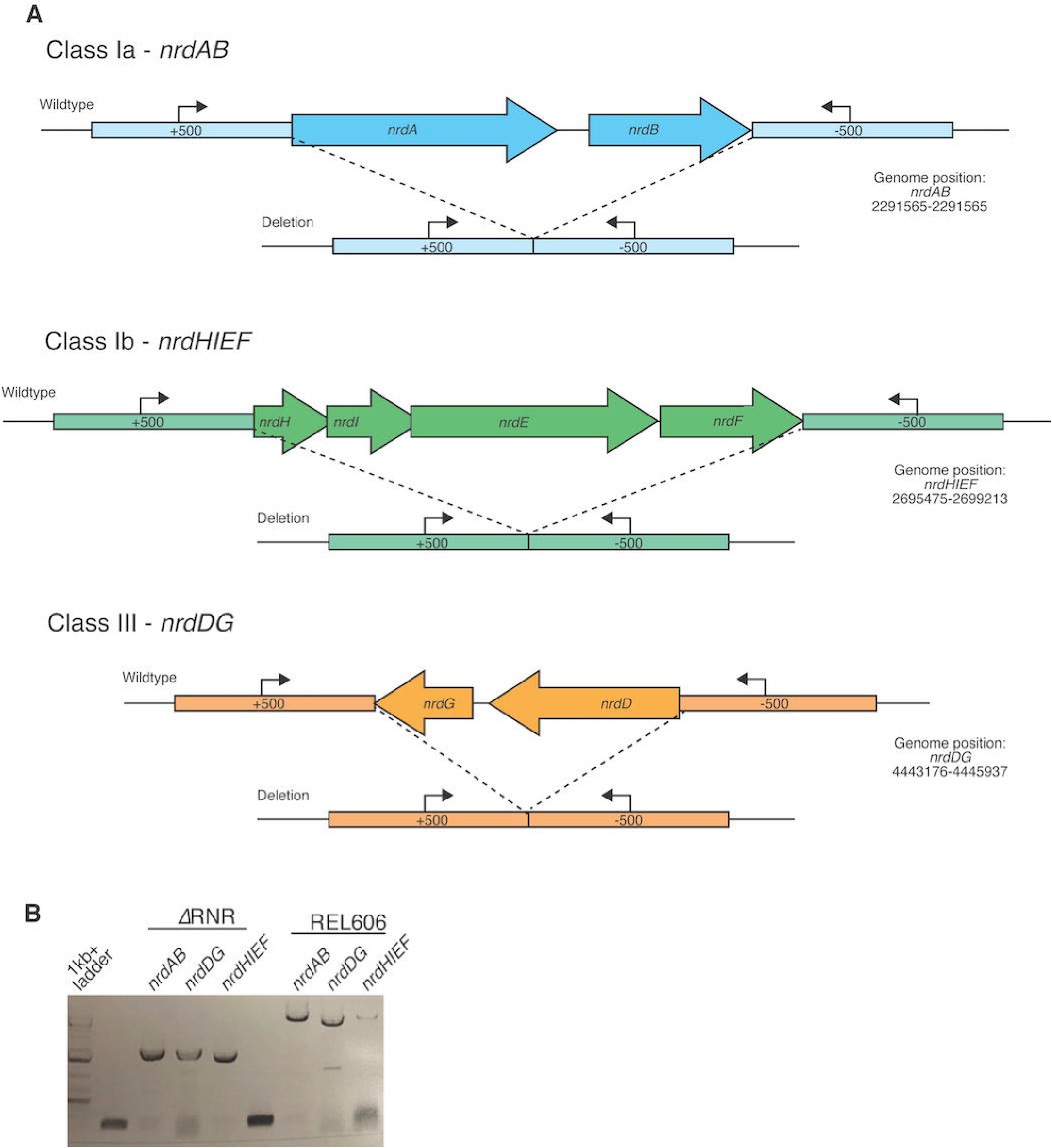
Deletion of ribonucleotide reductase genes in *E. coli*. **A**. Schematic of each of the three RNR operons in *E. coli* and the genomic regions following scarless deletion. Arrows indicate locations of PCR primers (Supplementary Table 2) used to confirm RNR operon presence/absence. **B**. Results of PCR with primers external to all three *nrd* operons in ΔRNR compared with the wild-type progenitor strain (REL606). Lane 1, ladder; Lanes 2 and 6, no template. Lanes 3-5, band sizes consistent with operon deletion (*ΔnrdAB* 1645 bp, *ΔnrdDG* 1620 bp and *ΔnrdHIEF* 1477 bp) in the ΔRNR line. Lanes 6-9, band sizes for all three operons in wild type *E. coli* REL606 (*nrdAB* 5303 bp, *nrdDG* 4382 bp and *nrdHIEF* 5191 bp).

We reasoned that, while ribonucleotide reduction is an essential process, it should be possible to compensate for its loss via media supplementation. To eliminate ribonucleotide reduction, we sequentially deleted the RNR operons using a scarless deletion protocol (Methods). We first deleted *nrdDG*, as this operon is known not to be expressed under aerobic conditions. We next deleted the *nrdHIEF* operon from the resulting *ΔnrdDG* line. Finally, we attempted to delete the *nrdAB* operon the *ΔnrdHIEF ΔnrdDG* line, grown in the presence of deoxyribonucleosides (dNS). We were initially unsuccessful in deleting *nrdAB*. This is not unexpected as, while deoxyribonucleosides are known to be taken up, some (dG, dC, dA) are unavailable for DNA synthesis, even when genes in the DERA pathway are mutated, preventing their catabolism^31^. We reasoned that may be because of the absence of a suitable deoxynucleoside kinase activity for converting those deoxyribonucleosides to deoxyribonucleotides. We therefore tried knocking out *nrdAB* in the presence of a heterologously expressed deoxyadenosine kinase (dAK) gene from *Mycoplasma mycoides*. Mycoplasmas are known to be dependent on salvage for dNTP production and possess deoxyadenosine kinases genes that permit utilisation of deoxyribonucleosides for DNA synthesis^32^. To establish if resulting lines lacked all three *nrd* operons, we first PCR screened for evidence of chromosomal *nrd* operon deletions. Our results indicate that all three operons were successfully deleted (**Figure 1**). As deletion does not exclude relocation of functional gene copies to another genomic location, we confirmed gene absence via PCR using primers internal to *nrd* genes and performed RT-PCR to confirm absence of gene expression (**Supplementary Figure 1**). Finally, knockout status was confirmed with whole genome sequencing. This confirmed deletion of all genes for ribonucleotide reduction, with the *nrdAB* operon successfully deleted under heterologous expression of dAK from *M. mycoides* (**Supplementary Text 1; Supplementary Figure 2**). Of four knockout lines, we selected one isolate (hereafter called ΔRNR) for all subsequent work. In addition to lacking all *nrd* genes, genome sequencing revealed 23 SNPs that likely appeared during the creation of ΔRNR from wild-type progenitor line (REL606) (**Supplementary Table 3**).

**Figure 2.**
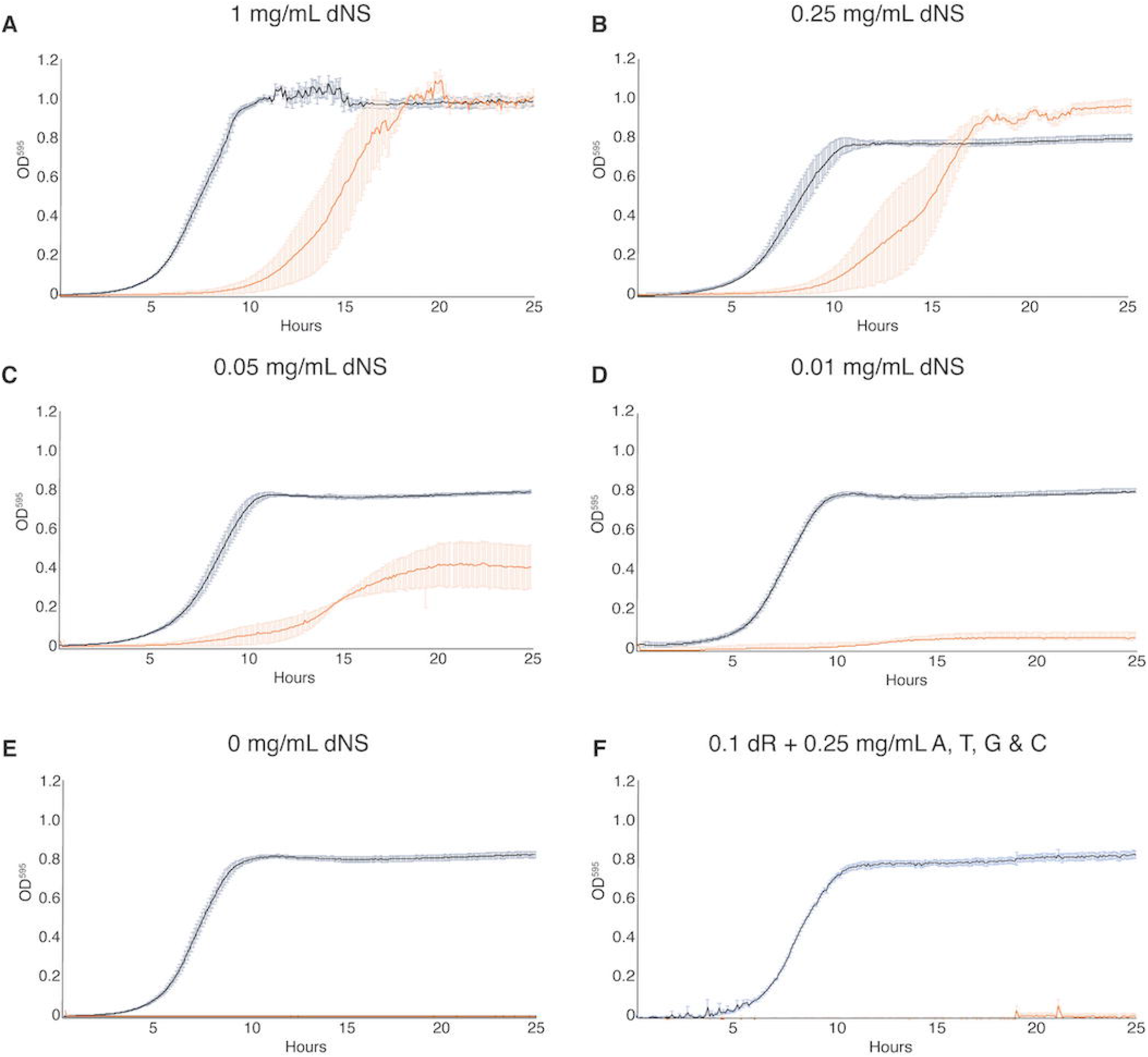
ΔRNR shows growth across a range dNS concentrations but cannot grow in the absence of dNS supplementation. Growth was monitored for 25 hours in 1x MOPS media + 1% glucose with supplementation, as indicated. Wild-type (REL606, black), ΔRNR (orange). Curves show mean OD_595_, error bars show SEM, all experiments performed in triplicate. **A**. 1mg/mL, **B. 0**.25 mg/mL, **C. 0.0**5 mg/mL, **D. 0.0**1 mg/mL & **E**. 0 mg/mL dNS. **F**. 0.1 mg/mL deoxyribose (dR) + 0.25 mg/mL of adenine (A), cytosine (C), thymine (T) and guanine (G).

### ΔRNR is dependent on deoxyribonucleoside supplementation

Our ΔRNR line was created in the presence of dNS supplementation, but this does not mean it is dependent on dNS supplementation for growth. We therefore sought to understand what supplementation, if any, our ΔRNR line requires. To determine the lowest concentration of dNS that permits growth, we generated a series of growth curves for differing dNS concentrations. At high dNS concentrations (1 mg/mL), ΔRNR grows favourably, though shows a clear lag compared to wild-type on equivalent media (**Figure 2A**). At lower dNS concentrations, ΔRNR growth is impaired, but there is still discernible growth at 0.05 mg/mL (**Figure 2C**). When this is dropped to 0.01 mg/mL we observed only marginal growth of ΔRNR (**Figure 2D**). As expected, in the absence of dNS supplementation no growth is observed, while wild-type lines are unaffected (**Figure 2E**). Finally, we tried growing ΔRNR on deoxyribose (dR) plus each of the four bases (A, G, C, T). No growth was observed (**Figure 2F**). These results indicate that deletion of the three *nrd* operons from ΔRNR has completely eliminated the capacity for *de novo* deoxyribonucleotide synthesis; no other genes appear able to compensate for this deficiency.

We next sought to establish whether ΔRNR requires supplementation of all four dNSs, or whether any are dispensable. Strains were once again grown in 1x MOPS+1% glucose, with individual deoxyribonucleosides (dA, dG, dT or dC) added at a concentration of 0.25 mg/mL. This revealed that ΔRNR is unable to grow in the presence of dA, dG, or dT (either alone (**Figure 3A-C**), or in combination (**Figure 3E**), even after 44 hours of monitoring (data not shown). However, ΔRNR can grow on dC alone **(Figure 3D)**. Together these data indicate deoxycytidine (dC) is the sole deoxyribonucleoside required for ΔRNR growth in minimal media.

**Figure 3.**
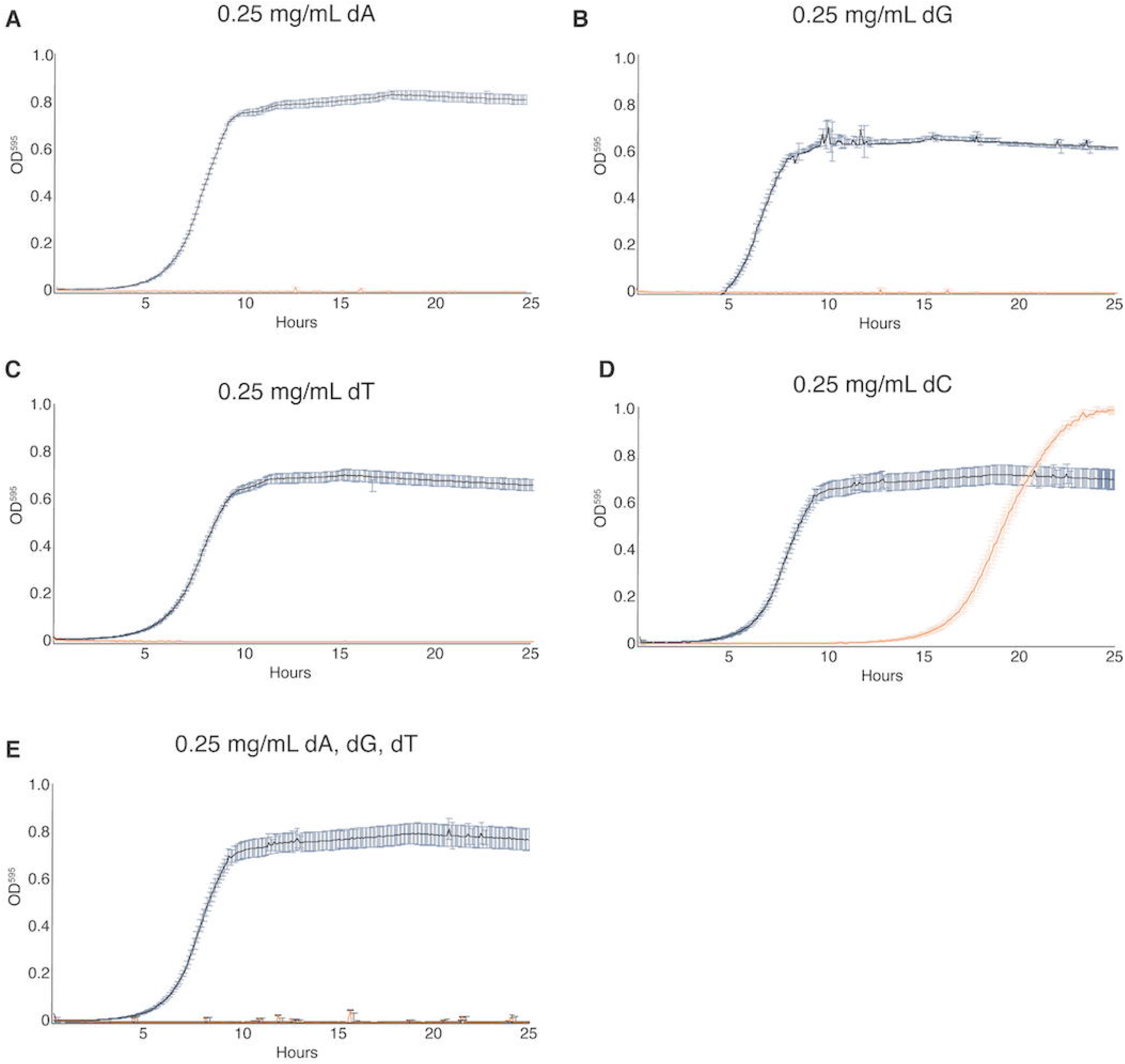
ΔRNR requires only deoxycytidine for growth in minimal media. Growth was monitored for wild-type (REL606, black) and ΔRNR (orange) in 1x MOPS media + 1% glucose, with dNS supplementation, as indicated. Growth was monitored for 25 hours. Curves show mean OD_595_, error bars show SEM, all experiments performed in triplicate. **A**. dA (0.25 mg/mL), **B**. dG (0.25 mg/mL), **C**. dT (0.25 mg/mL), **D**. dC (0.25 mg/mL) **E**. dA, dG, dT, each at 0.25 mg/mL

### ΔRNR exhibits a filamentous cell morphology when grown under limiting dNS

During our growth assays we observed a **‘**clumping**’** phenotype when ΔRNR is grown in liquid media at concentrations of dNS that limit growth. This contrasts with the uniform cloudy appearance of wild-type *E. coli* (**Figure 4C, top left panel**). Examination of cells at 100x magnification revealed that ΔRNR cells are elongated and filamentous at very low levels of dNS, whereas ΔRNR cells have a similar morphology to wild-type at higher dNS concentrations (**Figure 4A**). Some cells reached lengths several times that of wild-type (**Supplementary Table 4**).

**Figure 4.**
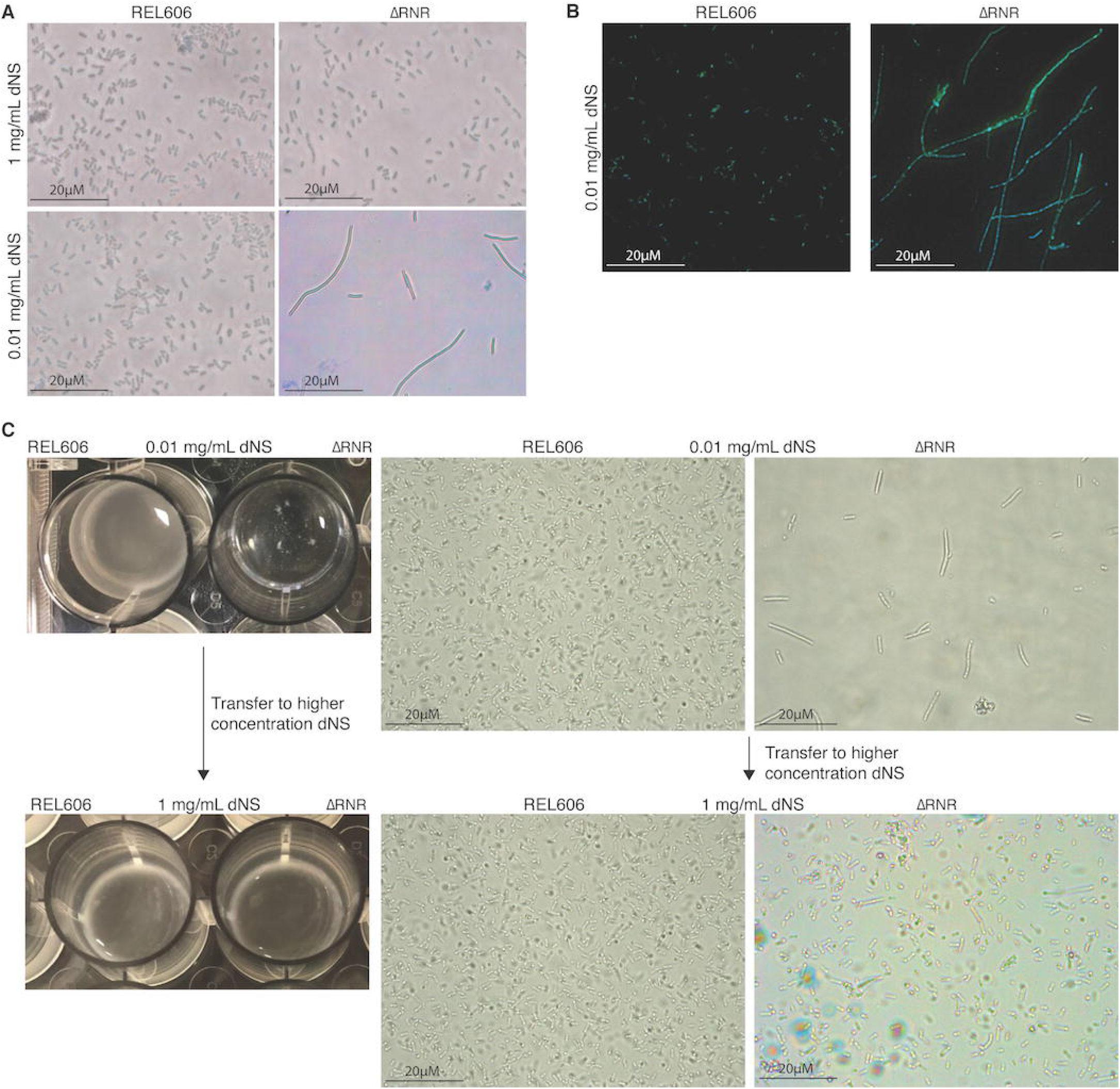
ΔRNR exhibit a reversible elongated, filamentous cell morphology with multiple nucleoids when grown in limiting dNS concentrations. **A**. 100x Brightfield photographs of REL606 and **Δ**RNR at 1 and 0.01 mg/mL **B**. FITC and DAPI staining at 100x of REL606 and **Δ**RNR at 0.01 mg/mL **C. Δ**RNR phenotype is reversible if dNS concentration is increased. Loss of clumpy phenotype occurs when **Δ**RNR cells are transferred from low (0.01 mg/mL) to high (1 mg/mL) dNS. Brightfield microscopy images reveal that this change is accompanied by a reduction in elongated cell morphology upon transfer to high [dNS]. (Magnification 1000X; Scale bar 20 µM)

This phenotype appears most pronounced at low dNS concentrations. We therefore sought to establish if this phenotype is the result of environmental conditions. We transferred a population of ΔRNR cells exhibiting the filamentous cell phenotype from low dNS (0.01 mg/mL) media to a less restrictive environment (dNS = 1 mg/mL). Following transfer to higher [dNS], the filamentous phenotype is heavily diminished (**Figure 4C)**. This demonstrates that the phenotype is environmental and can be reversed if cells are moved to a high dNS environment.

One possibility is that, under low [dNS], cells are growing but unable to complete cell division. If so, this might be reflected by the presence of multiple DNA-dense regions across the length of the cells. To visualise DNA within filamentous cells, we stained ΔRNR cells grown in low dNS (0.01 mg/mL) with FITC and DAPI. This revealed the presence of multiple DAPI-stained regions across the length of the cells **(Figure 4B)**, suggestive of the presence of multiple DNA nucleoids in ΔRNR cells grown at low [dNS].

### Evolution of ΔRNR lines under restricted dNS supplementation

The loss of ribonucleotide reduction in obligate intracellular lifestyles presumably resulted from relaxed selection on deoxyribonucleotide production when deoxyribonucleotides are available from the environment. However, for both parasitism and endosymbiosis, the host dNTP pool must be shared. We were therefore interested to assess if the ΔRNR line adapts to a reduction in dNS availability, as might occur in the evolutionary switch from *de novo* synthesis to dependency on host production.

In order to allow the strains to adapt to a lower concentration of dNS supplementation in the growth media, we initiated evolution experiments at dNS concentrations where ΔRNR lines grow, but not as well as wildtype (1 mg/mL and 0.25 mg/ml). Five independent lines of ΔRNR were serially passaged in each condition by growing the culture to stationary phase, then transferring 50 µL into a new 6-well plate containing 5 mL of fresh MOPS media+ 1% glucose and dNS (**Figure 5**). We also established three control lines of the wild-type progenitor strain (REL606) at each dNS concentration. The initial experiment was run for a total of 30 transfers. Glycerol stocks were created every 5 transfers for each line, with contamination checks performed concurrently.

**Figure 5.**
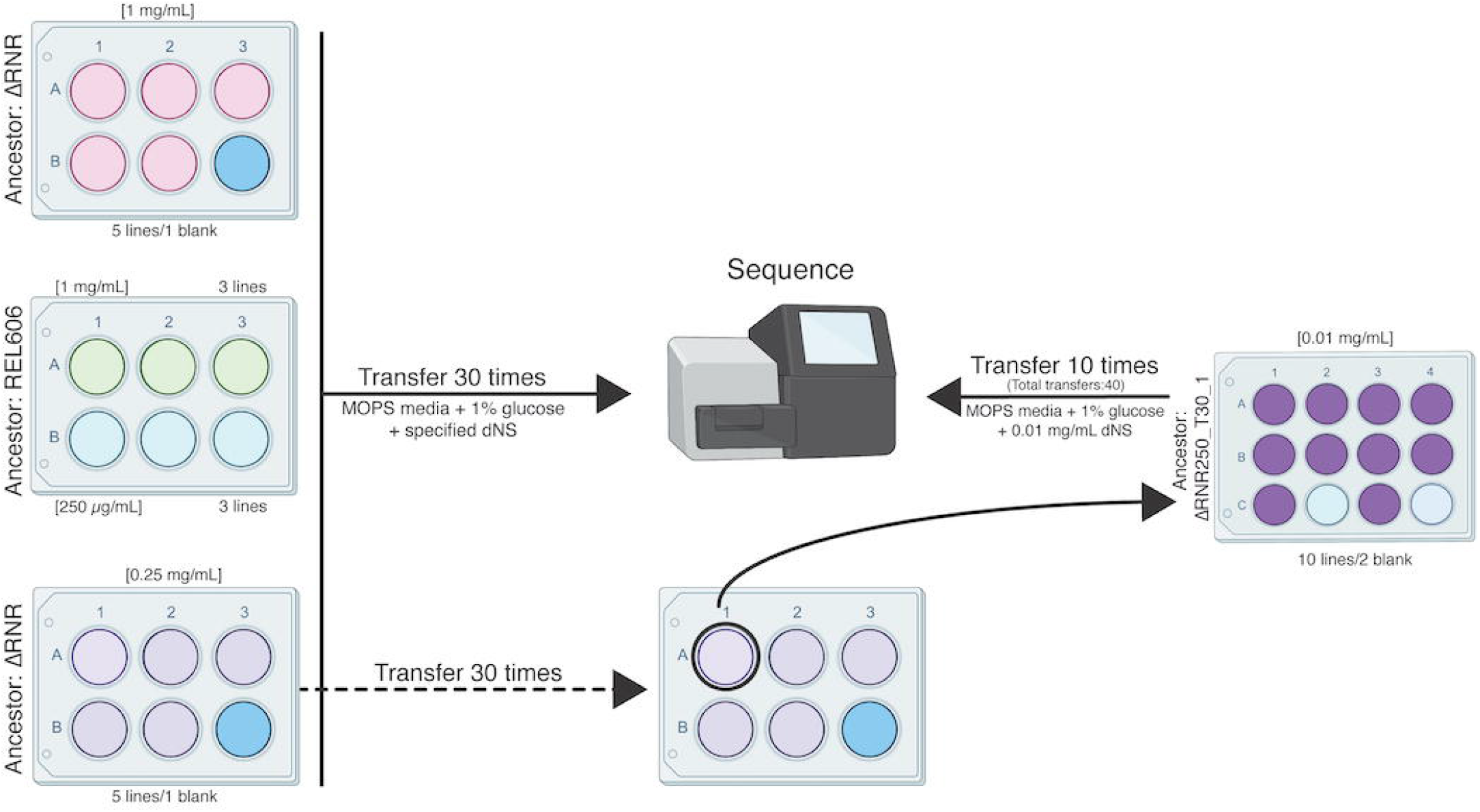
Overview of the ΔRNR evolution experiment. Five lines of ΔRNR and three lines of wild type progenitor (REL606) were established at one of two conditions (1 mg/mL or 0.25 mg/mL dNS in MOPS+1% glucose), and serially passaged for 30 transfers. Genomic material from each line was then extracted and sent for sequencing. To further investigate adaptation to low concentrations of dNS, the **“**fittest**”** ΔRNR line grown at 0.25 mg/mL dNS (ΔRNR_250_T30_1) was used to seed a subsequent experiment. Ten replicate lines of ΔRNR_250_T30 _1 were serially passaged for an additional 10 transfers in MOPS+1% glucose and 0.01 mg/ml dNS. DNA from each of these 10 lines (ΔRNR_10_T40_1-10) was then extracted and sent for whole genome sequencing.

### Elongate cell morphology in ΔRNR lines diminishes over the course of the evolution experiment

Our initial observations of unevolved ΔRNR (henceforth ΔRNR_T0) revealed an elongated filamentous cellular phenotype at lower concentrations of dNS (**Figure 4**). We therefore monitored cell morphology during our evolution experiment. Every 5 transfers, we examined length of ΔRNR and REL606 cells from each condition (1 mg/mL or 0.25 mg/mL dNS; cells from well position A1 (ΔRNR) and A1 or B1 (REL606) were used for all measurements, and mean length calculated). At the beginning of the experiment, ΔRNR cells are substantially longer than wild-type, particularly at 0.25 mg/mL dNS (**Figure 6**). We recorded cell lengths of 20 µM at this lower concentration (average of 6 µM), a staggering 10-20 times the length of wild-type cells. By the time the experiment had reached 30 transfers, average cell length was comparable to wild type at both dNS concentrations (**Figure 6A**). These changes in gross morphology suggest cells had begun to adapt to restricted dNS availability.

**Figure 6.**
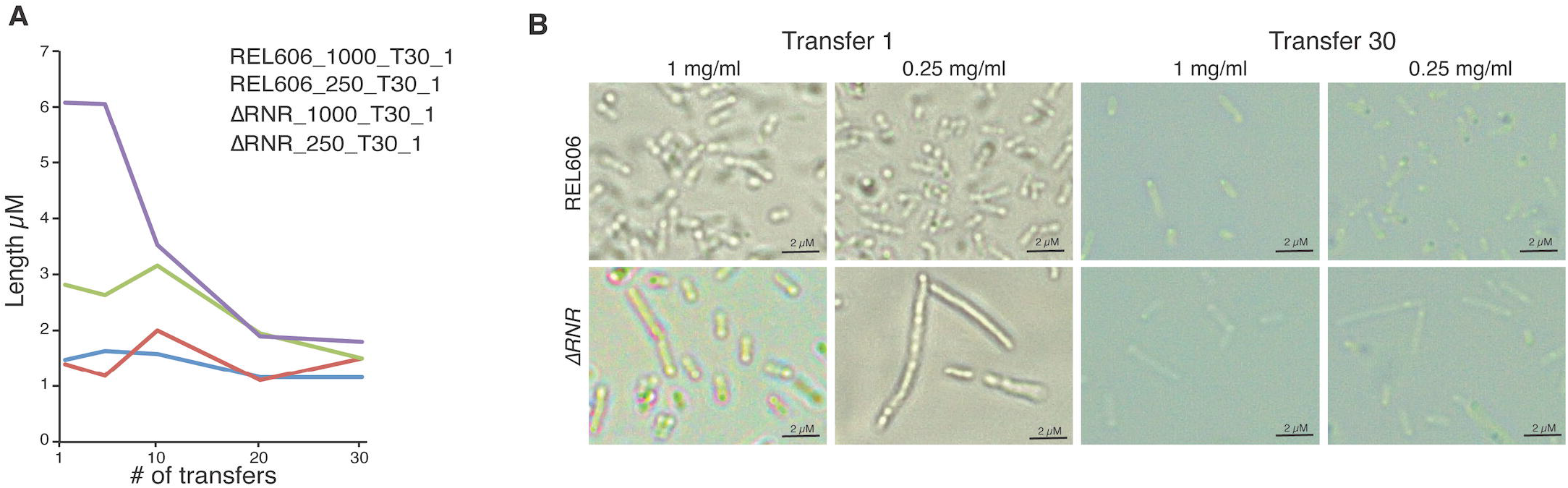
The ΔRNR elongate cell phenotype diminishes over the course of an evolution experiment. **A**. Mean cell length during the course of a 30-transfer evolution experiment. Cell length was determined for 20 cells for replicate line 1 of ΔRNR and REL606 evolved in either 1 mg/mL or 0.25 mg/mL dNS. Cell measurements were taken every 5 transfers. Over the course of the evolution experiment ΔRNR cell length gradually reduces at both dNS concentrations. **B**. Morphology of wild-type (top row) and ΔRNR (bottom row) cells at Transfer 1 and Transfer 30 at each dNS concentration (1 mg/mL or 0.25 mg/mL). (Magnification 1000X; Scale bar 2µM)

### ΔRNR lines exhibit improved growth following evolution under restricted dNS availability

After 30 transfers, we sought to determine if ΔRNR lines had improved their capacity to grow under restricted (either 1 mg/mL or 0.25 mg/mL) dNS availability. We use the following nomenclature for our evolution lines: ΔRNR_[dNS] _transfer#_line#, so ΔRNR1000_T30_L1 is replicate line #1 of ΔRNR evolved at 1 mg/mL (i.e. 1000 µg/mL) dNS for 30 transfers. At 1 mg/mL, ΔRNR1000_T30_L1 through L5 all showed improved growth relative to ΔRNR_T0 (**Figure 7A**). Moreover, compared to ΔRNR_T0, which exhibited almost no growth at 0.01 mg/ml dNS (**Figure 2D**), all evolved lines (ΔRNR1000_T30_L1-5) showed improved growth at this low concentration (**Figure 7D**). A similar overall pattern of improved was also seen for the five ΔRNR replicate lines evolved in 0.25 mg/mL (ΔRNR250_T30_L1-5) (**Supplementary Figure 3**). Note however that elongate cell morphology and clumping preclude use of OD measurements for accurate estimation of doubling time or cell counts^33^ (reflected in the large standard error relative to REL606 controls), so can only be used to give a general indication of growth. All lines failed to grow in the absence of dNS supplementation (**Figure 7E, Supplementary Figure 3**), indicating they remain reliant on media supplementation.

**Figure 7.**
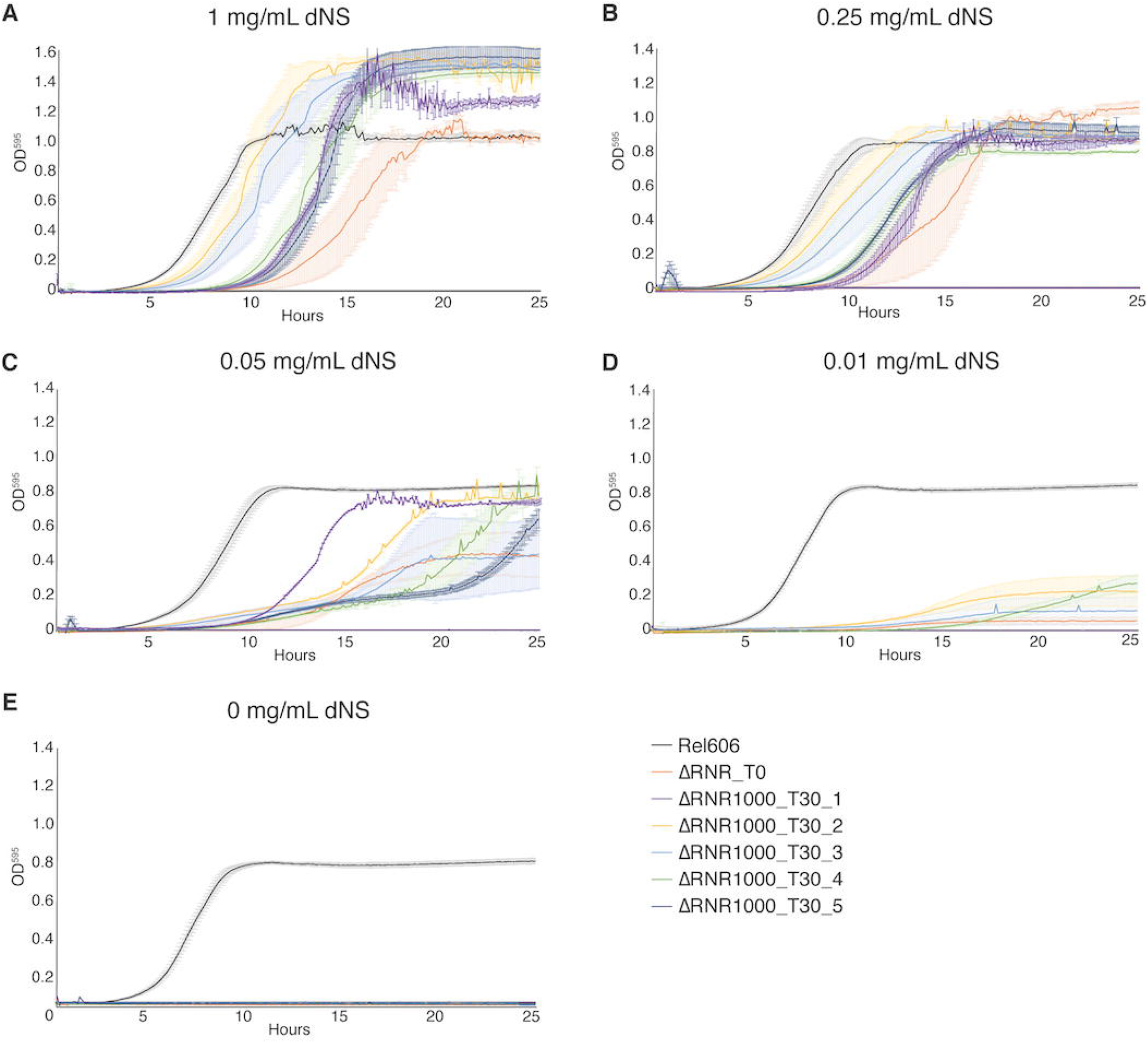
Growth characterisation of ΔRNR lines evolved at 1mg/mL dNS for 30 transfers. Growth was monitored for wild-type (REL606), ancestor (ΔRNR_T0) and evolved lines (ΔRNR1000_T30_L1-5, evolved in 1mg/mL dNS for 30 transfers). Growth experiments were performed in 1x MOPS media + 1% glucose with the addition of dNS at indicated concentrations. Growth was monitored for 25 hours. Curves show mean OD_595_, error bars show SEM, all experiments performed in triplicate. **A**. 1 mg/mL dNS, **B. 0**.25 mg/mL dNS, **C. 0.0**5 mg/mL dNS, **D. 0.0**1 mg/mL dNS, **E**. 0 mg/mL dNS.

### Evolution of ΔRNR at 0.25 mg/mL dNS results in A:T → G:C mutational skew

Following 30 transfers, we sequenced the genomes of ΔRNR_T0, our lines evolved in 0.25 mg/mL dNS (ΔRNR250_T30_L1-5) and 1 mg/mL dNS (ΔRNR1000_T30_L1-5), and each set of three wild-type control lines (REL606_1000_T30_L1-3 and REL606_250_T30_L1-3). The filamentous phenotype of our ΔRNR lines under restricted dNS supplement precluded accurate estimation of the number of generations per transfer, which in turn precluded reliable calculation of mutation rates. We therefore report total observed mutations (single nucleotide substitutions plus indels) for each experimental line (**Figure 8, Supplementary Table 5**). Our evolved knockout lines (ΔRNR250_T30_L1-5 & ΔRNR1000_T30_L1-5) all accumulated substantially more mutations than wild-type controls evolved at the same dNS concentration (**Figure 9A & Supplementary Table 5**). Our ΔRNR250_T30 lines accumulated the greatest numbers of mutations (range: 39-118 SNPs). ΔRNR1000_T30 lines did exhibit a small but significant increase in total SNPs relative to wild type (**Supplementary Figure 4 & Supplementary Table 5**; p=0.01, unpaired t-test). There was no significant difference in total mutations between wild type lines evolved in either 1 mg/ml or 0.25 mg/ml dNS (p=0.10; unpaired t-test)

**Figure 8.**
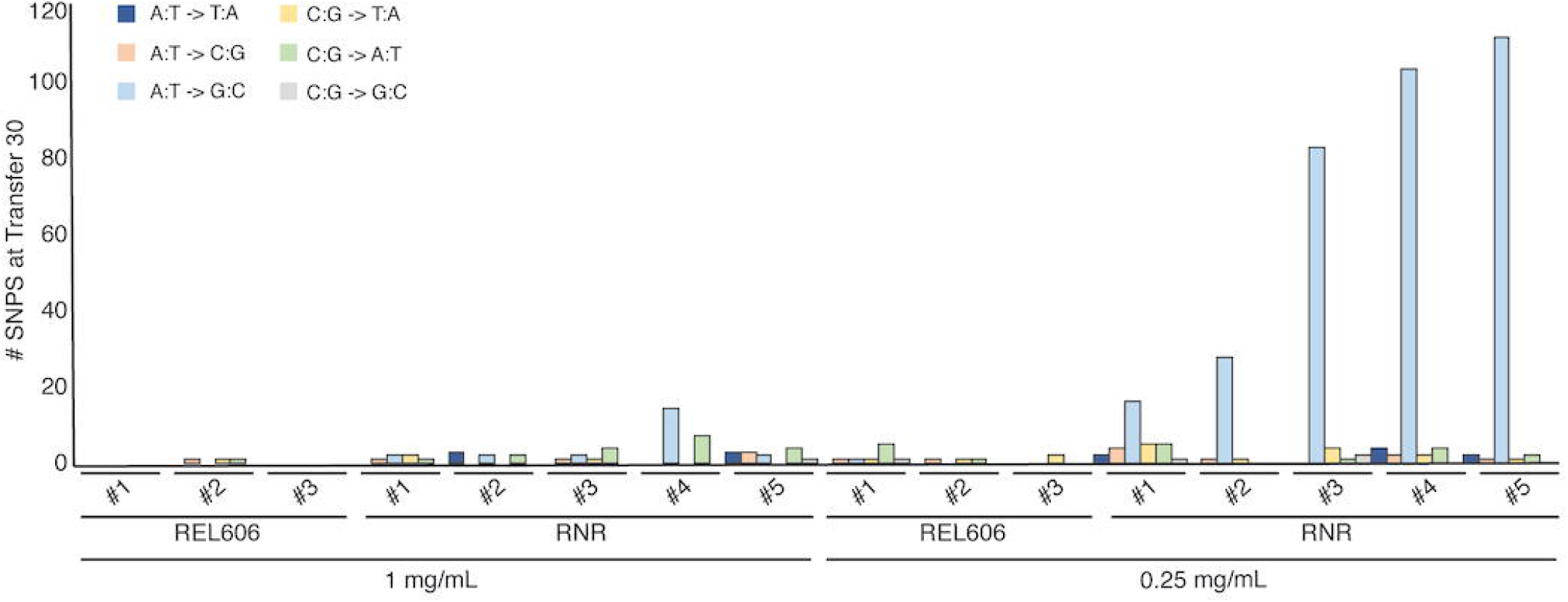
Mutations observed following experimental evolution of ΔRNR for 30 transfers. Total single nucleotide substitutions plus indels present in each line at T30. For evolution in 1mg/ml dNS, REL606 = control, RNR = ΔRNR1000_T30_L1-5. For evolution in 0.25mg/ml, REL 606 = control, RNR = ΔRNR250_T30_L1-5.

**Figure 9.**
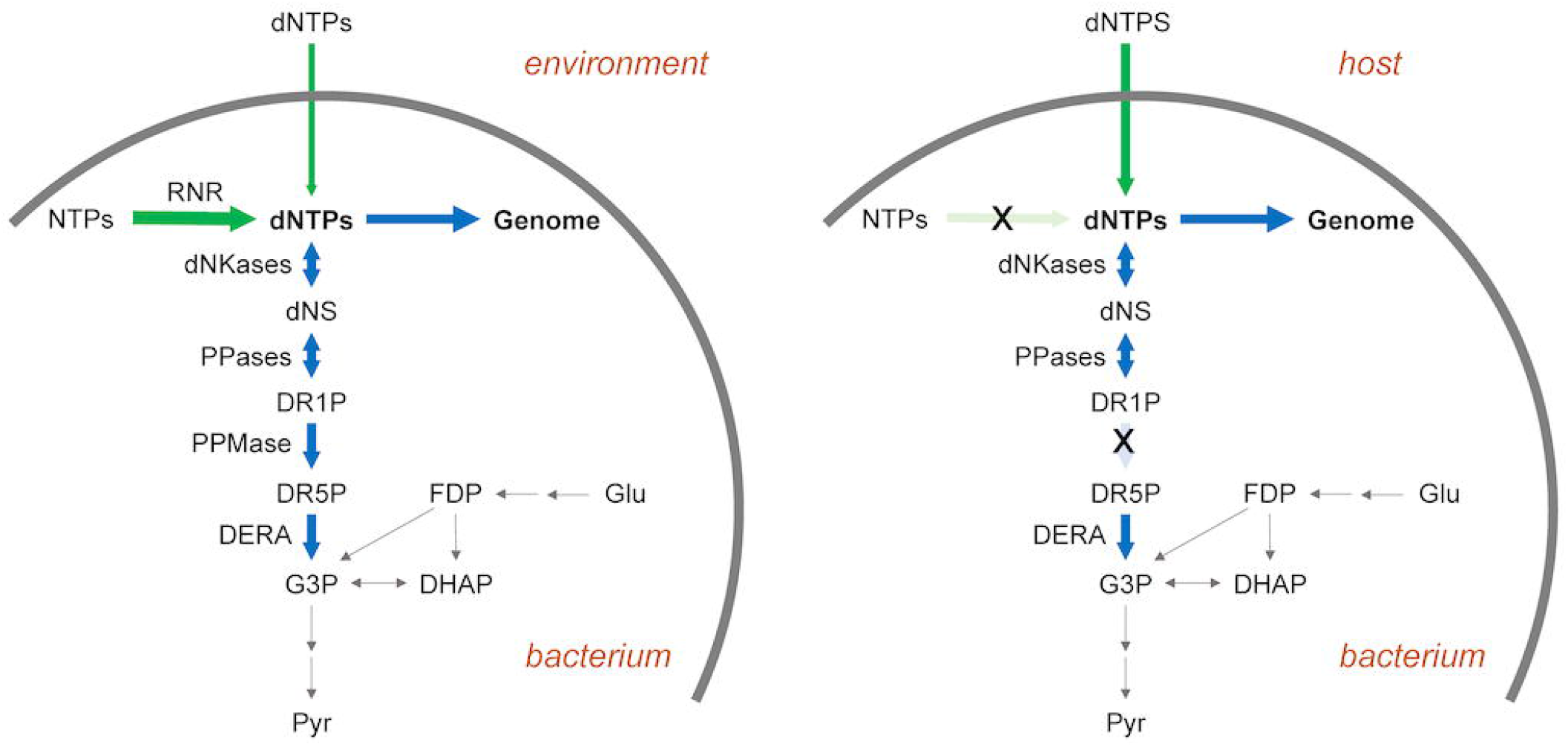
Impact of mutations on dNTP synthesis. **A**. dNTPs in free-living bacteria are primarily synthesised intracellularly via ribonucleotide reduction (RNR), with some uptake from the environment also possible in principle. dNTPs are required for genome synthesis, but may also be metabolised via the DERA pathway, the end product (G3P) of which feeds into glycolysis. **B**. In intracellular bacteria lacking RNR (crossed-out light green arrow), dNTPs must instead be derived from the host (green arrow). Loss of RNR leads to loss of phosphopentomutase (PPMase) activity (crossed-out light blue arrow), which prevents loss of deoxyribonucleosides to glycolysis. Abbreviations: RNR—ribonucleotide reductase; NTPs—nucleoside triphosphates; dNTPs—deoxyribonucleoside triphosphates; dNS— deoxyribonucleosides; DR1P—2-deoxyribose 1-phosphate; DR5P—2-deoxyribose 5-phosphate; G3P— D-glyceraldehyde 3-phosphate; DHAP—dihydroxyacetone phosphate; Glu—glucose; FDP—fructose 1,6-diphosphate; Pyr—pyruvate, dNKases— deoxynucleotide kinases; PPases— phosphorylases; DERA—deoxyriboaldolase.

It is well documented that ribonucleotide reductases keep deoxyribonucleotide pool sizes balanced through allosteric regulation^10^. We were therefore interested to see whether loss of ribonucleotide reduction resulted in mutational skew in *E. coli*. Our null expectation was that there would not be any detectable skew as our lines are supplemented with equal concentrations of all four deoxyribonucleosides. If there was skew, we expected this to be most evident in our ΔRNR250_T30 lines, as these accumulated the greatest number of mutations. Indeed, 88% of SNPs in our ΔRNR250_T30 lines are of one type: A:T**→**G:C (**Figure 8, Supplementary Figure 4**).

### ΔRNR lines lose cytidine deaminase

Analysis of our genome data from lines at transfer 30 revealed that one gene, *cdd*, which codes for cytidine deaminase, was mutated in 8 of 10 lines from transfer 30 (**Supplementary Tables 6 & 7**). Cytidine deaminase catalyses deamination of cytidine and deoxycytidine to uridine and deoxyuridine. Analysis of the data revealed that 7 of 8 lines carried an identical deletion (**Supplementary Tables 6 & 7; Supplementary Figure 5**), while one line possessed a SNP that was predicted to result in a truncation. As the deletions are identical, we suspected that this deletion may have occurred in the ancestral population, but was not fixed at the time we initiated our evolution experiment. These genome data were derived from colony isolates, meaning it was not possible to determine if this was the case. Thus, the significance of this mutation was unclear in the context of our evolution experiment. To determine whether *cdd* is lost in response to the experimental conditions, we screened our original ΔRNR isolates to establish if any carried intact *cdd*. We undertook PCR screening of glycerol stocks for two knockout lines (ΔRNR31, ΔRNR34). This revealed that the glycerol stock that we used to establish our evolution experiments (ΔRNR31) was a mixed population, with both intact and *cdd* deletion present (**Supplementary Figure 6**). Moreover, it appears that, after thirty transfers, all ten lines (ΔRNR_T30_1000_L1-5, ΔRNR_T30_250_L1-5) were polymorphic for the deleted locus. Together, this indicates the deletion is very likely to have occurred in the ancestral population, and that our lines were derived from a genetically heterogeneous population, despite the knockout lines being established from individual colonies following scarless deletion of *nrdAB* (Methods). Screening of ΔRNR34 glycerol stock however revealed no evidence of a deletion (**Supplementary Figure 6**). We therefore established a short evolution experiment with five independent ΔRNR34 lines, in 0.25 mg/mL dNS, mimicking the early stages of our evolution experiment with the ΔRNR31 isolate. After two transfers, we observed the emergence of a *cdd* deletion in several replicate lines (**Supplementary Figure 6**). In similar experiments not reported here and using these lines, we repeatedly see loss of the *cdd* locus under similar growth conditions (*data not shown*). Finally, that ΔRNR_T30_250_L5 carried an inactivating mutation in *cdd* but not the upstream locus (*yohk*) suggests that it is deletion of the *cdd* locus is selected under these conditions.

### ΔRNR lines evolved in 0.01 mg/ml dNS mutate the deoxyriboaldolase (DERA) salvage pathway

By transfer 30, one of our lines (ΔRNR_T30_250_L1) exhibited growth similar to wild-type when grown in 0.25 mg/ml dNS (**Supplementary Figure 3**). Using this line as progenitor, we initiated a second evolution experiment using ten parallel lines, grown in 0.01 mg/ml dNS (**Figure 5**). After ten transfers, we sequenced these lines (ΔRNR_T40_10_L1.1-1.10). All exhibited a marked accumulation of mutations, with L1.7 accumulating 220 SNPs (**Supplementary Figure 7**). As with the lines from our first 30 transfers, the majority of mutations were A:T**→**G:C substitutions.

Genome analysis of T40 lines revealed 648 mutations (**Supplementary Table 8**), of which 80 were found in more than one line. Of these 59 were present in two lines, 16 in three lines and 4 in four lines (**Supplementary Table 9**).

One gene, *deoB*, was mutated in eight lines (**Table 1**), and is particularly noteworthy in the context of our experiment in that it codes for phosphopentomutase. This enzyme catalyzes the transfer of a phosphate group between the C1 and C5 carbon atoms of deoxribose and is responsible for the commited step in deoxyribonucleotide salvage (**Figure 9**). Closer analysis revealed that several of the mutations to *deoB* were identical. It is possible that these mutations have occurred independently, but it may also be possible that these were the result of cross-contamination between lines during the evolution experiment. To assess this, we examined whether the affected lines carried a large number of shared mutations. Lines L1.1 and L1.8 had 47 and 157 unique mutations, with only the deoB mutation shared. For this pattern to be explained by cross-contamination, this mutation would have had to have been the very first mutation to appear, and cross-contamination would have had to have occurred before any other mutation occurred. With no evidence of *deoB* mutation in the progenitor line at transfer 30 (ΔRNR250_T30_L1), we consider it highly unlikely that cross-contamination between these lines has occurred. In the second case, L1.3 (53 SNPs), L1.5 (40 SNPs) and L1.10 (123 SNPs) share a common *deoB* mutation along with 13 other mutations. On these data, it is harder to rule out cross-contamination. Conservatively, it thus appears that *deoB* has independently mutated five or six times under the conditions of our experiment.

**Table 1.** Genome sequencing of ΔRNR10_T40 lines identifies multiple mutations to *deoB*.

| Mutation | Location in ORF | Effect | Lines impacted |
| --- | --- | --- | --- |
| Substitution | 1057 (A→C) | AA substitution (T353P) | L1<br>L8 |
| Deletion | 583 (G(1)→G(0)) | ORF Truncation<br>(frameshift) | L2 |
| Deletion | 926 (G(4)→G(3)) | ORF Truncation<br>(frameshift) | L3<br>L5<br>L10 |
| Substitution | 932 (A→C) | AA substitution (H311P) | L4 |
| Substitution | 1147 (A→C) | AA substitution (T383P) | L6 |

**Table 2.**
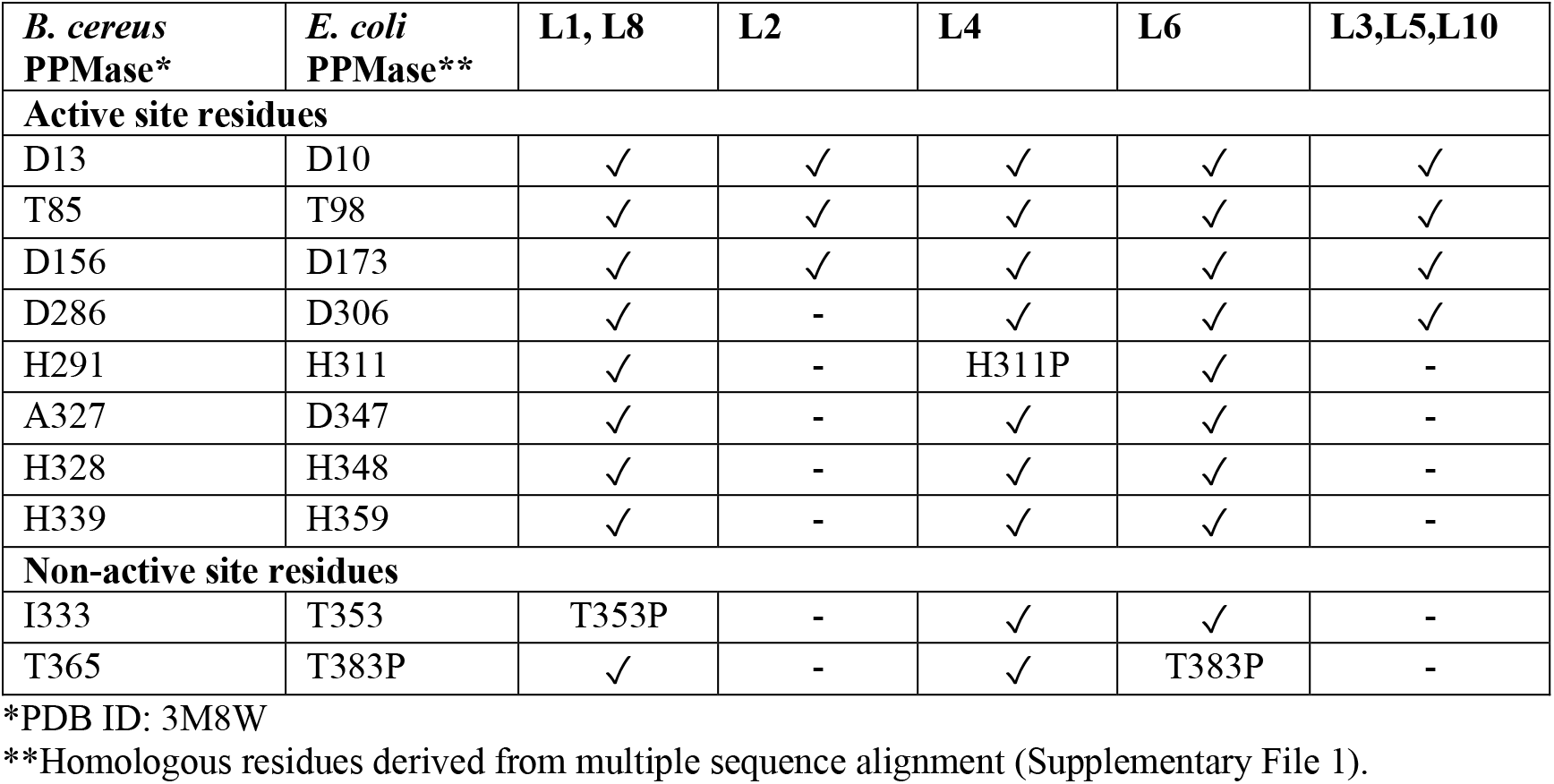
Predicted impact of observed *deoB* mutations on phosphopentomutase function.

| *B. cereus*<br>PPMase* | *E. coli*<br>PPMase** | L1, L8 | L2 | L4 | L6 | L3,L5,L10 |
| --- | --- | --- | --- | --- | --- | --- |
| **Active site residues** |  |  |  |  |  |  |
| D13 | D10 | ✓ | ✓ | ✓ | ✓ | ✓ |
| T85 | T98 | ✓ | ✓ | ✓ | ✓ | ✓ |
| D156 | D173 | ✓ | ✓ | ✓ | ✓ | ✓ |
| D286 | D306 | ✓ | - | ✓ | ✓ | ✓ |
| H291 | H311 | ✓ | - | H311P | ✓ | - |
| A327 | D347 | ✓ | - | ✓ | ✓ | - |
| H328 | H348 | ✓ | - | ✓ | ✓ | - |
| H339 | H359 | ✓ | - | ✓ | ✓ | - |
| **Non-active site residues** |  |  |  |  |  |  |
| I333 | T353 | T353P | - | ✓ | ✓ | - |
| T365 | T383P | ✓ | - | ✓ | T383P | - |

\*PDB ID: 3M8W
\*\*Homologous residues derived from multiple sequence alignment (Supplementary File 1).

To next sought to assess the impact of the mutations in *deoB*, on phosphopentomutase function. We first generated an alignment of 100 homologous phosphopentomutases, including from *E. coli* (REL606) and *Bacillus cereus* (**Supplementary File 1**). The latter was included as protein structure and active site residues have been characterised^28^. Using the alignment as a guide, we next assessed functional impact of observed *deoB* mutations by mapping these onto the structure of *B. cereus* phosphopentomutase (PDB ID: C3M8). Of the five unique mutations observed in our experiment (**Table 1**), three directly impact conserved active site residues, either through truncation or mutation of active site residues (**Table 2**). The remaining two mutations are not directly associated with the active site but result in amino acid substitutions not observed in any sequences in our alignment. We therefore conclude that these mutations to the *deoB* gene are all likely to impact phosphopentomutase function, with some expected to completely inactivate the enzyme.

## Discussion

While all life is dependent on ribonucleotide reduction for the synthesis of deoxyribonucleotides, a handful of obligate intracellular species are known to lack ribonucleotide reductase genes^2^ and presumably depend on their host for a source of dNTPs. In this work, we report the successful deletion of all ribonucleotide reductase operons from *E. coli* (**Figure 1**). The resulting line (ΔRNR) was sequenced to confirm genomic deletion of ribonucleotide reductase genes. Our ΔRNR knockout line grows on MOPS minimal media +1% glucose supplemented with deoxyribonucleosides (dNS) (**Figure 2**), but does not grow when provided with their constituent molecules (deoxyribose and the four bases A, G, C, T). Growth experiments reveal that, of the four dNSs, only deoxycytidine (dC) is essential for growth (**Figure 3**). Under limiting levels of dNS, our lines exhibit a filamentous phenotype that can be reversed by increasing dNS supplementation (**Figure 4**).

In order to understand how obligate intracellular bacteria adapted to the loss of ribonucleotide reduction, we established an evolution experiment where we serially passaged replicate lines in either 1 mg/mL or 0.25 mg/ml dNS. The resulting lines were sequenced after 30 transfers and one line evolved in 0.25 mg/ml dNS (ΔRNR_250_T30_1) was selected for a second round of evolution at 0.01 mg/ml dNS through a further ten transfers (**Figure 5**).

Genome sequencing revealed independent mutations to two genes across our replicate lines lines. One gene, *cdd*, carried mutations in 8/10 lines by transfer 30. The *cdd* locus codes for cytidine deaminase, which converts deoxycytidine to deoxyuridine, which in turn will be available for dTTP production. Our data reveal two independent mutations (a segmental deletion in 7 lines, and an inactivating SNP in the eighth). The segmental deletion deleted part of the *cdd* ORF and part of ORF of the upstream gene, *yohK*, which, together with *yohJ*, has been shown to code a 3-hydroxypropionate transporter^34^, and was postitionally identical across all seven lines. This deletion likely occurred prior to the establishment of our replicate lines, but appeared not to have gone to fixation. An independent ΔRNR line (ΔRNR34) lacked this deletion but rapidly lost *cdd* when passaged in the experimental growth media (**Supplementary Figure 6**). This, together with the independent truncation mutation in ΔRNR_T30_250_L5, which impacts *cdd* but not *yohK*, suggests there is selection against *cdd* but not necessarily *yohK*. Given the fact that our ΔRNR lines can grow with deoxycytidine (dC) as sole deoxyribonucleoside supplement but do not grow if dC is omitted from the deoxyribonucleoside mix, it may be that eliminating cytidine deamination prevents loss of this key deoxyribonucleoside through deamination. We have not assessed growth on dU, but one possibility is that deoxyuridine is less amenable as a substrate for production of all four dNTPs. One intriguing observation is that, despite the rapid loss of *cdd*, this loss of function appears not to have occurred across all replicates by transfer 30. This may be a consequence of the length of the evolution experiment. Comparing the genomes of species known to lack genes for ribonucleotide reduction^2^ reveals absence of *cdd* frequently coincides with this state. For instance, all confirmed members of the genus *Ureaplasma* lack both *nrd* genes and *cdd* (**Supplementary Text 2**), suggesting loss of cdd function is selectively advantageous. Establishing whether loss of one gene drives the loss of the other is not straightforward however. An analysis of strains of *Buchnera aphidicola* revealed that 48/74 strains retain ribonucleotide reduction, but 0/74 possess *cdd* (**Supplementary Text 2; Supplementary Table 14**), indicating that the latter can be lost in lines capable of synthesising their own deoxyribonucleotides. The picture is also not clear in *Borrelia*, with *cdd* found in species variously possess or lack ribonucleotide reduction (**Supplementary Text 2**). Deletion of *yohK* and *cdd* results in a small, but intact ORF (**Supplementary Figure 5**). While this seems unlikely to be functional, the creation of an ORF, particularly where the upstream gene codes for a transmembrane protein, may be consistent with the **‘**Car Trunk**’** hypothesis for the avoidance of mutations that would create a toxic protein^35^. The fact that deletion of these genes is reproducible under defined experimental conditions (**Supplementary Figure 6**) suggests that this region might be suitable for testing this hypothesis.

All lines at grown in 0.25 mg/ml dNS showed a marked mutational skew (A:T**→**G:C) that differs the expected G:C**→**A:T mutational skew expected for *E. coli*^*36*^, and more generally for bacteria^37^. Knockout of *cdd* does not appear linked to this skew; 4/5 lines grown at 1mg/ml dNS do not show an obvious A:T**→**G:C skew (**Figure 8**). For lines grown at 0.25 mg/ml dNS, two (L2, L3) carry intact *cdd* genes and both genomes show evidence of A:T**→**G:C mutational skew. Such skew has been previously shown to be associated with defective mismatch repair^36^. By transfer 30, we observe only two lines with SNPs in mismatch repair genes, with only one mutation being nonsynonymous (Supplementary Table 5). Thus, the skew we observe cannot be explained by widespread mutation to the mismatch repair pathway. The skew may instead be driven in part by the growth dependency; our data show that dC is the sole deoxyribonucleoside required for growth (**Figure 3D**), with the other dNSs unable to compensate for lack of dC (**Figure 3E**) but appearing to provide minor improvements to growth (**Figure 2B**). If these other dNSs are less readily taken up, the majority of intracellular dNTP production would derive from dC supplied in the media. If so, the dCTP pool would be higher than other dNTPs.

We also observed loss of function of *deoB* in our experiments (**Table 1**). This observation is interesting for several reasons. First, it indicates that loss of ribonucleotide reduction does not lead to compensatory deoxyribonucleotide synthesis via the DERA pathway, which has been speculated to be an ancestral route for dNTP synthesis, predating the advent of ribonucleotide reduction^20,21^. While the enzymes from this pathway can operate biosynthetically^16,17,38^, loss of phosphopentomutase function via mutation of *deoB* suggests that this pathway is not readily accessible for synthesis of deoxyribonucleotides *in vivo*, even though there would have been a strong selective advantage to such an alternative synthetic route during our experiment. We note however that both acetaldehyde toxicity (to both deoxyriboaldolase and to cells more generally) and low intracellular availability may preclude a simple switch to using the DERA pathway for dNTP production^17,21^.

Indeed, under the conditions of our experiment, where dNS supplementation is reduced by 25-fold, we observe strong selection to dispense with *deoB*. This suggests that not only is the reverse DERA pathway is inaccessible for biosynthesis, but salvage is selected against when dNTPs are scarce. In the second phase of our experiment, dNS supplementation is dropped from 0.25mg/ml to 0.01mg/ml, and it is under these conditions of severe dNS restriction that we observe parallel loss-of function mutations at the *deoB* locus (**Table 1**). We conclude that, when deoxyribonucleotides are in short supply, *deoB* loss-of-function ensures dNTPs are not lost to glycolysis via salvage (**Figure 9**). The biochemistry of the DERA pathway supports mutation of *deoB* as the most plausible target, since this step is responsible for interconversion of deoxyribose-5-phosphate and deoxyribose-1-phosphate. The latter is the substrate for phosphorylase-mediated addition of nucleobases, generating deoxyribonucleosides, whereas the former is the substrate for deoxyriboaldolase-mediated salvage. Thus, *deoB* loss-of-function mutations serve to prevent deoxyribose loss to central metabolism while still permitting phosphorylase-mediated deoxyribonucleoside resynthesis (**Figure 9**). Thus, while conversion of DERA to a synthetic pathway might be a better long-term solution to loss of ribonucleotide reduction, loss of function of this pathway is advantageous in the short term, precluding its cooption to dNTP synthesis. This interpretation is consistent with a recent experiment where mutation of dihydrofolate reductase led to *deoB* loss of function to prevent loss of deoxyribose-5-phosphate to glycolysis^39^.

Finally, analysis of species that lack *nrd* genes reveals frequent loss of *deoB*, suggesting that loss of ribonucleotide reduction might drive loss of *deoB*, as observed in our experiments. For members of *Ureaplasma*, the pattern is clear (**Supplementary Text 2**): all members of this genus completely lack genes for ribonucleotide reduction and show no evidence of coding for either *deoB* or *cdd*, the two most mutated genes in our experiments. For *Buchnera* aphidicola *cdd* appears to have been completely lost from all sequenced strains. Close to half of sequenced *Buchnera* strains show co-occurrence of ribonucleotide reduction and phosphopentomutase genes (48/74 strains), with 30% (22/74 strains) lacking genes for both functions (**Supplementary Table 14**), suggesting that loss of one function results in loss of the other. However, two strains appear to carry *deoB* while lacking *nrd* genes, and two further strains show the opposite pattern. If confirmed, this would indicate loss of one function may not always precipitate loss of the other.

We observe a very clear pattern in members of the *Borreliaceae*. The NCBI Taxonomy Browser^40^ lists two genera with genome data, *Borrelia* and *Borreliella* following a recent proposal^41-43^ to split the genus *Borrelia* in two. (though this has been contested^44-46^). Our analyses indicate that *Borrelia* (sensu Adeolu & Gupta^41^) code class Ib ribonucleotide reductases but lack *deoB* (**Supplementary Text 2**), while *Borrelliela* lack both *deoB* and RNR genes. This pattern could be taken to indicate that members of the *Borrelia* lost *deoB* but retained ribonucleotide reduction. However, phylogenetic analyses^47^ and location on linear plasmids^48^ indicate that the class Ib RNR genes (*nrdEF*) in *Borrelia* has been acquired via horizontal gene transfer, presumably following initial loss of ribonucleotide reduction in the *Borreliaceae*.

It is noteworthy that class Ib RNR, carried by members of the *Borrelia* (**Supplementary Text 2**), is manganese-dependent^49^, whereas class Ia is an iron-dependent enzyme^50^. Interestingly, the most well-studied member of the *Borreliaceae, Borreliella burgdorferi* (*sensu* Adeolu & Gupta) does not require iron^51^. It is tempting to speculate that initial loss of ribonucleotide reduction was driven by selection for iron independence, with subsequent reacquisition of ribonucleotide reduction favouring the manganese-dependent enzyme.

More generally, the switch to an intracellular existence may result in relaxed selection on ribonucleotide reduction if host-derived deoxyribonucleotides are sufficient to replace endogenous production. This appears to be an infrequent outcome as most intracellular bacteria retain ribonucleotide reduction. This rare event might be driven by additional pressures, as evolution of iron independence in *B. burgdorferi* suggests. Our experiments indicate that, in the absence of endogenous ribonucleotide reduction, disruption of the DERA pathway via mutations to *deoB* may serve to minimise loss of these building blocks when dNTP availability is limited. Given the necessity of genome replication, we conclude that life without ribonucleotide reduction drives the loss of dNTP catabolism via DERA, enabling building block reuse. It remains to be seen whether there are conditions wherein the reverse DERA pathway^21^ can operate synthetically *in vivo* as an alternative to ribonucleotide reduction.

## Author Contributions

Designed experiments: SDMA, NS, RJC, AMR, NH, DS, MH, KT, MT, JO, AMP.

Performed experiments: SDMA, NS, RJC, AMR, NH, DS, KD, MT.

Analysed data: SDMA, NS, AMR, NH, DS, MH, KT, MT, JO, AMP.

Wrote the paper: SDMA, AMP.

All authors read and approved final submitted version of the manuscript.

## Acknowledgments

We thank Auckland Genomics for technical support with genome sequencing, and the Faculty of Science Imaging Centre (University of Auckland) for image analysis.

