## Supplementary Materials for "Characterisation of an *Escherichia coli* line that completely lacks ribonucleotide reduction yields insights into the evolution of obligate intracellularity"

**Supplementary Text 1.** Knocking out ribonucleotide reduction in *E. coli*.
**Supplementary Text 2.** Comparative analysis of obligate intracellular bacterial genomes.

*Figures:*
**Supplementary Figure 1.** RT-PCR indicates that ribonucleotide reductase genes are not expressed in ΔRNR.
**Supplementary Figure 2.** RT-PCR confirms dAK expression in ΔRNR.
**Supplementary Figure 3.** Growth characterisation of ΔRNR lines evolved at 250 μg/mL dNS for 30 transfers.
**Supplementary Figure 4.** Mutations observed following experimental evolution of ΔRNR for 30 transfers.
**Supplementary Figure 5.** Schematic showing mutations detected in the *cdd* locus upon genome sequencing of all evolution lines at transfer 30.
**Supplementary Figure 6.** Genomic deletions of *cdd* in transfer 30 lines are also present in the ancestor but rapidly re-evolve under the same experimental conditions.
**Supplementary Figure 7.** Overview of mutational changes after experimental evolution for 10 transfers (Transfers 31-40).
*Tables:*
**Supplementary Table 1.** Strains
**Supplementary Table 2.** Primers
**Supplementary Table 3.** Mutations in ΔRNR ancestor (Arras_Supplementary_Table_3_Mutations_in_deltaRNR.xlsx)
**Supplementary Table 4.** ΔRNR cells are elongated relative to wild-type
**Supplementary Table 5.** T30 SNPs (Arras_Supplementary_Table_5_SNPs_at_T30.xlsx)
**Supplementary Table 6.** Mutations to the *cdd* gene in T30 lines
**Supplementary Table 7.** T30 parallel SNPs (Arras_Supplementary_Table_7_Parallel_SNPs_at_T30.xlsx)
**Supplementary Table 8.** T40 SNPs (Arras_Supplementary_Table_8_SNPs_at_T40.xlsx)
**Supplementary Table 9.** T40 parallel SNPs (Arras_Supplementary_Table_9_Parallel_SNPs_at_T40.xlsx)
**Supplementary Table 10.** Query sequences for blastp searches.
**Supplementary Table 11.** Blastp search results confirm absence of RNRs and PPM in species listed in Lundin et al. (2009)
**Supplementary Table 12.** *Candidatus* Ureaplasma intestinipullorum hits
**Supplementary Table 13.** *Candidatus* Borreliella tachyglossi hits
**Supplementary Table 14.** *Buchnera* comparative analysis (Arras_Supplementary_Table_14_Buchnera_comparative_analysis.xlsx)
*Data:*
**Appendix 1.** dAK gene from *Mycoplasma mycoides* (*mm-dAK*) codon-optimised for *E. coli*.
**Supplementary File 1.** Alignment of phosphopentomutases. (Arras_Supplementary_File1.pdf)

### Supplementary Text 1: Knocking out ribonucleotide reduction in *E. coli*.

***Stepwise generation of E. coli line lacking ribonucleotide reduction***

E. coli has three ribonucleotide reductase operons: *nrdAB* codes for class Ia ribonucleotide reductase, *nrdDG* codes for class III ribonucleotide reductase, and *nrdHIEF* codes for the class Ib enzyme. To create a line that lacks all of these enzymes, we used a scarless protocol[1] to delete operons successively, in the BL-21 *E. coli* line REL606 (see methods). All knockouts are detailed in **Supplementary Table 1.** As the class III enzyme only functions under anaerobic conditions, we knocked this out first, creating a *ΔnrdDG* line. We next deleted the *nrdHIEF* operon, yielding a double knockout line.

To knock out the *nrdAB* operon, we sought to identify conditions where *de novo* deoxyribonucleotide synthesis could be replaced by supplementation. We determined that deoxyribonucleoside (dNS) supplementation could replace de novo synthesis provided that we first introduced deoxyribonucleoside kinase (dNKase) activity to make dNSs available for dNTP synthesis. We therefore transformed our lines with a pBAD33 construct containing a dNKase gene from *Mycoplasma mycoides*[2] that had been previously shown to phosphorylate dA, dG and dC. As *E. coli* codes for thymidine kinase activity (*tdk*), we reasoned that expression of the *M. mycoides* dNKase (*Mm-dAK* — characterised as a deoxyribonucleoside adenosine kinase as it has highest affinity for this deoxyribonucleoside) should render the *nrdAB* operon non-essential and thus amenable to knockout. We cloned the *Mm-dAK* gene (codon-optimised for *E. coli* expression (GenScript; Appendix 1)) into pBAD33 under control of an IPTG-inducible *tac* promoter with *rrnB* terminator into our double knockout line (*ΔnrdDGΔnrdHIEF*). We used scarless deletion with lines grown in LB plus 0.5%w/v dA, dC, dG. We omitted dT as dTTP is synthesised via deamination of dCTP and dC to dUTP and dU respectively.

Under these conditions, we were able to successfully knock out the *nrdAB* locus. The resulting line, ΔRNR (Supplementary Table 1), was confirmed to lack all three operons coding for ribonucleotide reduction using PCR (see Supplementary Table 2 for primers), RT-PCR (Supplementary Figure 1) and genome sequencing. *Mm-dAK* expression was confirmed using RT-PCR. We observed *Mm-dAK* expression in our ΔRNR line with and without IPTG induction (Supplementary Figure 2). We therefore elected to perform evolution experiments without IPTG.

### Supplementary Text 2: Comparative analysis of obligate intracellular bacterial genomes.

***Obligate intracellular bacteria that lack ribonucleotide reduction***

Three species of bacteria were noted by Lundin et al.[3] as completely lacking ribonucleotide reduction:

*Borrelia burgdorferi*
*Ureaplasma urealyticum*
*Buchnera aphidicola str. Cc*

To confirm this, and assess whether other members of species or genus lack these, we performed a series of blast searches, using RNR protein sequences as query.

To establish whether species and genera also carry or lack homologues of phosphopentomutase (encoded by *deoB*) and cytidine deaminase (*cdd*), both of which were mutated in our evolution experiment, we also screened for these proteins. As the deletions of *cdd* involve loss of *yohK* in several instances, we also screened for presence/absence of this protein. Sequences used for blast searches are given in **Supplementary Table 10**, and results summarised in **Supplementary Table 11**.

***All but one Ureaplasma genomes lack ribonucleotide reduction coding potential.***

To establish whether the results from Table B hold for other members of the genus *Ureaplasma*, we expanded our search to all members of the genus for which there were genome assemblies. At the time of writing, there are 58 genome assemblies available for *Ureaplasma* (taxid:2129) in the NCBI genome database https://www.ncbi.nlm.nih.gov/genome/?term=txid2129[Organism:exp]

Only one genome assembly, *Candidatus* Ureaplasma intestinipullorum — a MAG from a chicken gut microbiome study[4] returned hits to RNR query sequences (**Supplementary Table 12**). Reciprocal blast (blastp, nr_protein) was used to confirm whether the hits recover the same protein family. No other *Ureaplasma* assemblies revealed evidence of RNR, PPMase, CDD or YohK coding potential.

These results suggest that, with the possible exception of *Candidatus* Ureaplasma intestinipullorum, no members of the genus *Ureaplasma* carry genes for RNR, PPM or CDD. These results align with the results from our experimental evolution study, where loss of ribonucleotide reduction resulted in subsequent loss of *deoB* and *cdd* genes.

***Most Borellia genomes code ribonucleotide reduction but none code for phosphopentomutase.***

To establish whether *B. burgdorferi* is representative of other *Borrelia* (taxid:138) in lacking ribonucleotide reduction, we screened 77 genome assemblies available for the genus in NCBI genome (https://www.ncbi.nlm.nih.gov/genome/?term=txid138[Organism:exp]).

Using the sequences from **Supplementary Table 10** as query, we detected 73 hits to NrdE and 67 to NrdF, indicating that the majority of *Borrelia* genome species carry a class Ib RNR. This discrepancy in number was not further investigated, but could be down to either gaps in assemblies or the fact that the NrdE protein is considerably larger than NrdF. No other significant hits to ribonucleotide reductase proteins was detected.

In our searches, we found that, in the NCBI taxonomy database, *Borrelia burgdorferi* has been renamed *Boreliella burgdorferi*, following a proposal from Adeolu & Gupta[5] to split *Borrelia* into two genera, *Borrelia* and *Borreliella*, though this reclassification is the subject of ongoing debate[6-10]. As NCBI Taxonomy lists both genera and, we expanded our search to all members of *Borreliella* (taxid: 64895, 17 species, 607 genome assemblies), and use the taxonomic distinction here so as to enable replication of our analyses.

Interestingly, only one member of the *Borelliella* returned any hits to RNR: *Candidatus* Borreliella tachyglossi appears to have a class Ib RNR (**Supplementary Table 13**). Reciprocal blast (blastp, nr_protein) hits multiple RNR Ib proteins from members of the *Borrelia* genus. Given that the placement of of *Candidatus* Borreliella tachyglossi is deep but appears unresolved, it may be that this is species helps to pinpoint the timing of loss of RNR from the clade/genus *Borelliella*, though it is also possible that this loss is convergent.

For CDD, search against *Borrelia* (taxid:138) returned 20 hits, all annotated as cytidine deaminases but these are approximately half the length of the query sequence. This is not unexpected as the *E. coli* protein is known to carry two cytidine deaminase domains and is a homodimer, whereas the shorter proteins from *Borrelia* are common among bacteria (and are homotetramers). Both forms are cytidine deaminases. For the same search against *Borreliella* (taxid:64895), 33 hits were returned, all annotated as cytidine deaminases.

For all genome assemblies from both genera, no significant hits to either PPM (*deoB*) or YohK were returned.

These data indicate that no members of either *Borrelia* or *Borreliella* carry *deoB*, with only the *Borreliella* apparently lacking ribonucleotide reduction.

***Multiple strains of Buchnera aphidicola lack both ribonucleotide reduction and phosphopentomutase.***

Lundin et al.[3] noted that only one specific strain of *Buchnera aphidicola* (from the cedar aphid *Cinara cedri*) lacked ribonucleotide reductase genes. At the time, this was one of the smallest genomes reported for a bacterium, and the smallest *Buchnera* genome, indicating it is heavily reduced[11]. As indicated in **Supplementary Table 11**, a search against *B. aphidicola Str. Cc* (taxid:372461) revealed no significant similarity for any of the proteins listed in table A, suggesting that all have been lost.

We next examined all members of *Buchnera aphidicola* (taxid:9). This revealed widespread presence of RNR Ia (NrdA, NrdB) and PPM, but no evidence of either CDD or YohK coding potential.

To determine the distribution of ribonucleotide reduction and phosphopentomutase, and whether other strains also lack both these processes, we screened for presence of sequences from **Supplementary Table 10** coded in available *B. aphidicola* strain genome assemblies in NCBI Genome. Of 74 assemblies, we found 22 strains that lack both processes, two strains that possess a *deoB* gene but lack genes for ribonucleotide reduction, and two with the opposite pattern (**Supplementary Table 14**).

### Supplementary Figures

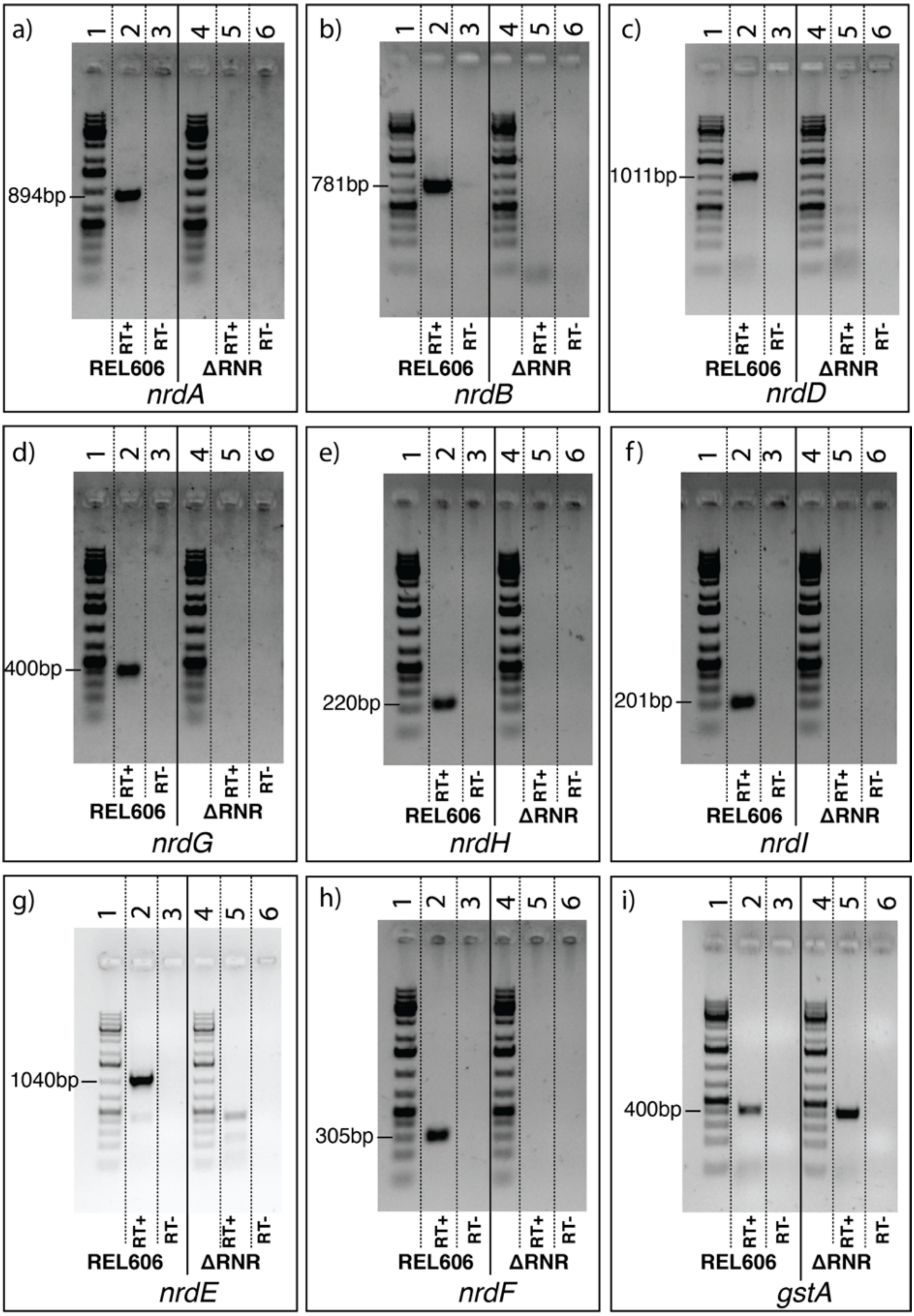

**Supplementary Figure 1. RT-PCR indicates that ribonucleotide reductase genes are not expressed in ΔRNR.** RT-PCR using primers for **A.** *nrdA,* **B.** *nrdB,* **C.** *nrdD,* **D.** *nrdG,* **E.** *nrdH,* **F.** *nrdI,* **G.** *nrdE* and **H.** *nrdF* on total RNA from ΔRNR or wild-type (REL606) indicate transcripts are not detected in ΔRNR, whereas expression was observed in REL606. **I.** Internal primers for the gstA housekeeping gene were used as a control for RT-PCR, confirming that the absence of amplification in ΔRNR was not due to the absence of RNA in the samples. For each primer pair the order of lanes is as follows: **lane 1**, 1kb+ ladder; **lane 2**, REL606 RT+ (RT enzyme added); **lane 3**, REL606 RT- (no RT enzyme added); **lane 4**, 1kb+ ladder; **lane 5**, ΔRNR RT+; **lane 6**, ΔRNR RT-.

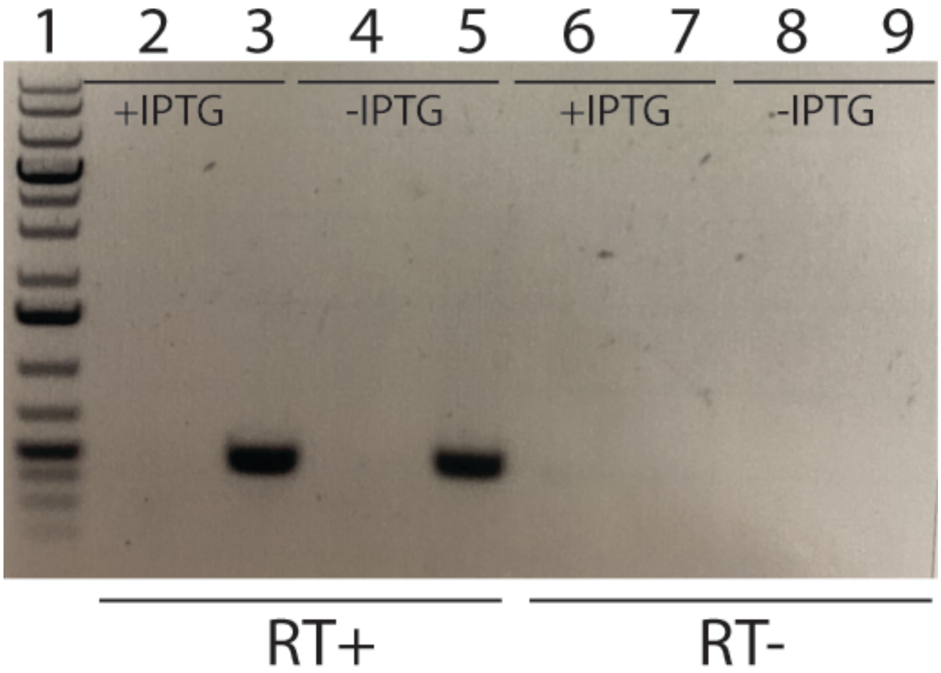


**Supplementary Figure *2*. RT-PCR indicates that Mm-dAK *is* expressed in ΔRNR.** RT-PCR using primers for codon-optimised *Mm-dAK* on total RNA from ΔRNR and wild-type (REL606) indicates Mm-dAK expression is observed in ΔRNR carrying the pBAD33::Mm-dAK construct, regardless of whether the cells are IPTG-induced (+IPTG) or uninduced (-IPTG) lines. Lane 1, 1kb+ ladder; Lanes 2-5 RT enzyme added (RT+): lane 2, WT +IPTG, Lane 3 ΔRNR +IPTG, Lane 4 WT -IPTG, Lane 5 ΔRNR -IPTG. Lanes 6-9 controls without RT enzyme (RT-): lane 6, WT +IPTG, Lane 7, ΔRNR +IPTG, lane 8 WT -IPTG, lane 9 ΔRNR -IPTG.

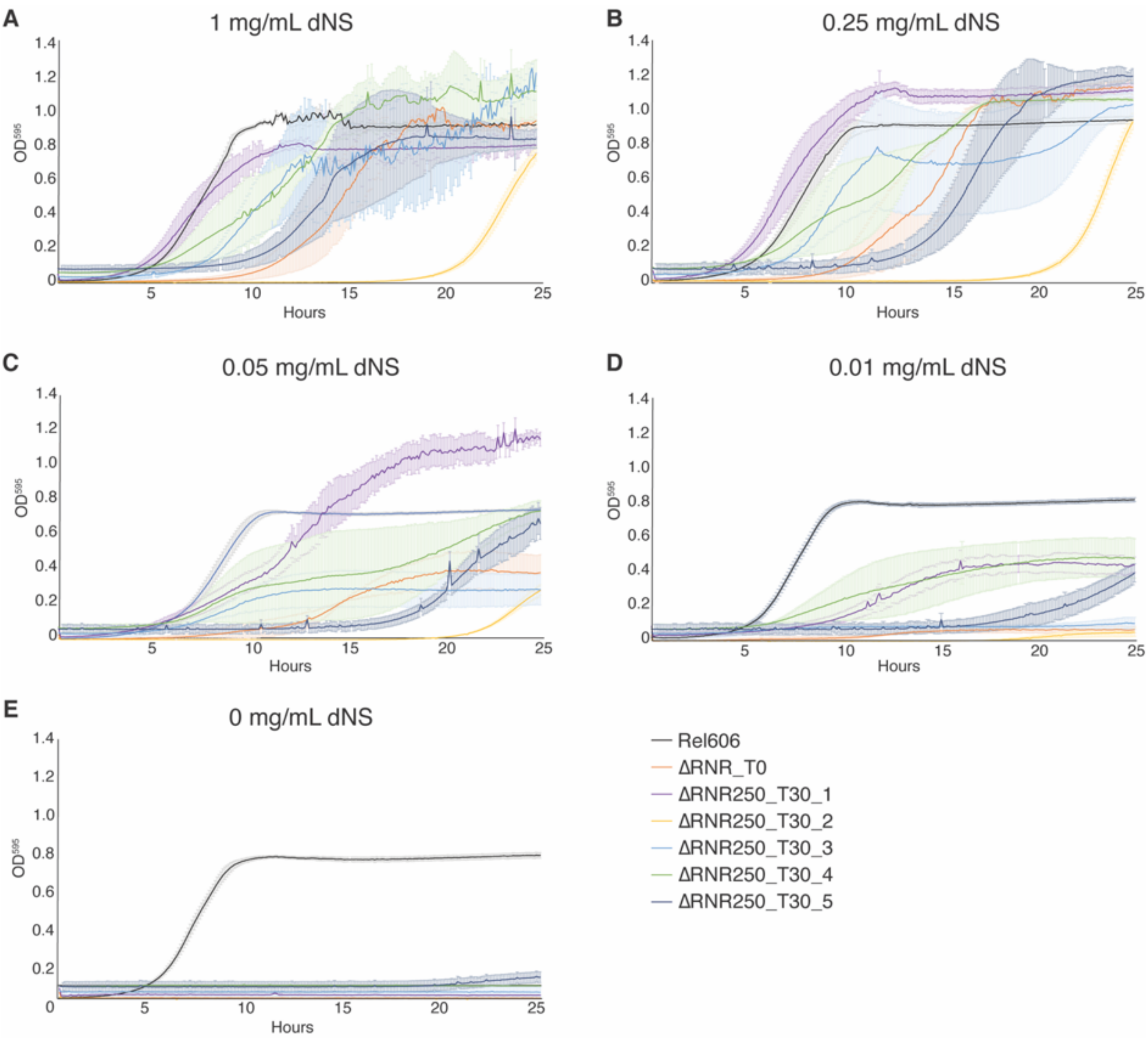


**Supplementary Figure 3. Growth characterisation of ΔRNR lines evolved at 250 µg/mL dNS for 30 transfers.** Growth was monitored for wild type (REL606), ancestor (ΔRNR_T0) and evolved lines (ΔRNR250_T30_L1-5, evolved in 250 µg/mL dNS for 30 transfers) . Growth experiments were performed in 1x MOPS media + 1% glucose with dNS supplementation, as indicated. Growth was monitored for 25 hours. Curves show mean $OD_{595}$, error bars show SEM, all experiments performed in triplicate. **A.** 1 mg/mL dNS, **B.** 0.25 mg/mL dNS, **C.** 0.05 mg/mL dNS, **D.** 0.01 mg/mL dNS, **E.** 0 mg/mL

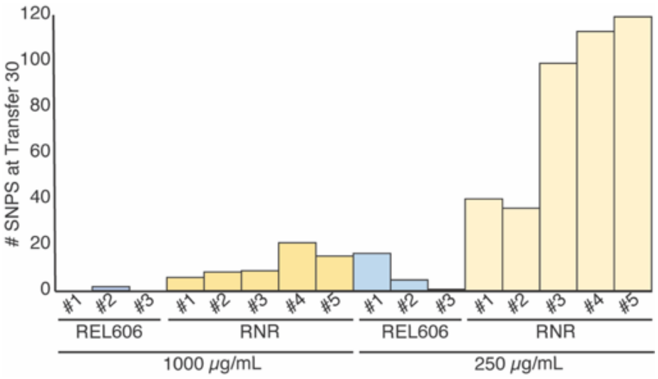


**Supplementary Figure 4. Mutations observed following experimental evolution of ΔRNR for 30 transfers.** Total single nucleotide substitutions plus indels present in each line at T30 (REL606 control lines; RNR = ΔRNR1000_T30_L1-5 (left) and ΔRNR250_T30_L1-5 (right).

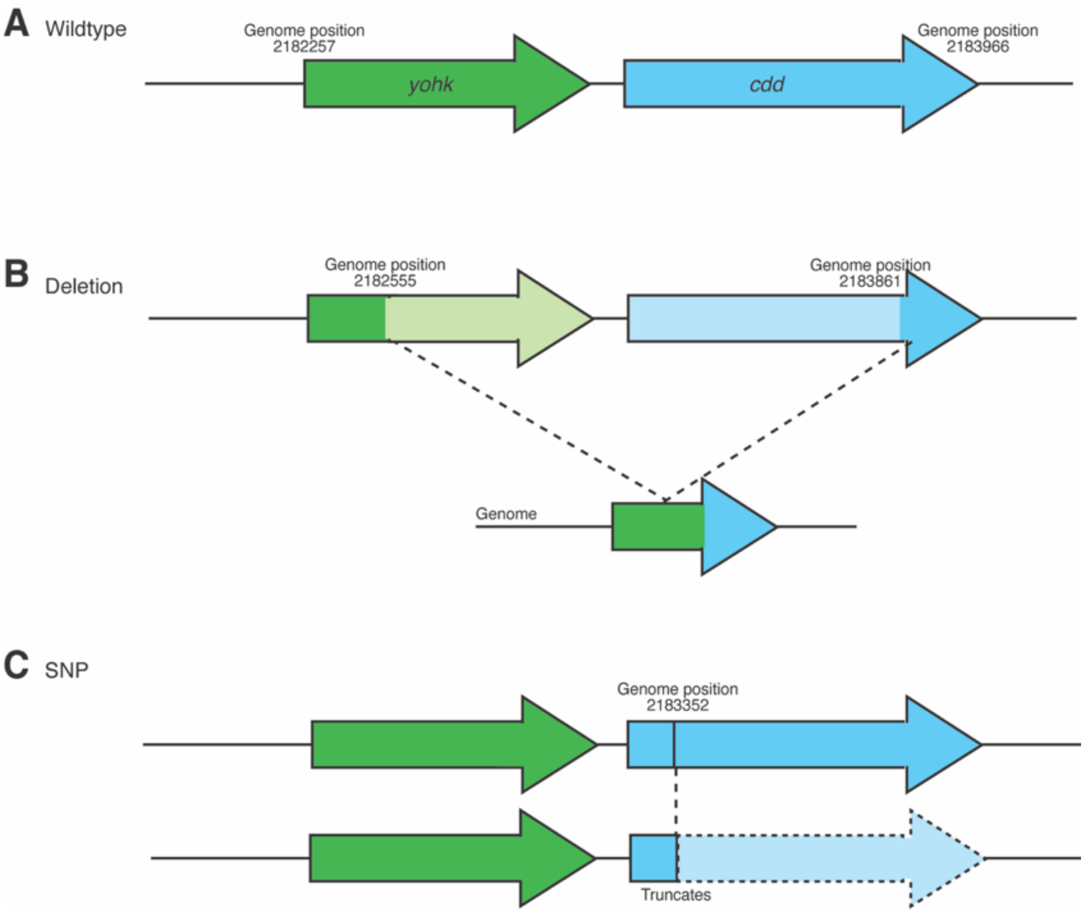


**Supplementary Figure 5. Schematic showing mutations detected in the *cdd* locus upon genome sequencing of all evolution lines at transfer 30** (ΔRNR_T30_1000_L1-5, ΔRNR_T30_250_L1-5). **A.** Wild-type genomic region. **B.** Deletion of *cdd* and upstream gene (*yohk*) occurs in 7 lines (Supplementary Tables 3 & 5). **C.** SNP in *cdd* locus of ΔRNR_T30_250_L5 results in ORF truncation at genomic position 2,183,352, resulting in a stop codon at codon position 91 in *cdd*.

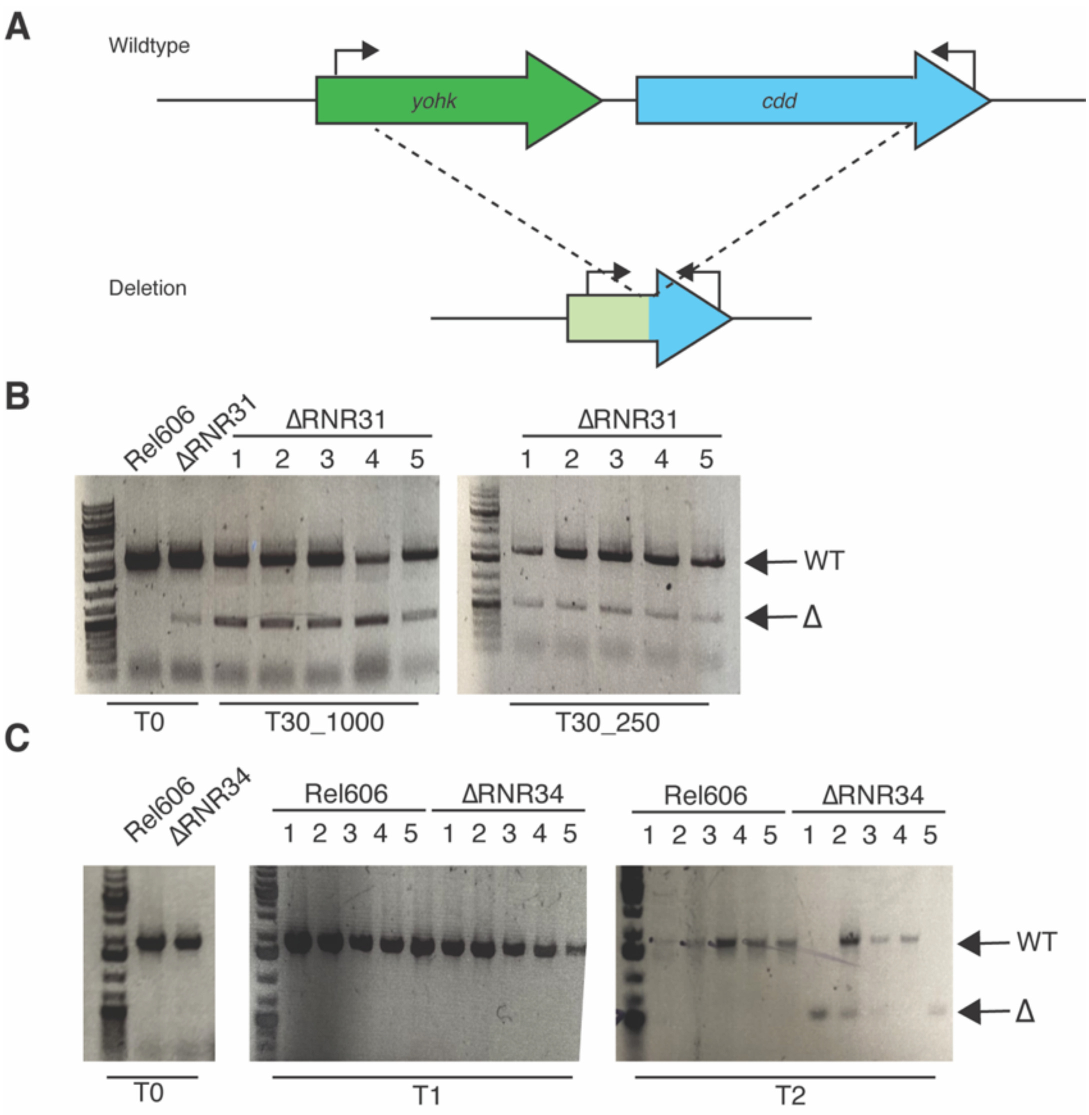


**Supplementary Figure 6. Genomic deletions of *cdd* in transfer 30 lines are also present in the ancestor but rapidly re-evolve under the same experimental conditions. A.** Schematic indicating approximate position of primers. See Supplementary Table 2 for primer sequences. **B.** PCR shows *cdd* deletion in the ΔRNR31 (ΔRNR_T0) population but not in wildtype. All ten lines at T30 carry deletion as well as WT variant, suggesting this deletion has not gone to fixation. **C.** The population derived from another ΔRNR isolate (ΔRNR34) lacks the *cdd* deletion (left panel). The *cdd* locus was monitored in the cell populations over two transfers. By the second transfer (T2), evidence of *cdd* deletion appeared in four lines (1-3, 5), indicating it is readily lost under the conditions of the evolution experiment. Lines grown in MOPS + 1% Glucose + 250µg/mL dNS. Note: experiment terminated after two transfers due to COVID-related lockdown.

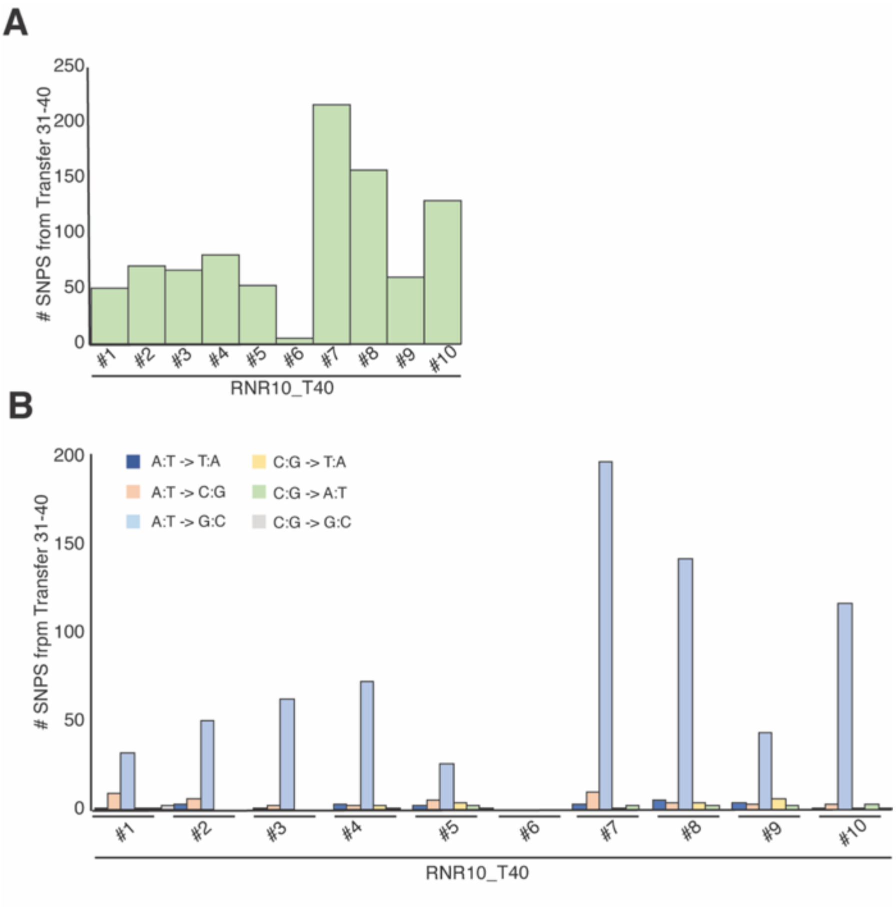


**Supplementary Figure 7. Overview of mutational changes after experimental evolution for 10 transfers (Transfers 31-40).** (For a description of the lines, see **Figure 5**). **A.** Total SNPs present in each line at T40. **B.** Types of mutations for all SNPs over the genome.

### Supplementary Tables

**Supplementary Table 1.** ***Escherichia coli*** **strains**

| Name | Description |
|---|---|
| REL606[12] | F-, *tsx467*(Am), *araA230*, *lon-*, *rpsL227*(strR), *hsdR-*, [*mal+*](*LamS*) |
| ΔRNR | REL606 Δ*nrdDG,* Δ*nrdHIEF,* Δ*nrdAB*, pBAD33::P*tac–mm-dAK–rrnB*t |
| ΔRNR1000_T30_1 | ΔRNR evolved in 1 mg/ml dNS at transfer 30. Line #1 |
| ΔRNR1000_T30_2 | ΔRNR evolved in 1 mg/ml dNS at transfer 30. Line #2 |
| ΔRNR1000_T30_3 | ΔRNR evolved in 1 mg/ml dNS at transfer 30. Line #3 |
| ΔRNR1000_T30_4 | ΔRNR evolved in 1 mg/ml dNS at transfer 30. Line #4 |
| ΔRNR1000_T30_5 | ΔRNR evolved in 1 mg/ml dNS at transfer 30. Line #5 |
| ΔRNR250_T30_1 | ΔRNR evolved in 250 μg/ml dNS at transfer 30. Line #1 |
| ΔRNR250_T30_2 | ΔRNR evolved in 250 μg/ml dNS at transfer 30. Line #2 |
| ΔRNR250_T30_3 | ΔRNR evolved in 250 μg/ml dNS at transfer 30. Line #3 |
| ΔRNR250_T30_4 | ΔRNR evolved in 250 μg/ml dNS at transfer 30. Line #4 |
| ΔRNR250_T30_5 | ΔRNR evolved in 250 μg/ml dNS at transfer 30. Line #5 |
| ΔRNR10_T40_1 | ΔRNR evolved in 10 μg/ml dNS at transfer 40. Line #1 |
| ΔRNR10_T40_2 | ΔRNR evolved in 10 μg/ml dNS at transfer 40. Line #2 |
| ΔRNR10_T40_3 | ΔRNR evolved in 10 μg/ml dNS at transfer 40. Line #3 |
| ΔRNR10_T40_4 | ΔRNR evolved in 10 μg/ml dNS at transfer 40. Line #4 |
| ΔRNR10_T40_5 | ΔRNR evolved in 10 μg/ml dNS at transfer 40. Line #5 |
| ΔRNR10_T40_6 | ΔRNR evolved in 10 μg/ml dNS at transfer 40. Line #6 |
| ΔRNR10_T40_7 | ΔRNR evolved in 10 μg/ml dNS at transfer 40. Line #7 |
| ΔRNR10_T40_8 | ΔRNR evolved in 10 μg/ml dNS at transfer 40. Line #8 |
| ΔRNR10_T40_9 | ΔRNR evolved in 10 μg/ml dNS at transfer 40. Line #9 |
| ΔRNR10_T40_10 | ΔRNR evolved in 10 μg/ml dNS at transfer 40. Line #10 |

**Supplementary Table 2. Primers**

| Name | Sequence |
|---|---|
| nrdA internal F | 5′ –CTATGAGAGCGCCCAGTTCC– 3′ |
| nrdA internal R | 5′ –GTACACAGCGCGATTTCACC– 3′ |
| nrdB internal F | 5′ –ATCAGACGCTGCTGGATTCC– 3′ |
| nrdB internal R | 5′ –AGCCAGGTGTTGATCCATGG– 3′ |
| nrdD internal F | 5′ –ATTACTCGCCGTTCTTCCCG– 3′ |
| nrdD internal R | 5′ –ACAGCGCGTTAATGGTTTCG– 3′ |
| nrdG internal F | 5′ –CTGCACCCTGTTTGTCTCCG– 3′ |
| nrdG internal R | 5′ –ATGCACCACCTGATTGCTGC– 3′ |
| nrdH internal F | 5′ –TGCGCATTACTATTTACACTCGTA– 3′ |
| nrdH internal R | 5′ –GGATGCAGACGGTTAATCATGT– 3′ |
| nrdI internal F | 5′ –CGGATTCAGGTAGACGAGCC– 3′ |
| nrdI internal R | 5′ –GCATTTCCGGGCAATCACAT– 3′ |
| nrdE internal F | 5′ –TGTGACCTTCAGTAGCCAGC– 3′ |
| nrdE internal R | 5′ –ACTCTGAGGCGCTGTTAACC– 3′ |
| nrdF internal F | 5′ –ATGCACTCACGCCTCATGAA– 3′ |
| nrdF internal R | 5′ –CGCGGTATTGGTCAGCTTTC– 3′ |
| gstA internal F | 5′ –CTTTGCCGTTAACCCTAAGGG– 3′ |
| gstA internal R | 5′ –GCTGCAATGTGCTCTAACCC– 3′ |
| nrdAB +800 | 5′ –CGATACTCCAGTCCTGCGTAATGC– 3′ |
| nrdAB -800 | 5′ –CGACCAACGATTGTCCGTGAGG– 3′ |
| nrdDG +800 | 5′ –GGCTTCTGGGTAGAGAATGGCG– 3′ |
| nrdDG -800 | 5′ –CCAGTAAATCCCGTGACAACAGCC– 3′ |
| nrdHIEF +800 | 5′ –GGCCTACCTTACCTGATTGCGC– 3′ |
| nrdHIEF -800 | 5′ –GCGTTCTTCGGCATTAATTCCGG– 3′ |
| cdd diagnostic F | 5′ –TGGTCATTACCGCTGACATTG– 3′ |
| cdd diagnostic R | 5′ –GCAGGAGACTCGATCTAAGGA– 3′ |

**Supplementary Table 4. ∆RNR cells are elongated relative to wild-type**

| Strain | Mean cell length* (μM) |
|---|---|
| REL606 | 1.33 |
| ∆RNR | 23.55 |

*Mean length (μM) of 100 cells each from REL606 and ∆RNR lines grown in 10 μg/mL dNS shows that ∆RNR cells are elongated relative to wild type (t-test, p=<0.000).

**Supplementary Table 6. Mutations to the *cdd* gene in T30 lines**

| Mutation | Genomic location | Effect | Lines impacted |
|---|---|---|---|
| Deletion | 2182555-3861 | ORF Deletion | ΔRNR1000_T30_L1-5, ΔRNR250_T30_L1,L4 |
| Substitution | 2183352 (G→T) | ORF Truncation | ΔRNR250_T30_L5* |

*A second nonsynonymous SNP was detected in *ccd* in this line (genomic position 2183737), but this is downstream of the truncation (Supplementary Table 7).

**Supplementary Table 10. Query sequences for blastp searches**

| Protein | Gene name | Species | Accession |
|---|---|---|---|
| NrdA (RNR Ia large subunit) | *nrdA* | *E. coli* | NP_416737.1 |
| NrdB (RNR Ia small subunit) | *nrdB* | *E. coli* | NP_416738.1 |
| NrdE (RNR Ib large subunit) | *nrdE* | *E. coli* | NP_417161.1 |
| NrdF (RNR Ib small subunit) | *nrdF* | *E. coli* | NP_417162.1 |
| NrdJ (RNR II) | *nrdJ* | *Caulobacter vibrioides* | YP_002517339.1 |
| NrdD (RNR III catalytic subunit) | *nrdD* | *E. coli* | NP_418659.1 |
| NrdG (RNR III activase) | *nrdG* | *E. coli* | NP_418658.1 |
| PPM (phosphopentomutase) | *deoB* | *E. coli* | P0A6K6 |
| CDD (cytidine deaminase) | *cdd* | *E. coli* | P0ABF6 |
| YohK Inner membrane protein | *yohK* | *E. coli* | P0AD19 |

**Supplementary Table 11. Blastp search results confirm absence of RNRs and PPM in species listed in Lundin et al. (2009)[3]***

| Taxonomy ID | Species | RNRs | CDD | PPM | YohK |
|---|---|---|---|---|---|
| 139 | *Borrelia burgdorferi* | NSS | WP_044002316.1 | NSS | NSS |
| 2130 | *Ureaplasma urealyticum* | NSS | NSS | NSS | NSS |
| 372461 | *Buchnera aphidicola str. Cc (Cinara cedri)* | NSS | NSS | NSS | NSS |

*Query sequence accessions given in Supplementary Table 10. All blastp searches performed against nr_protein database, restricted by taxonomy ID; NSS: No Significant Similarity found.

**Supplementary Table 12. *Candidatus* Ureaplasma intestinipullorum hits***

| Protein | Hit (accession#)** | Reciprocal blast | Species | Comments |
|---|---|---|---|---|
| NrdA | MBU3830637.1 | MBD5445710.1 | *Mycoplasma sp.* | Reciprocal blast hits NrdA |
| NrdB | MBU3830638.1 | WP_004025021.1 | *Malacoplasma iowae* | Reciprocal blast hits NrdB |
| NrdE | NSS | | | |
| NrdF | MBU3830638.1 | | | Note same sequence retrieved as for NrdB search. NrdB and NrdF are homologous, lower bit score and reciprocal blast result indicates this is an NrdB sequence, not NrdF. |
| NrdD | NSS | | | |
| NrdG | NSS | | | |
| NrdJ | NSS | | | |
| CDD | NSS | | | |
| PPM | NSS | | | |
| YohK | NSS | | | |

*Searches using query sequences listed in Supplementary Table 10.
**NSS: No significant similarity

**Supplementary Table 13. *Candidatus* Borreliella tachyglossi hits***

| Protein | Hit (accession#)** | Reciprocal blast | Species | Comments |
|---|---|---|---|---|
| NrdA | WP_108729659.1 | | | Note same sequence retrieved as for NrdE search. NrdA and NrdE are homologous, lower bit score and reciprocal blast result indicates this is an NrdE sequence, not NrdA. |
| NrdB | NSS | | | |
| NrdE | WP_108729659.1 | WP_025420044.1 | *Borrelia anserina* | Reciprocal blast hits NrdE |
| NrdF | WP_108729658.1 | WP_120104792.1 | *Borrelia turcica* | Reciprocal blast hits NrdF |
| NrdD | NSS | | | |
| NrdG | NSS | | | |
| NrdJ | NSS | | | |
| CDD | WP_159076631.1 | WP_120104392.1 | *Borrelia turcica* | CDD |
| PPM | NSS | | | |
| YohK | NSS | | | |

*Searches performed using query sequences listed in Supplementary Table 10.
**NSS: No significant similarity

**Appendix 1.** dAK gene from *Mycoplasma mycoides* (*mm-dAK*) codon-optimised for *E. coli* (GenScript).

```
>mm-dAK ORF
ATGAAAATTGCGATCTTCGGCACCGTTGGCGCGGGTAAAAGCACCATCAGCGCGGAAATCAGCAAAAAACT
GGGCTATGAAATCTTCAAAGAGCCGGTGGAGGAAAACCCGTACTTCGAACAGTACTATAAAGACCTGAAGA
AAACCGTTTTCAAGATGCAAATCTATATGCTGACCGCGCGTAGCAAGCAGCTGAAACAAGCGAAGAACCTG
GAGAACATCATTTTCGACCGTACCCTGCTGGAAGATCCGATTTTTATGAAAGTGAACTACGACCTGAACAA
CGTTGACCAGACCGATTACAACACCTACATCGATTTCTACAACAACGTGGTTCTGGAGAACCTGAAAATTC
CGGAAAACAAGCTGAGCTTTGACATCGTGATTTACCTGCGTGTTAGCACCAAGACCGCGATCAGCCGTATT
AAGAAACGTGGTCGTAGCGAGGAACTGCTGATCGGCGAGGAATACTGGGAGACCCTGAACAAAAACTACGA
GGAGTTCTATAAGCAAAACGTGTATGATTTTCCGTTCTTTGTGGTTGACGCGGAACTGGATGTGAAGACCC
AGATTGAACTGATCATGAATAAGCTGAATAGCATTAAGAACCCGAACTAA
```
