## Supplementary File 1 for "Characterisation of an *Escherichia coli* line that completely lacks ribonucleotide reduction yields insights into the evolution of obligate intracellularity"

Supplementary File 1: Alignment of phosphopentomutases.

```
Consensus                         MNMMKRAFIMVLDSFGIGAAADADXF----------GDVGADTLGHIAEACAEGEADN-G-RKGPLKLPNLSRLGLGKAA   68
deoB_Bcereus_WP_098782964.1       MNKYKRIFLVVMDSVGIGEAPDAEQF----------GDLGSDTIGHIAE----------HMNGLQMPNMVKLGLGNIR   58
deoB_Rel606                       ---MKRAFIMVLDSFGIGATEDAERF----------GDVGADTLGHIAEACAKGEADN-G-RKGPLNLPNLTRLGLAKAH   65
sp|Q89A57|DEOB_BUCBP              ---MKRAFLIVLDSFGIGATKDAHRF----------GDVGANTFGNIAKFCFLNKANNYG-RKGLLYIPHLLSLGLAKVA   66
sp|Q8K936|DEOB_BUCAP              ---MKRVFLMILDSLGIGSSVDACKF----------NDSGADTLGHIAEKCFFNEANVQ--RRGSLYIPNLIKLGIVKSY   65
sp|B8D866|DEOB_BUCAT              ---MKRVFLIVLDSFGIGSSPDADKF----------NDVGSNTFGHIVEKCFLGEANV-G-RKGVLCIPNLVKLGIINAA   65
sp|P57607|DEOB_BUCAI              ---MKRVFLIVLDSFGIGSSPDADKF----------NDVGSNTFGHIVEKCFLGEANV-G-RKGVLCIPNLVKLGIINAA   65
sp|B8D9W4|DEOB_BUCA5              ---MKRVFLIVLDSFGIGSSPDADKF----------NDVGSNTFGHIVEKCFLGEANV-G-RKGVLCIPNLVKLGIINAA   65
sp|Q483R0|DEOB_COLP3              ---MARAIIIVIDSLGIGYSPDAVDF----------GDVGANTFANLARAYYEETGKE-------IFLPNLSALGMIKAC   60
sp|Q3ICU9|DEOB_PSET1              ---MARAIILMADSLGIGAAPDADKF----------GDIGANTLAHLLKAYYDETGTA-------LSLPNLSKLGLIDAC   60
sp|Q5QXT9|DEOB_IDILO              ---MSRAIVLVLDSFGIGSAPDADKF----------GDEGADTFGHIAEYRYQ--------HGQPLQLPNLTRLGLAEAH   59
sp|Q17YP1|DEOB_HELAH              --MQKRVVVLLLDSFGIGASEDAKDF----------GDLGANTLGNIAKACFNNLADSND-RNGALKLPNLESLGLGLSA   67
sp|Q9ZK37|DEOB_HELPJ              --MQKRVVILLLDSFGIGASEDAKDF----------GDLGANTLGNIAKACFNNLADSND-RNGVLKLPNLEGLGLGLSA   67
sp|B6JN18|DEOB_HELP2              --MQKRVVILLLDSFGIGASEDAKDF----------GDLGANTLGNIAKACFNNLADSND-RSGALKLPYLEGLGLGLSA   67
sp|P56195|DEOB_HELPY              --MQKRVVILLLDSFGIGASEDAKDF----------GDLGANTLGNIAKACFNNLADSND-RSGALKLPYLESLGLGLSA   67
sp|B5Z8H3|DEOB_HELPG              --MQKRVVVLLLDSFGIGASEDAKDF----------GDFGANTLGNIAKACFNNLANSND-RNGALKLPYLESLGLGLSA   67
sp|B2UUU2|DEOB_HELPS              --MQKRVVILLLDSFGIGASEDARDF----------GDLGANTLGNIAKACFNNLADSND-RNGALKLPYLESLGLGLSA   67
sp|Q1CS87|DEOB_HELPH              --MQKRVVILLLDSFGIGASEDAKDF----------GDLGANTLGNIAKACFNNLADSND-RKGALKLPYLESLGLGLSA   67
sp|A4YPV1|DEOB_BRASO              ----MRALIVVMDSVGIGGAPDAAGY----------GDEGADTVGHIAEACAAGQADSDQ-RQGPLQLPHLAALGLGEAC   65
sp|B8IJW7|DEOB_METNO              ---MSRALLIVLDSVGIGGAPDAASY----------GDEGADTVGHIAASCARGAGDRAGLRQGPLRLPHLAALGLGLAA   67
sp|B0UPC1|DEOB_METS4              ---MTRALVIVLDSVGIGGAPDGAAY----------GDGGADTLGHIAEACAEGRGDRPGLRAGPLRLPHLAGLGLGLAA   67
sp|B1LYJ9|DEOB_METRJ              ---MARALIIVLDSVGVGGAPDADRY----------GDTGSDTLGHIAERCAEGRGDRAGLRAGPLRLPHLAELGLGLAA   67
sp|B1ZIG8|DEOB_METPB              ---MARALLIVLDSVGIGGAPDADRY----------GDAGSDTVGHIAEACAAGRGDRPGLRAGPLRLPNLAALGLGLAC   67
sp|A9W427|DEOB_METEP              ---MARALLIVLDSVGIGGAPDADRY----------GDAGSDTVGHIAEACAAGRGDRPGLRAGPLHLPNLAALGLGLAC   67
sp|B7KXI3|DEOB_METC4              ---MARALLIVLDSVGIGGAPDADRY----------GDAGSDTVGHIAEACAAGRGDRPGLRAGPLRMPNLTALGLGLAC   67
sp|C3MBH5|DEOB_SINFN              ---MARVFLFVLDSFGIGGAPDAEAF----------GDLGADTLGHIAEFCASGAGDRAGLREGPLQLPNMSALGLVHAA   67
sp|Q92T47|DEOB_RHIME              ---MARAFLFVLDSFGIGNAPDAEAF----------GDLGADTLGHIAEFCAAGAADRAGLREGPLHLPNMSALGLMHAA   67
sp|A6UET6|DEOB_SINMW              ---MARAFLFVLDSFGIGNAPDAGAF----------GDLGADTLGHIAEFCAAGAADRAGLREGPLNLPNMSALGLMHAA   67
sp|B9JYP7|DEOB_AGRVS              ---MARAFLFVLDSFGVGGAPDAPAY----------GDDGADTLGHIAEFCAAGAGDRDGLRAGPLMLPNLSALGLLQIA   67
sp|Q8UJ04|DEOB_AGRFC              ---MARAFLLVLDSFGVGGAPDAERY----------GDLGANTLGHIAEFCAAGAADRAGLRAGPLKLPNMCSLGLLEIA   67
sp|B9J6V9|DEOB_AGRRK              ---MARAFLFVLDSFGVGGSPDATSY----------GDEGADTLGHIAEFCAAGAADREGLRSGPLSLPIMSGLGLLEIA   67
sp|Q2KDR4|DEOB_RHIEC              ---MARAFLFVLDSFGVGGAPDAAAY----------GDEGADTLGHIAEFCAAGAGDRAGLRSGPLSLPNMSELGLMHIA   67
sp|B5ZXI4|DEOB_RHILW              ---MARAFLFVLDSFGVGGAPDAAAY----------GDEGADTLGHIAEFCAAGAGDRAGLRDGPLSLPNLSELGLMQIA   67
sp|Q1MMV6|DEOB_RHIL3              ---MARAFLFVLDSFGVGGAPDAAAY----------GDEGADTLGHIAEFCAAGAADRAGLREGPLSLPNMSELGLMQIA   67
sp|Q98BG5|DEOB_RHILO              ---MARAFLFVLDSFGIGGAADAERY----------GDAGANTLAHIAEACAEGRADRERLRQGPLFVPHMASLGLGKAA   67
sp|Q11AV9|DEOB_CHESB              ---MARAFLFVLDSFGIGHAPDAARF----------GDEGANTFGHIAGACAAGRADRAGLRTGPLDLPHMQSLGLNAAA   67
sp|A5IGR8|DEOB_LEGPC              --MTGRVCVLVMDSFGIGASLDAARY----------GDVGANTLVHIYEACKRGECDIDGVRKGSLMLPNLASKGLYHAA   68
sp|Q5X7B2|DEOB_LEGPA              --MTGRVCVLVMDSFGIGASLDAARY----------GDAGANTLVHIYEACKRGECDIDGVRKGPLMLPNLAGKGLYHAA   68
sp|Q5WYR0|DEOB_LEGPL              --MTGRICILVMDSFGIGASLDAARY----------GDAGANTLVHIYEACKRGECDIEGARKGPLMLPNLAGKGLYHAA   68
sp|Q5ZXU2|DEOB_LEGPH              --MTGRVCVLVMDSFGIGASLDAARY----------GDAGANTLVHIYEACKRGECDIEGVRKGPLMLPNLAGKGLYHAA   68
sp|B2KBT7|DEOB_ELUMP              ---MKRVIILMMDSFGIGGAADAAKF----------GDEGANTLASVAK----------LNNGLKIPNLISLGLVKAA   55
sp|Q2SHN3|DEOB_HAHCH              ---MSRAILIVLDSFGIGATADAQRF----------GDAGANTLLHIAEACAAGACDSQG-RTGPLRLPNLNRLGLGRAG   66
sp|Q1QZC2|DEOB_CHRSD              ---MTRAILLVLDSFGIGNAPDAAAF----------GDAGADTLGHIAHHRAAS-----G-R--PLALPHLASLGLYHAH   59
sp|A1SYK3|DEOB_PSYIN              ---MKRTVILLLDSFGVGGAADADKFIGSLADGSQYDDMGSNTLGHIAEQCAKGNANE-G-RHGPLTIPNLNKLGFGRAC   75
sp|A6VM02|DEOB_ACTSZ              ---MKRVLIMMLDSFGIGGAEDADKF----------GDKGANTLGHIAS----------HQPSLNLPHLESLGLGLAA   55
sp|C4LAY9|DEOB_TOLAT              ---MKRTIILMLDSLGIGASADAVRF----------GDEGANTLGHIAQACVTGNADN-G-RKGPLTLPNLTRLGLAHAC   65
sp|A4SRU2|DEOB_AERS4              ---MKRTFILMMDSFGIGAAADADKF----------GDVGANTLGHIAKACAAGDIE---GRSALNLPNLNKLGLGHAG   63
sp|A0KPE2|DEOB_AERHH              ---MKRTFILMMDSFGIGAAADADKF----------GDVGANTLGHIAKACAAGEIE---GRGALNLPNLSKLGLGHAG   63
sp|Q7NRT1|DEOB_CHRVO              ---MKRAIILILDSLGIGAAADAPKF----------GDAGSNTLGHIAMEAAAGRADI-G-RSGPLKLPNLSRMGLGHAC   65
sp|Q086F8|DEOB_SHEFN              ---MKRTFILMLDSFGVGAATDADKF----------GDVGSDTFGHIAQACADGKADI-G-RQGPLKLPNLARFGLGHAG   65
sp|Q12QG1|DEOB_SHEDO              ---MKRTFILMLDSFGIGAASDADKF----------GDIGADTFGNIAKACAEGKANI-G-RDGSLKLPNLAKLGLGHAG   65
sp|A1S476|DEOB_SHEAM              ---MKRTIILMLDSFGIGAADDADKF----------GDVGSDTFGHIAQMCAEGKANN-G-REGELKLPNLSRLGLAHAA   65
sp|Q8EHK2|DEOB_SHEON              ---MKRTVIMMLDSFGVGAAGDAAKF----------GDVGSDTFGHIAKACAEGKADT-G-RKGPLALPNLARLGLAHAA   65
sp|A0KU09|DEOB_SHESA              ---MKRTVIMMLDSFGVGAAGDAAKF----------GDLGSDTFGHIAKACAEGEADI-G-REGPLTLPNLARLGLAHAA   65
sp|Q0HLE8|DEOB_SHESM              ---MKRTVIMMLDSFGVGAAGDAAKF----------GDLGSDTFGHIAKACAEGKADI-G-REGPLTLPNLARLGLAHAA   65
sp|Q0HXQ2|DEOB_SHESR              ---MKRTVIMMLDSFGVGAAGDAAKF----------GDLGSDTFGHIAKACAEGKADI-G-REGPLTLPNLARLGLAHAA   65
sp|A1RH89|DEOB_SHESW              ---MKRTIIMMLDSFGVGASADAASF----------GDVGSDTFGHIAKACAEGKADI-G-REGPLKLPNLARLGLGHAA   65
sp|A4Y9A6|DEOB_SHEPC              ---MKRTIIMMLDSFGVGASADAASF----------GDVGSDTFGHIAKACAEGKADI-G-REGPLKLPNLARLGLGHAA   65
sp|A6WRB6|DEOB_SHEB8              ---MKRTIIMMLDSFGVGASADAASF----------GDVGSDTFGHIAKACAEGKADI-G-REGPLKLPNLARLGLGHAA   65
sp|A9KZ78|DEOB_SHEB9 (2)          ---MKRTIIMMLDSFGVGASADAASF----------GDVGSDTFGHIAKACAEGKADI-G-REGPLKLPNLARLGLGHAA   65
sp|B8E6P6|DEOB_SHEB2              ---MKRTIIMMLDSFGVGASADAASF----------GDVGSDTFGHIAKACAEGKADI-G-REGPLKLPNLARLGLGHAA   65
sp|B0TQ89|DEOB_SHEHH              ---MKRTIIMMLDSFGIGEAHDAHAF----------GDVGSNTFGNIAKACAEGKAND-G-REGPLKLPNLARLGLGHAS   65
sp|A8H726|DEOB_SHEPA              ---MKRTIIMMLDSFGVGAAHDAEAF----------GDVGSNTFGHIAKACAEGKAND-G-REGPLKLPNLARLGLGHAS   65
sp|A8FYQ7|DEOB_SHESH              ---MKRTFIMMLDSFGIGAAADAEAF----------GDTGSDTFGNIAKACAEGKANE-G-REGPLKLPNLARLGLAQAS   65
sp|A3QGT1|DEOB_SHELP              ---MKRTVIMMLDSFGVGAAKDAEAF----------GDTGSNTFGHIAKACAEGKAND-G-REGPLKLPNLAKLGLALAA   65
sp|B1KRP6|DEOB_SHEWM              ---MKRTIIMMLDSFGVGAATDAESF----------GDVGSDTFGSIAKACAEGRADI-G-REGPLKLPNLSKLGLALAA   65
sp|Q6LUH2|DEOB_PHOPR              ---MKRSIILVLDSFGIGATEDAVDF----------GDVGSNTMGHIAQACARGEADN-GDRSGPLHLPNLNKLGLGKAC   66
sp|B6ENG5|DEOB_ALISL              ---MKRAIILVLDSFGIGAAGDAEKF----------GDVGSDTMGHIAEQCDKGLADN-GNRKGPLTLPNLSKLGLAMAG   66
sp|B5FAA0|DEOB_ALIFM              ---MKRAIILVLDSFGIGAAGDADKF----------GDVGSDTMGHIAEQCDKGLADN-GNRKGSLTLPNLSKLGLAMAG   66
sp|Q5E7J5|DEOB_ALIF1              ---MKRAIILVLDSFGIGAAGDADKF----------GDVGSDTMGHIAEQCDKGLADN-GNRKGPLTLPNLSKLGLAMAG   66
sp|C3LQC0|DEOB_VIBCM (2)          ---MKRAFILVLDSFGIGATADAQAF----------GDVGSDTLGHIADQCAQGLADNAE-RKGALQLPNLSKLGLAMAH   66
sp|Q9KPL9|DEOB_VIBCH              ---MKRAFILVLDSFGIGATADAQAF----------GDVGSDTLGHIADQCAQGLADNAE-RKGALQLPNLSKLGLAMAH   66
sp|A7MUW4|DEOB_VIBCB              ---MKRAFILVLDSFGIGAAADAEQF----------GDVGSDTLGHIADQCEQGLANNDK-REGALRLPNLSKLGLAMAH   66
sp|Q7MI40|DEOB_VIBVY (2)          ---MKRAFILVLDSFGIGATADAKEF----------GDVGSDTLGHIADQCEQGLANNDQ-RQGALRLPNLSKLGLAMAH   66
sp|Q87M24|DEOB_VIBPA              ---MKRAFILVLDSFGIGATADAKEF----------GDVGSDTLGHIADQCEKGLADNDK-RQGALRLPNLSKLGLAMAH   66
sp|Q2NW04|DEOB_SODGM              ---MQRVFIMVLDSFGIGAAEDADKF----------GDRGSDTLGHIAERCERGDANH-G-RKGPLKLPNLTRLGLAKAA   65
sp|B2VH52|DEOB_ERWT9              ---MKRAFIMVLDSFGIGSSKDAEKF----------GDKGSDTLGHIAEACFRGEADK-G-RKGPLHLPNLTALGLGKAA   65
sp|A7MGA8|DEOB_CROS8              ---MKRAFIMVLDSFGIGATEDAERF----------GDAGSDTLGHIAEACAKGEADI-G-RKGPLHLPNLTKLGLAKAH   65
sp|A4W6A0|DEOB_ENT38              ---MKRAFIMVLDSFGIGATEDADRF----------GDVGSDTMGHIAEACAKGEADI-G-RQGPLNLPNLTRLGLVKAH   65
sp|Q327L3|DEOB_SHIDS              ---MKRAFIMVLDSFGIGATEDAERF----------GDVGADTLGHIAEACAKGKADN-G-RKGPLNLPNLTRLGLAKAH   65
sp|Q1R260|DEOB_ECOUT (3)          ---MKRAFIMVLDSFGIGATEDAERF----------GDVGADTLGHIAEACAKGEADH-G-RKGPLNLPNLTRLGLAKAH   65
sp|B7UR11|DEOB_ECO27              ---MKRAFIMVLDSFGIGATEDAERF----------GDVGADTLGHIAEACAKGEADN-G-RKGPLNLPNLTRLGLAKAH   65
sp|Q3YU10|DEOB_SHISS              ---MKRAFIMVLDSFGIGATEDAERF----------GDVGADTLGHIAEACAKGEADN-G-RKGPLNLPNLTRLGLAKAH   65
sp|Q0SX28|DEOB_SHIF8 (22)         ---MKRAFIMVLDSFGIGATEDAERF----------GDVGADTLGHIAEACAKGEADN-G-RKGPLNLPNLTRLGLAKAH   65
sp|B5Y275|DEOB_KLEP3              ---MKRAFIMVLDSFGIGATEDADRF----------GDVGADTMGHIAEACAKGEADN-G-RKGPLNLPNLTRLGLVKAH   65
sp|A9MRA5|DEOB_SALAR              ---MKRAFIMVLDSFGIGATEDADRF----------GDVGSDTLGHIAEACAKGEADN-G-RKGPLNLPNLTRLGLVKAH   65
sp|B5BAJ9|DEOB_SALPK (2)          ---MKRAFIMVLDSFGIGATEDADRF----------GDVGSDTLGHIAEACAKGEADN-G-RKGPLNLPNLTRLGLVKAY   65
sp|C0Q7M5|DEOB_SALPC (2)          ---MKRAFIMVLDSFGIGATEDADRF----------GDVGSDTLGHIAEACANGEADN-G-RKGPLNLPNLTRLGLVKAH   65
sp|B5R9V1|DEOB_SALG2              ---MKRAFIMVLDSFGIGATADADRF----------GDVGSDTLGHIAEACAKGEADN-G-RKGPLNLPNLTRLGLVKAH   65
sp|B5FTC7|DEOB_SALDC              ---MKRAFIMVLDSFGIGATEDADRF----------GDVGSDTLGHIAEACAKGEADN-G-RKGPLNLPNLTRLGLVKAH   65
sp|P63924|DEOB_SALTI (8)          ---MKRAFIMVLDSFGIGATEDADRF----------GDVGSDTLGHIAEACAKGEADN-G-RKGPLNLPNLTRLGLVKAH   65
sp|C5BHJ4|DEOB_EDWI9              ---MKRAFIMVLDSFGIGEAKDAKSF----------GDEGADTLGHIARACARGEADI-G-RQGPLHLPNLSRLGLGKAA   65
sp|Q6D990|DEOB_PECAS              ---MKRVYIMVLDSFGIGSSADAERF----------GDVGSDTLGHIAQACAAGTADK-G-RSGSLHLPNLSRLGLGKAA   65
sp|C6DKL9|DEOB_PECCP              ---MKRAYIMVLDSFGIGSSADAERF----------GDAGSDTLGHIAQACAAGTADK-G-RSGPLHLPNLSRLGLGKAA   65
sp|B4EWA2|DEOB_PROMH              ---MKRVHIMVLDSFGIGAAGDAEKF----------GDQGADTLGHIAQAFAEGKADKDG-RKGPLHLPNLCRLGLGKAA   66
sp|Q7N931|DEOB_PHOLL              ---MKRTFIMVLDSFGIGASKDAEKF----------GDQGSNTLGHIAEACARGDADI-G-RKGPLYLPNLSRLGLGKAT   65
sp|A8G9H8|DEOB_SERP5              ---MKRTFIMVLDSFGIGASEDAAKF----------GDQGSDTLGHIAEVCARGEANV-G-RQGPLTLPNLSRLGLGKAA   65
sp|A1JJ99|DEOB_YERE8              ---MKRTFIMVLDSFGIGASADANKF----------GDEGADTLGHIAEACARGEADI-G-RSGPLTLPNLSRLGLGKAA   65
sp|A9R047|DEOB_YERPG              ---MKRTFIMVLDSFGIGASADAKKF----------GDEGADTLGHIAEACARGEANV-G-RSGPLTLPNLSRLGLGKAA   65
sp|A4TQJ1|DEOB_YERPP (4)          ---MKRTFIMVLDSFGIGASADAKKF----------GDEGADTLGHIAEACARGEANV-G-RSGPLTLPNLSRLGLGKAA   65
sp|B1JL35|DEOB_YERPY (4)          ---MKRTFIMVLDSFGIGASADAKKF----------GDEGADTLGHIAEACARGEANV-G-RSGPLTLPNLSRLGLGKAA   65
```

```
Consensus                         EESTG-RFP-AGLD--DNAEVIGAY-GHASELSSGKDTPSGHWEIAGVPVLFDWGYFSDKENSFPKELLDKIVARAXLPG  143
deoB_Bcereus_WP_098782964.1       EM--------KGIS--KVEKPLGYY-TKMQEKSTGKDTMTGHWEIMGLYIDTPFQVFPE---GFPKELLDELEEKTG-RK  123
deoB_Rel606                       EGSTG-FIP-AGMD--GNAEVIGAY-AWAHEMSSGKDTPSGHWEIAGVPVLFEWGYFSDHENSFPQELLDKLVERANLPG  140
sp|Q89A57|DEOB_BUCBP              IASSG-QDL-LGIQ--DTNNVIGSY-AYSNEISSGKDTSSGHWEIAGAPVLFDWDYFSSSLNSVPKYLIQDIVDQCNLSG  141
sp|Q8K936|DEOB_BUCAP              QSSTG-KYL-LGFNQKEDANIIASY-GYASELSSGKDTTSGHWEIAGAPFLKDWYYFKEKDNSFPSSLINKIITKLKLPG  142
sp|B8D866|DEOB_BUCAT              KESTG-QYP-LGFN--YSSNVIASY-GFASEISSGKDTTSGHWEIAGVPVLDDWCYFKEKQNSFPESLLEKIIRRSELTG  140
sp|P57607|DEOB_BUCAI              KESTG-QYP-LGFN--YSSNVIASY-GFASEISSGKDTTSGHWEIAGVPVLDDWYYFKEKQNSFPESLLEKIIIRSELTG  140
sp|B8D9W4|DEOB_BUCA5              KESTG-QYP-LGFN--YSSNVIASY-GFASEISSGKDTTSGHWEIAGVPVLDDWYYFKEKQNSFPESLLEKIIRRSELTG  140
sp|Q483R0|DEOB_COLP3              EQASNQTFP--YQ--GQEPSKGAY-GFAQEISTGKDTPSGHWEMAGVPVLFDWGYFTDKNSSFPTSLIDDINRETGFDG  134
sp|Q3ICU9|DEOB_PSET1              E-AAG-KEP-CLVS--NRSASQGAW-GYAKELSSGKDTPSGHWEMAGVPVLFDWGYFPKTQPSFPQEFIDELIARTGIPG  134
sp|Q5QXT9|DEOB_IDILO              KEATG-TYA-KGFH---EVETHGAY-ACAAELSSGKDTPSGHWELMGVPVHFEWGYFQDKENSFPNELLAQILKACELDG  133
sp|Q17YP1|DEOB_HELAH              LKATN-ELP-LGFD--SHPNLIGAY-AYAQELSSAKDTISGHWEMMGAPILFEWGYFKDKNNSFPKEILDEIMHKTKIKG  142
sp|Q9ZK37|DEOB_HELPJ              LKAAN-ELP-LGFQ--LKPNLIGAY-AYAKELSSAKDTISGHWEMMGAPVLFEWGYFKDKNDSFPKEILDEIMHKTKIKG  142
sp|B6JN18|DEOB_HELP2              LEAAN-ELP-LGFQ--PQPNLIGAY-AYAQELSSAKDTISGHWEMMGAPVLFEWGYFKDKNDSFPKEILDEIARKTKIKG  142
sp|P56195|DEOB_HELPY              LKAAN-ELP-LGFQ--SQPNLIGAY-AYAQELSSAKDTISGHWEMMGAPVLFEWGYFKDKTHSFPKEILDEIVRKTKIKG  142
sp|B5Z8H3|DEOB_HELPG              LKATN-ELP-LGFE--SKPNLIGAY-AYAKELSSAKDTISGHWEMMGAPVLFEWGYFKDKTHSFPKEILDEIMHKTKIKG  142
sp|B2UUU2|DEOB_HELPS              LKATN-ELP-LGFE--SKPNLIGAY-AYAQELSSAKDTISGHWEMMGAPVLFEWGYFKDKNNSFPKEILDEIMRKTKIKG  142
sp|Q1CS87|DEOB_HELPH              LKATN-ELP-LGFE--SKPNLIGAY-AYAKELSSAKDTISGHWEMMGAPVLFEWGYFKDKNNSFPKEILDEIMRKTKIKG  142
sp|A4YPV1|DEOB_BRASO              EHATG-RRP-PGLQ--QKAGAGAAF-GCASERSRGKDTPSGHWEIAGVPVPFEWGYFPKTVPAFPPDLIDQLCASAGLTG  140
sp|B8IJW7|DEOB_METNO              EAATG-RMP-PGLA--PAGSVTGAW-GHAVETARGKDTVSGHWEIAGVPVDFDWGRFPATEPSFPPELTQALIAEAGLPG  142
sp|B0UPC1|DEOB_METS4              REASG-RMP-PGLA--PDGPVAGAW-GHAVETARGKDTVSGHWEIAGAPVDFDWGRFPATRPSFPPDLTRALIEEGNLPG  142
sp|B1LYJ9|DEOB_METRJ              AGASG-RIP-PNLT--PVGPPRGVY-GHAVETAAGKDTPSGHWEIAGLPLPEPWGHFPDTRPAFPPELTRALVAEGDLPG  142
sp|B1ZIG8|DEOB_METPB              EGATG-RVP-PGLA--PGEPLRALW-GHAVETAAGKDTPSGHWEIAGVPVRAPWGHFPDTHPAFPAELTAALIARAGLPG  142
sp|A9W427|DEOB_METEP              EGATG-RVP-PGLA--PEGPVRALW-GHAVETAAGKDTPSGHWEIAGVPVREAWGHFPDTQPAFPAELTAALVARAGLPG  142
sp|B7KXI3|DEOB_METC4              EGATG-RVP-PGLA--PDGPVRALW-GHAVETAAGKDTPSGHWEIAGVPVREAWGHFPDTQPAFPAELTAALIERAGLPG  142
sp|C3MBH5|DEOB_SINFN              KLATG-RLP-AGMP--LPERVYGVY-GAASEVSRGKDTPSGHWEIAGTPVRFDWGYFPAEGDAFPPQLVEAICREGEVPG  142
sp|Q92T47|DEOB_RHIME              RLATG-RLP-AGMA--LPERVYGIY-GAASEVSRGKDTPSGHWEIAGTPVTFDWGYFPAEGDAFPPELVEAICREGDVPG  142
sp|A6UET6|DEOB_SINMW              RLATG-RLP-AGMA--LPERVYGVY-GAASEVSRGKDTPSGHWEIAGTPVTFDWGYFPADGDAFPPELVEAICREGDVPG  142
sp|B9JYP7|DEOB_AGRVS              SLASG-SLP-AGMA--LPERVFGLY-GAANEISRGKDTPSGHWEIAGTPVMFDWGYFPQEGDAFPADLVADICKRADLPG  142
sp|Q8UJ04|DEOB_AGRFC              RQASG-DIP-AGME--PPERIFGLH-GSASEISKGKDTPSGHWEIAGTPVTFDWGYFPTEGDAFSPELVEAICRQADIPG  142
sp|B9J6V9|DEOB_AGRRK              KAATG-RYP-LGMP--LPQKLYGLY-GCANEISRGKDTPSGHWEIAGTPVTFDWGYFPTEGDAFPPELVEAICKAADLPG  142
sp|Q2KDR4|DEOB_RHIEC              RAASG-QFP-AGMP--LPEKVYGVY-GTANEISRGKDTPSGHWEIAGTPVNFDWGYFPTEGDAFPPELIEALCRAGDVPG  142
sp|B5ZXI4|DEOB_RHILW              RSASG-RFP-AGMP--IPEKVYGIY-GAATEISRGKDTPSGHWEIAGTPVSFDWGYFPTEGDAFPEELIEALCRDAGVSG  142
sp|Q1MMV6|DEOB_RHIL3              RSASG-RFP-AGMP--VPEKVYGIY-GAATEISRGKDTPSGHWEIAGTPVSFDWGYFPIEGDAFPQEFIEALCREADVPG  142
sp|Q98BG5|DEOB_RHILO              ETATG-----LGFAHFGTDLIANAFHGAAQEVSSGKDTPSGHWEIAGLPVRFDWGYFPDTVPAFPADLTAAMIREGEVPG  142
sp|Q11AV9|DEOB_CHESB              AIAAG-REPERGFA--S----TGFF-GAAEERSTGKDTPSGHWEIAGVPVPFEWGYFPETTPSFPAELTGRLIREASLPG  139
sp|A5IGR8|DEOB_LEGPC              MASSG-LPF-IDLS--ALAIPSGYY-GYAVEQSLGKDTPSGHWEMAGVPVTFEWGYFPDKPYCFPEELISEFIKQCNLPG  143
sp|Q5X7B2|DEOB_LEGPA              MASSG-LPF-IDLS--ALAIPSGYY-GYAVEQSLGKDTPSGHWEMAGVPVTFEWGYFPDKSYCFPEELISEFIKQCNLPG  143
sp|Q5WYR0|DEOB_LEGPL              MASSG-LPF-IDLA--TLAIPSGYY-GYAVEQSLGKDTPSGHWEMAGVPVTFEWGYFPDKPYCFPEELISEFIKQCNLPG  143
sp|Q5ZXU2|DEOB_LEGPH              MASSG-LSF-IDLA--TLAIPSGYY-GYAVEQSLGKDTPSGHWEMAGVPVTFEWGYFPDKPYCFPEELISEFIKQCNLPG  143
sp|B2KBT7|DEOB_ELUMP              EASAGVKIE-TGPQPPAQVNIPSKY-GFMREQSHGKDTLSGHWEMAGVPVLFDWGYFKPGYPSFPKELIEQICKEAGIDK  133
sp|Q2SHN3|DEOB_HAHCH              EASCG-AAA-PGLE--QAPEIIGTY-GFAEELSSGKDTPSGHWEIAGVPVLFDWGYFTDKQQSFPPELLDAIIKEARLPG  141
sp|Q1QZC2|DEOB_CHRSD              HLATG-EWA-EGIT--PPETLDGAY-AAAAEISSGKDTPSGHWEIAGVPALFEWGYFPDKTQSFPPDLLEALIEQAKLPG  134
sp|A1SYK3|DEOB_PSYIN              AESSS-TFP-AGLD--PLVEPIAAY-GYAKEISTAKDTSSGHWEITGVPVLFDWGYFDKKQNSFPQELLDELVERADLPG  150
sp|A6VM02|DEOB_ACTSZ              KESCG-ELP-KHFQ--NQPHLIGGY-AFAREISSGKDTTSGHWEIAGVPVLFDWGLFPDKQNSFPKPLLDRIVAKAGIKG  130
sp|C4LAY9|DEOB_TOLAT              AASCG-EFP-AGLD--ASVTPIAAY-GYARELSSGKDTPSGHWEMAGAPVLFDWGYFTDKQNSFPPELLDALVAQGNLPG  140
sp|A4SRU2|DEOB_AERS4              ELASG-YFP-AGLK--KDIEVVGAY-GFAQELSSGKDTPSGHWEIAGVPVLFEWGYFKDHHNSFPQELLDAIVEKAGLSG  138
sp|A0KPE2|DEOB_AERHH              ELASG-TFP-AGLN--KDIDVVGAY-GFAQELSSGKDTPSGHWEIAGVPVLFEWGYFHDHHNSFPQELLDAIVEKAGLPG  138
```

```
sp|Q7NRT1|DEOB_CHRVO         QLSSG-YFP-EGMD---PAAPAAAY-GYAREISSGKDTPSGHWEIAGVPVLFDWGYFSDHDNSFPQQLLDAIVDKAGLPG   139
sp|Q086F8|DEOB_SHEFN         FESTG-KFA-AGFA--DNVEVIGAY-GHADELSSGKDTPSGHWEMAGVPVLYEWGYFSDLTNSFPKELTDKILERAGLDG   140
sp|Q12QG1|DEOB_SHEDO         FESTG-QFA-PGFS--QDVEVVGAY-GYADELSTGKDTPSGHWEMAGVPVLYDWGYFNDLQDSFPKELTDKILARAGLSG   140
sp|A1S476|DEOB_SHEAM         KEATG-AFA-PGFG--DNVDIIGAY-GHCQELSSGKDTPSGHWEMAGVPVLFEWGYFSEHQNSFPKELTDKILERAGLTE   140
sp|Q8EHK2|DEOB_SHEON         MESTG-AFA-PGFA--DNVDLIGAY-GHAQELSSGKDTPSGHWEMAGVPVLFEWGYFSEHQNSFPKELTDKILARAGLDG   140
sp|A0KU09|DEOB_SHESA         MESTG-AFA-PGFA--DNVELIGAY-GHAQELSSGKDTPSGHWEMAGVPVLFEWGYFSEHQNSFPKELTDKILARAGLDG   140
sp|Q0HLE8|DEOB_SHESM         MESTG-AFA-PGFA--DDVELIGAY-GHAQELSSGKDTPSGHWEMAGVPVLFDWGYFSEHQNSFPKELTDKILARAGLDG   140
sp|Q0HXQ2|DEOB_SHESR         MESTG-AFA-PGFA--DDVELIGAY-GHAQELSSGKDTPSGHWEMAGVPVLFDWGYFSEHQNSFPKELTDKILARAGLDG   140
sp|A1RH89|DEOB_SHESW         MESTG-AFA-PGFG--DNVELIGAY-GHAQELSSGKDTPSGHWEMAGVPVLFEWGYFSEHQNSFPKELTDKILARAGLDG   140
sp|A4Y9A6|DEOB_SHEPC         MESTG-AFA-PGFG--DNVELIGAY-GHAQELSSGKDTPSGHWEMAGVPVLFEWGYFSEHQNSFPKELTDKILARAGLDG   140
sp|A6WRB6|DEOB_SHEB8         MESTG-AFA-PGFG--DNVELIGAY-GHAQELSSGKDTPSGHWEMAGVPVLFEWGYFSEHQNSFPKELTDKILARAGLDG   140
sp|A9KZ78|DEOB_SHEB9 (2)     MESTG-AFA-PGFG--DNVELIGAY-GHAQELSSGKDTPSGHWEMAGVPVLFEWGYFSEHQNSFPKELTDKILARAGLDG   140
sp|B8E6P6|DEOB_SHEB2         MESTG-AFA-PGFG--DNVELIGAY-GHAQELSSGKDTPSGHWEMAGVPVLFEWGYFSEHQNSFPKELTDKILARAGLDG   140
sp|B0TQ89|DEOB_SHEHH         KESTG-EFP-AGFG--DDVEIIGAY-GHADELSSGKDTPSGHWEMAGVPVLYEWGYFGDKQNSFPKELTDKILARAGLSG   140
sp|A8H726|DEOB_SHEPA         KESTG-EFP-AGFG--DDVEVIGAY-GHADELSSGKDTPSGHWEMAGVPVLYEWGYFSELTNSFPQALTDKILARAGLTE   140
sp|A8FYQ7|DEOB_SHESH         KESTG-QLP-AGFS--DDVEIIGAY-GHADELSTGKDTPSGHWELAGVPVLYEWGYFSDHTNSFPKELTDKILARAGLSD   140
sp|A3QGT1|DEOB_SHELP         KESTG-SFA-EGFG--DDVEVIGAY-GHADELSSGKDTPSGHWEMAGVPVLYEWGYFSDLKDSFPKELTDKILERAGLSG   140
sp|B1KRP6|DEOB_SHEWM         KESTG-TFA-PGFS--DDVEVIGAY-GHADELSTGKDTPSGHWEMAGVPVLYEWGYFSDLTNSFPKELTDKILARAGLDG   140
sp|Q6LUH2|DEOB_PHOPR         EESSG-YFP-EGLD--PNVEITGAY-GHAKELSSGKDTPSGHWEIAGVPVLFDWGYFSDHDNSFPKELTDRILKRANLPG   141
sp|B6ENG5|DEOB_ALISL         KESTG-KLA-AGLD--ADAEIIGAY-GHAAELSSGKDTPSGHWEIAGVPVLFDWGYFSDKENSFPKELTDRILARANLSG   141
sp|B5FAA0|DEOB_ALIFM         KESTG-KFS-AGLD--ANAEIIGAY-GHAAELSSGKDTPSGHWEIAGVPVLFDWGYFTDKENSFPKELTDRILERANLPG   141
sp|Q5E7J5|DEOB_ALIF1         KESTG-KFS-AGLD--ANAEIIGAY-GHAAELSSGKDTPSGHWEIAGVPVLFDWGYFTDKENSFPKELTDRILERANLPG   141
sp|C3LQC0|DEOB_VIBCM (2)     KESTG-RFA-PGLD--ERADIIGAY-AHAAELSSGKDTPSGHWEIAGVPVLFEWGYFSDKQNSFPKELTDRILARAGLDG   141
sp|Q9KPL9|DEOB_VIBCH         KESTG-RFA-PGLD--ERADIIGAY-AHAAELSSGKDTPSGHWEIAGVPVLFEWGYFSDKQNSFPKELTDRILARAGLDG   141
sp|A7MUW4|DEOB_VIBCB         KESTG-RFA-PGLD--ADAEIIGAY-GHAAELSSGKDTPSGHWEIAGVPVLFDWGYFTDKANSFPKELTDRILERAGLDG   141
sp|Q7MI40|DEOB_VIBVY (2)     KESTG-RFA-PGLD--ADAEIIGAY-GHAAELSSGKDTPSGHWEIAGVPVLFEWGYFTDKANSFPKELTDRILARAGIDG   141
sp|Q87M24|DEOB_VIBPA         KESTG-RFA-PGLD--ADAEIIGAY-GHAAELSSGKDTPSGHWEIAGVPVLFDWGYFTDKANSFPKELTDRILERAGLDG   141
sp|Q2NW04|DEOB_SODGM         EQSTG-RFP-AGLD--KQAEIVGAY-AYASEISSGKDTLSGHWEIAGVPVLFDWGYFPAVENSFPPKLLEALVARAGLPG   140
sp|B2VH52|DEOB_ERWT9         EASSG-KFP-PGLD--KNAEIIGAY-AYASELSSGKDTPSGHWEIAGVPVLFDWGYFSDTENSFPQELLDLLVEKANLPG   140
sp|A7MGA8|DEOB_CROS8         EGATG-FIP-AGMD--GNAEITGAY-AWAHELSSGKDTPSGHWEIAGVPVLFDWGYFTDEKNSFPQELLDKLVERANLPG   140
sp|A4W6A0|DEOB_ENT38         EGSTG-KVA-AGMD--ANAEVVGAY-AWAHELSSGKDTPSGHWEIAGVPVLFDWGYFSDHENSFPQKLLDKLVERGNLPG   140
sp|Q327L3|DEOB_SHIDS         EGSTG-FIP-AGMD--GNAEVIGAY-AWAHEMSSGKDTPSGHWEIAGVPVLFEWGYFSDHENSFPQELLDKLVERANLPG   140
sp|Q1R260|DEOB_ECOUT (3)     EGSTG-FIP-AGMD--GNAEVIGAY-AWAHEMSSGKDTPSGHWEIAGVPVLFEWGYFSDHENSFPQELLDKLVERANLPG   140
sp|B7UR11|DEOB_ECO27         EGSTG-FIP-AGMD--GNAEVIGAY-AWAHEMSSGKDTPSGHWEIAGVPVLFEWGYFSDHENSFPQELLDKLVERANLPG   140
sp|Q3YU10|DEOB_SHISS         EGSTG-FIP-AGMD--GNAEVIGAY-AWAHEMSSGKDTPSGHWEIAGVPVLFEWGYFSDHENSFPQELLDKLVERANLPG   140
sp|Q0SX28|DEOB_SHIF8 (22)    EGSTG-FIP-AGMD--GNAEVIGAY-AWAHEMSSGKDTPSGHWEIAGVPVLFEWGYFSDHENSFPQELLDKLVERANLPG   140
sp|B5Y275|DEOB_KLEP3         EGSTG-KIA-AGMD--GNAEVIGAY-AWAHELSSGKDTPSGHWEIAGVPVLFDWGYFSDHENSFPQELLDKLVKRANLPG   140
sp|A9MRA5|DEOB_SALAR         EGSTG-KIA-AGMD--GNADVIGAY-AWAHELSSGKDTPSGHWEIAGVPVLFDWGYFSDHENSFPQELLDKLVKRANLPG   140
sp|B5BAJ9|DEOB_SALPK (2)     EGSTG-KIA-AGMD--GNADVIGAY-AWAHELSSGKDTPSGHWEIAGVPVLFDWGYFSDHENSFPQELLDKLVKRANLPG   140
sp|C0Q7M5|DEOB_SALPC (2)     EGSTG-KIA-AGMD--GNADVIGAY-AWAHELSSGKDTPSGHWEIAGVPVLFDWGYFSDHENSFPQELLDKLVKRANLPG   140
sp|B5R9V1|DEOB_SALG2         EGSTG-KIA-AGMD--GNADVIGAY-AWAHELSSGKDTPSGHWEIAGVPVLFDWGYFSDHENSFPQELLDKLVKRANLPG   140
sp|B5FTC7|DEOB_SALDC         EGSTG-KIA-AGMD--GNADVIGAY-AWAHELSSGKDTPSGHWEIAGVPVLFDWGYFSDHENSFPQELLDKLVKRANLPG   140
sp|P63924|DEOB_SALTI (8)     EGSTG-KIA-AGMD--GNADVIGAY-AWAHELSSGKDTPSGHWEIAGVPVLFDWGYFSDHENSFPQELLDKLVKRANLPG   140
sp|C5BHJ4|DEOB_EDWI9         LESTG-RFP-EGLD--ENAEVIGAY-GYANELSSGKDTPSGHWEIAGVPVLFDWGYFHEHQNSFPQALLDTLVERANLPG   140
sp|Q6D990|DEOB_PECAS         EASTG-TFP-AGLD--ENADIIGAY-AHASEISSGKDTPSGHWEIAGVPVLFDWGYFKDEENSFPQELLDKLVKRANLPG   140
sp|C6DKL9|DEOB_PECCP         EASTG-TFP-AGLD--ENADIIGAY-AHASEISSGKDTPSGHWEIAGVPVLFDWGYFKDEENSFPQDLLDKLVKRANLPG   140
sp|B4EWA2|DEOB_PROMH         EESTG-KFP-VGLD--KDAEIIGAY-GYASEISSGKDTPSGHWEIAGVPVLFDWGYFKDLKNSFPQELLDNIVKRANLPG   141
sp|Q7N931|DEOB_PHOLL         EESCG-TFP-AGLD--KDADIIGAY-AYASELSSGKDTPSGHWEIAGVPVLFDWGYFKDEENSFPQALLDKLVERANLPG   140
sp|A8G9H8|DEOB_SERP5         EESTG-TFP-QGLD--RNADIIGAY-AHASELSSGKDTPSGHWEIAGVPVLFDWGYFSDEHNSFPQELLDKLVERANLPG   140
sp|A1JJ99|DEOB_YERE8         EESTG-KFP-VGLD--KNADIIGAY-GYASELSSGKDTPSGHWEIAGVPVLFDWGYFSDVENSFPQELLDKLVKRANLPG   140
sp|A9R047|DEOB_YERPG         EESTG-TFP-VGLD--KNADIIGAY-GYASELSSGKDTPSGHWEIAGVPVLFDWGYFSDVENSFPQELLDKLVKRANLSG   140
sp|A4TQJ1|DEOB_YERPP (4)     EESTG-TFP-VGLD--KNADIIGAY-GYASELSSGKDTPSGHWEIAGVPVLFDWGYFSDVENSFPQELLDKLVKRANLPG   140
sp|B1JL35|DEOB_YERPY (4)     EESTG-TFP-VGLD--KNADIIGAY-GYASELSSGKDTPSGHWEIAGVPVLFDWGYFSDVENSFPQELLDKLVKRANLPG   140
```

```
Consensus                    YLGNCHASGTVILDQLGEEHMKTGKPIFYTSADSVFQIACHEETFGLDRLYELCEIAREELEP--YNIGRVIARPFIGDK   221
deoB_Bcereus_WP_098782964.1  IIGNKPASGTEILDELGQEQMETGSLIVYTSADSVLQIAAHEEVVPLDELYKICKIARELTLDEKYMVGRVIARPFVGE-   202
deoB_Rel606                  YLGNCHSSGTVILDQLGEEHMKTGKPIFYTSADSVFQIACHEGTFGLDKLYELCEIAREELTNGGYNIGRVIARPFIGDK   220
sp|Q89A57|DEOB_BUCBP         FLGNCHASGTDILDRFGEIHISTKKPILYTSIDSVCQIACHESIFGLKRLYNLCRSIRKIFDKRKINIARIIARPFTGFK   221
sp|Q8K936|DEOB_BUCAP         ILGNCHASGTEIIKILGEEHIKTGKPIFYTSCDSVFQIACHEIKFGLSNLYKICENIREILDQNKYQVARVIARPFIGTN   222
sp|B8D866|DEOB_BUCAT         FIGNCHASGTDIISRLGEEHIQTKKPIVYTSSDSVFQIACHEEFFGLSNLYKLCKTVRFILDRYNYKVARVIARPFIGND   220
sp|P57607|DEOB_BUCAI         FIGNCHASGTDIISRLGEEHIQTKKPIVYTSSDSVFQIACHEEFFGLSNLYKLCKTVRFILDRYNYKVARVIARPFIGND   220
sp|B8D9W4|DEOB_BUCA5         FIGNCHASGTDIISRLGEEHIQTKKPIVYTSSDSVFQIACHEEFFGLSNLYKLCKTVRFILDQYNYKVARVIARPFIGND   220
sp|Q483R0|DEOB_COLP3         ILGNCHASGTEILTRLGQEHIETGLPICYTSADSVFQIAAHEEHFGLDNLYKYCETVRELLGD--LNIGRVIARPFVGDS   212
sp|Q3ICU9|DEOB_PSET1         ILGNCHASGTTILEQLGEEHVKTGKPICYTSADSVFQIAAHEESFGLEKLYQVCETARSLLDE--MNIGRVIARPFLGSN   212
sp|Q5QXT9|DEOB_IDILO         YLGNCHASGTEILKQLGEQHIRSSYPIFYTSADSVFQIAAHEEHFGLERLYQVCEKVRALIEP--YNIGRVIARPFVGDK   211
sp|Q17YP1|DEOB_HELAH         YLGNCHASGTEIIKDLGEKHLETLYPIFYTSADSVFQIAVHEEKFGLDKLYALCEEVFEILEP--LKIARVIARSFIGTN   220
sp|Q9ZK37|DEOB_HELPJ         YLGNCHASGTEIIKDLGEKHLETLYPIFYTSADSVFQIAAHEERFGLDHLYALCEEAFQILEP--LKIARVIARPFIGTN   220
sp|B6JN18|DEOB_HELP2         YLGNCHASGTEIIKDLGEKHLETLYPIFYTSADSVFQIAAHEEKFGLDNLYALCEEAFQILEP--LKIARVIARPFIGTN   220
sp|P56195|DEOB_HELPY         YLGNCHASGTEIIKDLGEKHLETLYPIFYTSADSVFQIAAHEERFGLDNLYALCEEAFQILEP--LKIARVIARPFIGTN   220
sp|B5Z8H3|DEOB_HELPG         YLGNCHASGTEIIKDLGEKHLETLYPIFYTSADSVFQIAAHEEKFGLDNLYALCEEVFQILEP--LKIARVIARPFIGAN   220
sp|B2UUU2|DEOB_HELPS         YLGNCHASGTEIIKDLGEKHLETLYPIFYTSADSVFQIVAHEEKFGLDNLYALCEEAFQILEP--LKIARVIARPFIGAN   220
sp|Q1CS87|DEOB_HELPH         YLGNCHASGTEIIKDLGEKHLETLYPIFYTSADSVFQIAVHEEKFGLDNLYALCEEAFQILEP--LKIARVIARPFIGAN   220
sp|A4YPV1|DEOB_BRASO         ILGNRHASGTDIISDLGEAHLQTGQPICYTSADSVFQIAAHEEAFGLDRLYRLCEATRPLVDR--FNIGRVIARPFVGRS   218
sp|B8IJW7|DEOB_METNO         ILGDCHASGTAIIEELGAEQVRTGKPICYTSVDSVFQIAAHEEAFGLERLSAVCAVARRLCDP--YRIGRVIARPFTGDA   220
sp|B0UPC1|DEOB_METS4         ILGDCHASGTAVIEAYGAEHLRSGKPICYTSVDSVFQIAAHEEAFGLERLYALCGIARRLCDP--YRIGRVIARPFAGSA   220
sp|B1LYJ9|DEOB_METRJ         ILGDCHAPGVAIIDALGAEHLRTGRPICYTSADSVFQIAAHEEAFGLERLYDLCRVARRLCDR--YRVCRVIARPFVGSP   220
sp|B1ZIG8|DEOB_METPB         ILGDRHASGTAIIDALGAEHVRTGRPICYTSADSVFQIAAHEEAFGLEQLYETCRIARAVCDP--YRVGRVIARPFLGSE   220
sp|A9W427|DEOB_METEP         ILGDCHASGTAIIEALGAEHVRTGKPICYTSADSVFQIAAHEEAFGLERLYETCRIAREVCDP--YRVGRVIARPFLGSA   220
sp|B7KXI3|DEOB_METC4         ILGDCHASGTAIIEALGAEHVRTGKPICYTSADSVFQIAAHEEAFGLERLYETCRIAREVCDP--YRVGRVIARPFLGSA   220
sp|C3MBH5|DEOB_SINFN         ILGNCHASGTDIIARHGEEHMRSGKPICYTSSDSVFQIAAHERTFGLERLLELCQVVRRLVDD--YNIGRVIARPFVGDN   220
sp|Q92T47|DEOB_RHIME         ILGNCHASGTDIIARHGEEHMRSGKPICYTSSDSVFQIAAHEQTFGLERLLNLCEVVRRLVDD--YNIGRVIARPFVGSD   220
sp|A6UET6|DEOB_SINMW         ILGNCHASGTDIIARLGEEHMRTGKPICYTSSDSVFQIAAHEQTFGLERLQDLCAVVRRLVDE--YNIGRVIARPFVGSD   220
sp|B9JYP7|DEOB_AGRVS         ILGNCHASGTDILARLGEEHCRTGQPICYTSSDSVFQIAAHEHVFGLERLLRLCEIVRELLTP--YRIGRVIARPFIGNS   220
sp|Q8UJ04|DEOB_AGRFC         ILGNCHASGTEIIAALGEEHIRSGKPICYTSSDSVFQIAAHETHFGLDRLIALCETVRKLLDP--LNIGRVIARPFIGET   220
sp|B9J6V9|DEOB_AGRRK         ILGNCHASGTDIIARYGEEHIRSGKPICYTSSDSVFQIAAHEQHFGLERLISLCQIVRTLLDP--YNIGRVIARPFIGET   220
sp|Q2KDR4|DEOB_RHIEC         ILGNCHASGTEIIARLGEEHIRTGKPICYTSSDSVFQVAAQEAHFGLDRLLTFCRIARGLLDP--YNIGRIIARPFIGQS   220
sp|B5ZXI4|DEOB_RHILW         ILGNCHASGTEIIARLGEEHIRTGKPICYTSSDSVFQVAAHEVHFGLDRLLAFCQVARGLLDR--YNIGRVIARPFIGHS   220
sp|Q1MMV6|DEOB_RHIL3         ILGNCHASGTEIIARLGEEHIRTGKPICYTSSDSVFQVAAHEVHFGLDRLLAFCGLARGLLDS--YNIGRVIARPFIGQS   220
sp|Q98BG5|DEOB_RHILO         ILGNCHAPGTEIIERFGEEHIRTGKPICYTSVDSVLQIAAHEVHFGLDRLYEFCQVVRRLVDP--LRIGRVIARPFVGET   220
sp|Q11AV9|DEOB_CHESB         ILANCHASGTEVIARLGEEHIRTGKPICYTSADSVFQIAAHETHFGLDRLYAVCETARRLVDD--YRIGRVIARPFVGES   217
sp|A5IGR8|DEOB_LEGPC         VLGEKHASGTIIIDELGEEHIRTGKPIVYTSADSVFQIAAHEEAFGLQRLYDICKIARNLVDK--YQIGRVIARPFAG-K   220
sp|Q5X7B2|DEOB_LEGPA         VLGEKHASGTIIIDELGEEHIRTGKPIVYTSADSVFQIAAHEEAFGLQRLYDICKIARNLVDK--YQIGRVIARPFTG-K   220
sp|Q5WYR0|DEOB_LEGPL         VLGEKHASGTIIMDELGEEHIRTGKPIVYTSADSVFQIAAHEEAFGLQRLYDICKIARNLVDK--YQIGRVIARPFTG-K   220
sp|Q5ZXU2|DEOB_LEGPH         VLGEKHASGTIIIDELGEKHIRTGKPIVYTSADSVFQIAAHEEAFGLQRLYDICKIARNLVDK--YQIGRVIARPFTG-K   220
sp|B2KBT7|DEOB_ELUMP         ILGNKAASGTEILEELGEEHIKTGKPICYTSADSVFQIAAHEKHFGLERLYTICEIAFKYLKP--YKIARVIARPFEGER   211
sp|Q2SHN3|DEOB_HAHCH         VLGNVHASGTQIIKDLGMEHRATGKPIFYTSADSVVQIACHEETFGLQRLYELCQITRKHIDP--YNIARVIARPFVGAS   219
sp|Q1QZC2|DEOB_CHRSD         VLGNCHASGMPILETLGEAHIASGKPIVYTSADSVFQIAAHETHFGLDRLYALCEIARELLMP--YNIGRVIARPFVGET   212
sp|A1SYK3|DEOB_PSYIN         YLGNCHASGTTILEDLGEEHLKTGKPIFYTSADSVFQIACHEETFGLERLYKLCELTRELVNK--YNIGRVIARPFSGSA   228
sp|A6VM02|DEOB_ACTSZ         YLGNCHSSGTVILDQLGEEHMKTGLPIFYTSADSVFQIAAHEETFGLNNLYELCEIVRTELEG--YNIGRVIARPFIGNK   208
sp|C4LAY9|DEOB_TOLAT         YLGNCHASGTEILDRLGEEHMRSGKPIFYTSADSVFQIACHEETYGLDKLYELCKLARQLLEP--YNIGRVIARPFVGSK   218
sp|A4SRU2|DEOB_AERS4         YLGNCHASGTQVLDDLGEEHMSTGKPILYTSADSVFQIACHEETYGLEKLYELCHIVRELLEP--YNIGRVIARPFVGSG   216
sp|A0KPE2|DEOB_AERHH         YLGNCHASGTQVLDDLGEEHMRTGKPILYTSADSVFQIACHEETYGLEKLYELCHIVRELLEP--YNIGRVIARPFVGSG   216
sp|Q7NRT1|DEOB_CHRVO         YLGNCHASGTEILDRLGEEHMKTGKPIFYTSADSVFQIACHEETYGLERLYELCKITRELLEP--YNIGRVIARPFVGEG   217
sp|Q086F8|DEOB_SHEFN         FLGNCHASGTQILEELGEEHMKTGKPIFYTSADSVFQIACHEESFGVENLYNLCKIAREELEP--YNIGRVIARPFVGTG   218
sp|Q12QG1|DEOB_SHEDO         FLGNCHASGTTILEQLGEEHMRSGMPIFYTSADSVFQVACHEESFGLDNLLRLCEIAREELGP--YNIGRVIARPFIGTG   218
sp|A1S476|DEOB_SHEAM         FLGNCHASGTTILEELGEEHMKTGKPIFYTSADSVFQIACHEESFGLENLYRLCEIAREELEP--YNIGRVIARPFVGTG   218
sp|Q8EHK2|DEOB_SHEON         FLGNCHASGTTILEELGEEHMRSGKPIFYTSADSVFQIACHEGTFGLENLYRLCEIAREELEP--YNIGRVIARPFDGTG   218
sp|A0KU09|DEOB_SHESA         FLGNCHASGTTILEELGEEHMRSGKPIFYTSADSVFQIACHEGTFGLENLYRLCEIAREELEP--YNIGRVIARPFDGTG   218
sp|Q0HLE8|DEOB_SHESM         FLGNCHASGTTILEELGEEHMRSGKPIFYTSADSVFQIACHEGTFGLENLYRLCEIAREELEP--YNIGRVIARPFDGTG   218
sp|Q0HXQ2|DEOB_SHESR         FLGNCHASGTTILEELGEEHMRSGKPIFYTSADSVFQIACHEGTFGLENLYRLCEIAREELEP--YNIGRVIARPFDGTG   218
sp|A1RH89|DEOB_SHESW         FLGNCHASGTTILEELGEEHMRSGKPIFYTSADSVFQIACHEETFGLDNLYRLCEITREELAP--YNIGRVIARPFNGTG   218
sp|A4Y9A6|DEOB_SHEPC         FLGNCHASGTTILEELGEEHMRSGKPIFYTSADSVFQIACHEETFGLDNLYRLCEITREELAP--YNIGRVIARPFNGTG   218
sp|A6WRB6|DEOB_SHEB8         FLGNCHASGTTILEELGEEHMRSGKPIFYTSADSVFQIACHEETFGLDNLYRLCEITREELEP--YNIGRVIARPFDGTG   218
sp|A9KZ78|DEOB_SHEB9 (2)     FLGNCHASGTTILEELGEEHMRSGMPIFYTSADSVFQIACHEETFGLDNLYRLCEITREELEP--YNIGRVIARPFDGTG   218
sp|B8E6P6|DEOB_SHEB2         FLGNCHASGTTILEELGEEHMRSGKPIFYTSADSVFQIACHEETFGLDNLYRLCAITREELEP--YNIGRVIARPFDGTG   218
sp|B0TQ89|DEOB_SHEHH         FLGNCHSSGTIILEELGEEHMRTGKPIFYTSADSVFQIACHEESFGLEKLYELCIIAREELAD--YNIGRVIARPFVGTG   218
sp|A8H726|DEOB_SHEPA         FLGNCHSSGTVILDELGEEHMRTGKPIFYTSADSVFQIACHEESFGLEKLYELCIIAREELAD--YNIGRVIARPFVGTG   218
sp|A8FYQ7|DEOB_SHESH         YLGNCHASGTAILEELGNEHMTSGLPIFYTSADSVFQVACHEETFGLENLYTLCQIVREELEP--YNIGRVIARPFVGTG   218
sp|A3QGT1|DEOB_SHELP         FLGNCHASGTAILEELGEEHMTSGLPIFYTSADSVFQIACHEETFGLENLYTLCQIAREELEP--YNIGRVIARPFVGTG   218
sp|B1KRP6|DEOB_SHEWM         YLGNCHASGTAILEELGEEHMRTGKPIFYTSADSVFQIACHEESFGLENLYNLCIIAREELEP--YNIGRVIARAFVGTG   218
sp|Q6LUH2|DEOB_PHOPR         FLGNCHASGTHVLDELGEEHMKTGMPIFYTSADSVFQIACHEETFGLDNLLTLCQIAREELED--YNIGRVIARPFTGPG   219
sp|B6ENG5|DEOB_ALISL         YLGNCHASGTQVLDDLGEEHMKTGMPIFYTSADSVFQIACHEETFGLDNLLTLCQIAREELED--YNIGRVIARPFIGAG   219
sp|B5FAA0|DEOB_ALIFM         YLGNCHASGTQVLDDLGEEHMKTGMPIFYTSADSVFQIACHEETFGLDNLLTLCQIAREELED--YNIGRVIARPFIGPG   219
sp|Q5E7J5|DEOB_ALIF1         YLGNCHASGTQVLDDLGEEHMKTGMPIFYTSADSVFQIACHEETFGLDNLLTLCQIAREELED--YNIGRVIARPFIGPG   219
sp|C3LQC0|DEOB_VIBCM (2)     FLGNCHASGTQVLDDLGEEHMRTGKPIFYTSADSVFQIACHEETFGLARLLELCQIAREELAD--YNIGRVIARPFVGPG   219
sp|Q9KPL9|DEOB_VIBCH         FLGNCHASGTQVLDDLGEEHMRTGKPIFYTSADSVFQIACHEETFGLDRLLELCQIAREELAD--YNIGRVIARPFVGPG   219
sp|A7MUW4|DEOB_VIBCB         FLGNCHASGTQVLDDLGEEHMKTGKPIFYTSADSVFQIACHEETFGLDRLLELCQIAREELED--YNIGRVIARPFVGAG   219
sp|Q7MI40|DEOB_VIBVY (2)     FLGNCHASGTQVLDDLGEEHMKTGQPIFYTSADSVFQIACHEETFGLDRLLELCQIAREELAD--YNIGRVIARPFIGPG   219
sp|Q87M24|DEOB_VIBPA         FLGNCHASGTQVLDDLGEEHMKTGQPIFYTSADSVFQIACHEETFGLDRLLELCQIAREELED--YNIGRVIARPFIGPG   219
sp|Q2NW04|DEOB_SODGM         FLGNCHASGTVILDRLGEEHMRTGKPIFYTSADSVFQLACHEETFGLERLYSLCEIAREILTDGGYNIGRVITRPFVGAK   220
sp|B2VH52|DEOB_ERWT9         YLGNCHSSGTVILDQLGAEHMKSGKPIFYTSADSVFQIACHEETFGLERLYALCEIAREALTEGGYNIGRVIARPFVGDK   220
sp|A7MGA8|DEOB_CROS8         YLGNCHSSGTVILDELGEEHMKTGKPIFYTSADSVFQIACHEETFGLDRLYELCEIAREELTEGGYNIGRVIARPFVGDK   220
sp|A4W6A0|DEOB_ENT38         YLGNCHSSGTVILDQLGEEHMKTGKPIFYTSADSVFQIACHEETFGLDRLYELCEIAREELTEGGYNIGRVIARPFIGNK   220
sp|Q327L3|DEOB_SHIDS         YLGNCHSSGTVILDQLGEEHMKTGKPIFYTSADSVFQIACHEETFGLDKLYELCEIAREELINGGYNIGRVIARPFIGDK   220
sp|Q1R260|DEOB_ECOUT (3)     YLGNCHSSGTVILDQLGEEHMKTGKPIFYTSADSVFQIACHEETFGLDKLYELCEIAREELTNGGYNIGRVIARPFIGDK   220
sp|B7UR11|DEOB_ECO27         YLGNCHSSGTVILDQLGEEHMKTGKPIFYTSADSVFQIACHEETFGLDKLYKLCEIAREELTNGGYNIGRVIARPFIGDK   220
sp|Q3YU10|DEOB_SHISS         YLGNCHSSGTVILDQLGEEHMKTGKPIFYTSADSVFQIACHEETFGLDKLYELCEIAREELTNGGYNIGRVIARPFIGDK   220
sp|Q0SX28|DEOB_SHIF8 (22)    YLGNCHSSGTVILDQLGEEHMKTGKPIFYTSADSVFQIACHEETFGLDKLYELCEIAREELTNGGYNIGRVIARPFIGDK   220
sp|B5Y275|DEOB_KLEP3         YLGNCHSSGTVILDQLGEEHMKTGKPIFYTSADSVFQIACHEETFGLDKLYELCEIAREELTEGGYNIGRVIARPFIGDK   220
sp|A9MRA5|DEOB_SALAR         YLGNCHSSGTVILDQLGEEHMKTGKPIFYTSADSVFQIACHEETFGLDKLYELCEIAREELTEGGYNIGRVIARPFIGDK   220
sp|B5BAJ9|DEOB_SALPK (2)     YLGNCHSSGTVILDQFGEEHMKTGKPIFYTSADSVFQIACHEETFGLDKLYELCEIAREELTEGGYNIGRVIARPFIGDK   220
sp|C0Q7M5|DEOB_SALPC (2)     YLGNCHSSGTVILDQLGEEHMKTGKPIFYTSADSVFQIACHEETFGLDKLYELCEIAREELTEGGYNIGRVIARPFIGDK   220
sp|B5R9V1|DEOB_SALG2         YLGNCHSSGTVILDQLGEEHMKTGKPIFYTSADSVFQIACHEETFGLDKLYELCEIAREELTEGGYNIGRVIARPFIGDK   220
sp|B5FTC7|DEOB_SALDC         YLGNCHSSGTVILDQLGEEHMKTGKPIFYTSADSVFQIACHEETFGLDKLYELCEIAREELTVGGYNIGRVIARPFIGDK   220
sp|P63924|DEOB_SALTI (8)     YLGNCHSSGTVILDQLGEEHMKTGKPIFYTSADSVFQIACHEETFGLDKLYELCEIAREELTEGGYNIGRVIARPFIGDK   220
sp|C5BHJ4|DEOB_EDWI9         YLGNCHSSGTVILDQLGEEHMKSGKPIFYTSADSVFQIACHEETFGLERLYELCEIARDELNKGGYNIGRVIARPFVGDK   220
sp|Q6D990|DEOB_PECAS         YLGNCHSSGTVILDQLAEEHMKTGKPIFYTSADSVFQIACHEETFGLDKLYELCEIAREELTEGDYNIGRVIARPFIGDK   220
sp|C6DKL9|DEOB_PECCP         YLGNCHSSGTVILDQLAEEHMKTGKPIFYTSADSVFQIACHEETFGLDKLYELCEIAREELTEGGYNIGRVIARPFIGDK   220
sp|B4EWA2|DEOB_PROMH         YLGNCHASGTVILDELGEEHMKTGKPIFYTSADSVFQIACHEETFGLDKLYELCEIARDELNKGDYNIGRVIARPFIGDK   221
sp|Q7N931|DEOB_PHOLL         YLGNCHSSGTVILDKLGEEHMKTGKPIFYTSADSVFQIACHEETFGLDRLYELCEIAREELTDGGYNIGRVIARPFVGDK   220
sp|A8G9H8|DEOB_SERP5         YLGNCHSSGTVILDQLGEEHMKTGKPIFYTSADSVFQIACHEETFGLDRLYELCEIAREELTEGGYNIGRVIARPFLGDK   220
sp|A1JJ99|DEOB_YERE8         YLGNCHSSGTVILDQLGEEHMKTGKPIFYTSADSVFQIACHEETFGLDRLYELCEIAREELTEGGYNIGRVIARPFIGDK   220
sp|A9R047|DEOB_YERPG         YLGNCHSSGTVILDQLGEEHMKTGKPIFYTSADSVFQIACHEETFGLDRLYELCEIAREELTDGGYNIGRVIARPFIGDK   220
```

```
sp|A4TQJ1|DEOB_YERPP (4)      YLGNCHSSGTVILDQLGEEHMKTGKPIFYTSADSVFQIACHEETFGLDRLYELCEIAREELTDGGYNIGRVIARPFIGDK  220
sp|B1JL35|DEOB_YERPY (4)      YLGNCHSSGTVILDQLGEEHMKTGKPIFYTSADSVFQIACHEETFGLDRLYELCEIAREELTDGGYNIGRVIARPFIGDK  220

Consensus                     PGNFQRTGNRRDYAVEPPAPTVLDKL-DEKGGEVVSIGKIADIYAHCGITKKVKATGLDALFDATLE----EMKEA--GD  294
deoB_Bcereus_WP_098782964.1   PGNFTRTPNRHDYALKPFGRTVMNEL-KDSDYDVIAIGKISDIYDGEGVTESLRTKSNMDGMDKLVD----TLNM----D  273
deoB_Rel606                   AGNFQRTGNRHDLAVEPPAPTVLQKLVDEKHGQVVSVGKIADIYANCGITKKVKATGLDALFDATIK----EMKEA--GD  294
sp|Q89A57|DEOB_BUCBP          KEHFRRTGNRRDFSMEPHKITVMEKLIGEKKGRVIAIGKISDIYAGKGISCSMYATGLVNLFNTTIQ----EIKNA--KN  295
sp|Q8K936|DEOB_BUCAP          KSNFNRTGNRRDISMKPLSITFMEKLIQEKQGKVIAIGKISDIYAGVGISKNIKSTGLYELCHTTIH----QMKSA--KN  296
sp|B8D866|DEOB_BUCAT          KLQFQRTGNRRDFSIKPFATTVIKKLIDEKQGQVIAIGKVSDIYGGIGISKNIKSTGLYELCSTTIH----EMKKA--LN  294
sp|P57607|DEOB_BUCAI          KLQFQRTGNRRDFSIKPFATTVIKKLIDEKQGQVIAIGKVSDIYGGIGISKNIKSTGLYELCSTTIH----EMKKA--LN  294
sp|B8D9W4|DEOB_BUCA5          KLQFQRTGNRRDFSIKPFATTVIKKLIDEKQGQVIAIGKVSDIYGGIGISKNIKSTGLYELCSTTIH----EMKKA--LN  294
sp|Q483R0|DEOB_COLP3          ADNFARTGNRRDYSVLPPAPTVLDKI-SQEGTHVISVGKIADIFAHQGIDEKTKATGLNALFDATLD----HINTA--QD  285
sp|Q3ICU9|DEOB_PSET1          NQDFARTSNRRDYSVLPPAPTLLDVL-AKDGGEVISIGKISDIYAHQGITQKHKAPGLINLLKKTNE----LMQSA--PD  285
sp|Q5QXT9|DEOB_IDILO          ADNFERTANRKDYSVLPPKPTVLDKL-QNKGGKVIAIGKISDIFAGQGVSESVKASGLNGLLTATLN----AMEKA--PE  284
sp|Q17YP1|DEOB_HELAH          KDNFKRTSNRKDYAIKPHKKLLFETFIEEKQGEVISIGKIADIYAHVGITQKFKAGSLMELCDVTLE----QVKNA--QN  294
sp|Q9ZK37|DEOB_HELPJ          RENFKRTANRKDYAIKPHKKLLFETFIEEKQGEVISIGKIADIYAHVGITQKFKAGSLMELCDVTLD----QVKNA--PN  294
sp|B6JN18|DEOB_HELP2          RESFKRTANRKDYAIKPHKKLLFETFIEEKQGEVISIGKIADIYAHVGITQKFKAGSLMELCDVTLE----QVKNA--KN  294
sp|P56195|DEOB_HELPY          RESFKRTANRKDYAIKPHKKLLFETFIEEKRGEVISIGKIADIYAHVGITQKFKAGSLMELCDVTLE----QVKNA--KN  294
sp|B5Z8H3|DEOB_HELPG          REDFKRTAKRKDYAIKPHKKLLFEKFIEEKQGEVISIGKIADIYAHVGITQKFKAGSLMELCDVTLD----QVKNA--KN  294
sp|B2UUU2|DEOB_HELPS          REDFKRTAHRKDYAIKPHKKLLFEKFIEEKQGEVISIGKIADIYAHVGITQKFKAGSLMELCDVTLE----QIKNA--KN  294
sp|Q1CS87|DEOB_HELPH          REDFKRTANRKDYAIKPHKKLLFETFIEEKQGEVISIGKIADIYAHVGITQKFKAGSLMELCDVTLE----QIKNA--KN  294
sp|A4YPV1|DEOB_BRASO          ASGFRRTANRRDYAVPPPEPTILDLA-TAERRHVVTIGKIADIFAHRGTGHNLKGDSNDTLFTRMLE----GLDAL--DD  291
sp|B8IJW7|DEOB_METNO          RRGFVRTANRRDFATPPSDTLLDRL-VAAGRPVVSVGKIGDIFAHRSTGSEVKPAGNEACLDAALA----AFAESGLGP  295
sp|B0UPC1|DEOB_METS4          GTGFVRTANRRDFATPPGDTLLDRL-AAAGRPLVSVGKIGDIFAHRHTGTEVKPAGNDACLDAALA----AFADL--GP  293
sp|B1LYJ9|DEOB_METRJ          EAGFRRTGNRRDLAVAPPGRTLLDRA-EAAGRAVVSVGKIGDIFAHRATGREIKPGPNAACLTAGLD----ALATL--PQ  293
sp|B1ZIG8|DEOB_METPB          VDGFRRTSRRKDFSVAPPDGTLLDGL-EAAGRAIVSVGKIGDIFAHRATGREIKPAGNAACLDAALD----AFAEL--PE  293
sp|A9W427|DEOB_METEP          AEGFRRTSHRKDFSVAPPAGTLLDGL-ETAGRAVVSVGKIGDIFAHRATGREIKPAGNAACLDAALD----AFAGL--PE  293
sp|B7KXI3|DEOB_METC4          AEGFRRTSHRKDFSVAPPAGTLLDGL-EAAGRAVVSVGKIGDIFAHRATGREIKPAGNAACLDAALD----AFAGL--PE  293
sp|C3MBH5|DEOB_SINFN          PGNFTRTGHRRDFSVLPPEPTILDRL-EAAGRTVHAIGKIGDIFAHRGVTRLTKANGNMELFDASLT----VVEEA--EE  293
sp|Q92T47|DEOB_RHIME          PGSFTRTGNRRDYSVLPPEPTVLDRL-QEAGRTVHAIGKIGDIFAHQGVTRLTKANGNMALFDASLE----AIEEA--ED  293
sp|A6UET6|DEOB_SINMW          PGSFTRTGNRRDYSVLPPAPTVLDRL-KEAGRTVHAIGKIADIFAHQGVTRLTKANGNMALFDASLA----AIDEA--ED  293
sp|B9JYP7|DEOB_AGRVS          ASNFQRTGNRRDYSVPPPEPTLLDRL-SEAGRTVHAIGKIGDIFAHQGTGRDIKANGNAALMEATLA----VMDEA--AD  293
sp|Q8UJ04|DEOB_AGRFC          VATFERTGNRRDFSVPPPEPTLLDRL-VEAGRKVHAIGKIGDIYAHQGVTRVIKANGNAALMDATLH----AIDEA--EN  293
sp|B9J6V9|DEOB_AGRRK          PANFERTGNRRDFSVLPPEPTLLDRL-VAAERKVHAVGKIGDIFAHQGISRIIKANGNMKLMDATLK----TMDEA--AD  293
sp|Q2KDR4|DEOB_RHIEC          ASTFQRTGNRRDFSVLPPEPTLLDRL-LQHGRHVHAVGKIGDIFAHQGISRVIKATGNEALMDASLS----AIDAA--ED  293
sp|B5ZXI4|DEOB_RHILW          SSTFQRTGNRRDFSVLPPEPTLLDRL-IEHGRHVHAVGKIGDIFAHQGISRLIKANGNEALMDATLA----TIDEA--ED  293
sp|Q1MMV6|DEOB_RHIL3          ASTFQRTGNRRDFSVLPPEPTLLDRL-IEQGRHVHAVGKIGDIFAHQGISRVIKANGNEALMDASLS----AIDAA--ED  293
sp|Q98BG5|DEOB_RHILO          AATFQRTYNRHDYAVPPPEPTLLDRL-TARGSRVIAVGKIGDIFAHRGISEVRKAAGNMAMFDKALG----AMDDA--GD  293
sp|Q11AV9|DEOB_CHESB          AETFERTANRRDYAVPPPEPTLLDRV-EAAGRRVIGIGKIGDIFAHQGVTEVRKAAGNMALFDAALG----AMDDA--RE  290
sp|A5IGR8|DEOB_LEGPC          PGSFKRTGNRKDYATPPPEKTLLDFL-KEDGREVIAIGKIADIYAHQGVTQEIKADGNMALFDATLS----AMKTA--PQ  293
sp|Q5X7B2|DEOB_LEGPA          PGSFKRTGNRKDYATPPPEKTLLDFL-KEDGREVIAIGKIADIYAHQGVTQEIKADGNMALFDATLS----AMKTA--PQ  293
sp|Q5WYR0|DEOB_LEGPL          PGSFKRTGNRKDYATPPPEKTLLDFL-KEDGREVIAIGKIADIYAHQGVTQEIKADGNMALFDATLS----AMKTA--PQ  293
sp|Q5ZXU2|DEOB_LEGPH          PGSFKRTGNRKDYATPPPEKTLLDFL-KEDGREVIAIGKIADIYAHQGVTQEIKADGNMALFDATLS----AMKTA--PQ  293
sp|B2KBT7|DEOB_ELUMP          KGEFKRTKNRHDYAVKPPAPTVLDFL-KENGGNVISIGKINDIYAKQGITKAVKASGLEELWNTTIE----ETKNA--SG  284
sp|Q2SHN3|DEOB_HAHCH          PEGFRRTGARRDFATPPHKPTLLDKL-NNHGKQVIAIGKISDIYAGKGVSRSVKADGLGALVEATLTVMN-ETDEAAAKQ  297
sp|Q1QZC2|DEOB_CHRSD          PETFERTGNRRDYAIEPPTPTVLQKL-HDDGGKVLGVGKIGDIYAHCGVSNVIKAHGHDALFDATLE----ALDEA--GD  285
sp|A1SYK3|DEOB_PSYIN          SSNFMRTGNRHDYSVKPPSPTLLESM-KESGGQVVSIGKISDIFAEQGITKATKANGLEALFDASLN----ELKQA--GD  301
sp|A6VM02|DEOB_ACTSZ          AGAFKRTGNRRDYSVEPPAKTVLQKFIEEKEGMVVSVGKIADIYAHTGISKKVKATGLEELFDKTLE----EVKSA--GD  282
sp|C4LAY9|DEOB_TOLAT          AGEFKRTGNRHDYAVEPPMPTLLDRM-KAAGGDVISIGKIADIYACCGITQQHKATGLDELWDMTLA----QVKTA--AD  291
sp|A4SRU2|DEOB_AERS4          KGNFKRTGNRHDYSVLPPAPTVLDYM-KEAGGQVVSIGKIADIYANKGITKQVKGTGLTELWDRTLE----EVKAA--GD  289
sp|A0KPE2|DEOB_AERHH          KGNFKRTGNRHDYSVLPPAPTVLDYM-KEAGGQVVSIGKIADIYAQQGITKQVKGTGLTELWDRTLE----EVKAA--GD  289
sp|Q7NRT1|DEOB_CHRVO          KGKFARTGNRKDLAVEPPAATVLKKL-ADAGGDVVSIGKIADIYAHVGITHKHKATGFDQLWDATLT----AMDQH--RD  290
sp|Q086F8|DEOB_SHEFN          PADFARTGNRHDYAVLPPAPTVLDKL-KDAGGEVVSIGKISDIYAHSGITQQFKATGLEELFDETLA----QIKRA--GD  291
sp|Q12QG1|DEOB_SHEDO          PSDFSRTGNRRDYAVEPPSKTVLDKL-KDAGGEVVSVGKIADIYAHCGITKKVKASGLEALFDATLE----QIKQA--GD  291
sp|A1S476|DEOB_SHEAM          PSDFARTGNRRDYAVEPPSKTVLDKL-KQAGGEVVSVGKISDIYAHCGITKKVKASGLEDLFDATLE----QIKQA--GD  291
sp|Q8EHK2|DEOB_SHEON          PSDFARTGNRKDYSLEPPAKTVLDKL-KAAGGEVVSVGKIADIYAYCGITKKVKANGLEALFDATLA----EVKSA--GE  291
sp|A0KU09|DEOB_SHESA          PSDFARTGNRKDYSLEPPAKTVLDKL-KAAGGEVVSVGKIADIYAYCGITKKVKANGLEALFDATLA----EVKSA--GE  291
sp|Q0HLE8|DEOB_SHESM          PSDFARTGNRKDYSLEPPAKTVLDKL-KAAGGEVVSVGKIADIYAYCGITKKVKANGLEALFDATLD----EVKSA--GE  291
sp|Q0HXQ2|DEOB_SHESR          PNDFARTGNRKDYSLEPPAKTVLDKL-KAAGGEVVSVGKIADIYAYCGITKKVKANGLEALFDATLA----EVKSA--GE  291
sp|A1RH89|DEOB_SHESW          PSDFARTGNRKDYSLEPPAKTVLDKL-KAAGGEVVSVGKIADIYAYCGITKKVKANGLEDLFDATLA----EVKSA--GD  291
sp|A4Y9A6|DEOB_SHEPC          QSDFARTGNRKDYSLEPPAKTVLDKL-KAAGGEVVSVGKIADIYAYCGITKKVKANGLEDLFDATLA----EVKSA--GD  291
sp|A6WRB6|DEOB_SHEB8          PSDFARTGNRKDYSLEPPAKTVLDKL-KEAGGEVVSVGKIADIYAYCGITKKVKANGLEALFDATLA----EVKSA--GD  291
sp|A9KZ78|DEOB_SHEB9 (2)      PSDFARTGNRKDYSLAPPAKTVLDKL-NEAGGEVVSVGKIADIYAYCGITKKVKANGLEALFDATLA----EVKSA--GD  291
sp|B8E6P6|DEOB_SHEB2          SSDFARTGNRKDYSLAPPAKTVLDKL-NEAGGEVVSVGKIADIYAYCGITKKVKANGLEALFDATLA----EVKSA--GD  291
sp|B0TQ89|DEOB_SHEHH          PSDFARTGNRRDYAVEPPAPTVLDKL-KAAGGEVVSVGKIADIYAHCGITKKVKATGLEELFDATLE----QVKQA--GD  291
sp|A8H726|DEOB_SHEPA          PGSFERTGNRRDYAVEPPAPTVLDKL-KAAGGEVVSVGKIADIYAHCGITKKVKATGLEALFDATLE----QVKLA--GD  291
sp|A8FYQ7|DEOB_SHESH          PSDFARTGNRRDYAVEPPAKTVLDKL-KDSGGEVVSVGKIADIYAHCGITKKIKASGLEALFDATLE----ELKVA--GD  291
sp|A3QGT1|DEOB_SHELP          PSDFARTGNRKDYAVEPPSKTVLDKL-KEAGGEVVSVGKIADIYAHCGITKKVKATGLEALFDATLE----QVKQA--GD  291
sp|B1KRP6|DEOB_SHEWM          PSDFARTGNRRDYAVEPPSKTVLDKM-KAAGGEVISVGKIADIYANCGITQKVKATGLEALFDATLE----QVKAA--GD  291
sp|Q6LUH2|DEOB_PHOPR          KGQFERTGNRRDLSLEPPATTVLQKLVDEKGGDVISIGKISDIYAGCGITKKVKANGIPALFEATLE----QIKQA--GD  293
sp|B6ENG5|DEOB_ALISL          KGQFERTGNRRDLSLEPPAITVLQKLVEEKNGHVHSIGKISDIYAGCGITQKTKATGIPALFDATKE----AITAA--SD  293
sp|B5FAA0|DEOB_ALIFM          KGQFERTGNRRDLSVEPPAATILQKLVDEKGGQVHSIGKISDIYAGCGITKKTKATGIPALFDATKE----AIEQA--GD  293
sp|Q5E7J5|DEOB_ALIF1          KGQFERTGNRRDLSVEPPAATILQKLVDEKGGQVHSIGKISDIYAGCGITKKTKATGIPALFDATKEAIEQAIEQA--GD  297
sp|C3LQC0|DEOB_VIBCM (2)      KGQFARTGNRRDLSVEPPSATVLQKLVEEKQGRVVSIGKIADIYAYCGITDKVKATGIPDLFEATLE----QIKQA--GD  293
sp|Q9KPL9|DEOB_VIBCH          KGQFARTGNRRDLSVEPPSATVLQKLVEEKQGRVVSIGKIADIYAYCGITDKVKATGIPDLFEATLE----QIKQA--GD  293
sp|A7MUW4|DEOB_VIBCB          KGQFERTGNRRDLSVEPPSATVLQKLVEEKQGDVVSIGKIADIYANCGITKKVKATGIPALFEATLE----QIKEA--GD  293
sp|Q7MI40|DEOB_VIBVY (2)      KGQFERTGNRRDLSVEPPSATVLQKLAEEKQGQVVSIGKIADIYANCGITKKVKATGIPALFEATLE----QIKQA--GD  293
sp|Q87M24|DEOB_VIBPA          KGQFERTGNRRDLSVEPPSATVLQKLVEEKQGNVVSIGKIADIYANCGITKKVKATGIPALFEATLE----QIKEA--GD  293
sp|Q2NW04|DEOB_SODGM          AGQFERTGNRHDLAVPPPSATMLQKLVEEKGGTVVSVGKIADIYAQVGISKKVKATGLDALFDATVR----EMDAA--GE  294
sp|B2VH52|DEOB_ERWT9          PGHFERTGNRHDLAVEPPAPTVLKKLVDEQGGEVISVGKIADIYAHVGITKKVKATGLDALFDATVG----EMKSA--PD  294
sp|A7MGA8|DEOB_CROS8          AGNFQRTGNRHDLAVEPPSPVVLKKLVDEKGGHVVSVGKIADIYANMGITKKVKATGLDALFDATIK----EMKEA--GD  294
sp|A4W6A0|DEOB_ENT38          PGEFQRTGNRHDLAVEPPAATVLQKLVDEKDGQVVSVGKIADIYANCGITKKVKATGLDALFDATVK----EMKEA--GD  294
sp|Q327L3|DEOB_SHIDS          AGNFQRTGNRHDLAVEPPAPTVLQKLVDEKHGQVVSVGKIADIYANCGITKKVKATGLDALFDATIK----EMKEA--GD  294
sp|Q1R260|DEOB_ECOUT (3)      AGNFQRTGNRHDLAVEPPAPTVLQKLVDEKHGQVVSVGKIADIYANCGITKKVKATGLDALFDATIK----EMKEA--GD  294
sp|B7UR11|DEOB_ECO27          AGNFQRTGNRHDLAVEPPAPTVLQKLVDEKHGQVVSVGKIADIYANCGITKKVKATGLDALFDATIK----EMKEA--GD  294
sp|Q3YU10|DEOB_SHISS          AGNFQRTGNRHDLAVEPPAPTVLQKLVDEKHGQVVSVGKIADIYANCGITKKVKATGLDALFDATIK----EMKEA--GD  294
sp|Q0SX28|DEOB_SHIF8 (22)     AGNFQRTGNRHDLAVEPPAPTVLQKLVDEKHGQVVSVGKIADIYANCGITKKVKATGLDALFDATIK----EMKEA--GD  294
sp|B5Y275|DEOB_KLEP3          AGNFQRTGNRHDLAVEPPAPTVLQKLVDEKNGHVVSVGKIADIYANCGITKKVKATGLDALFDATIK----EMKEA--GD  294
sp|A9MRA5|DEOB_SALAR          AGNFQRTGNRHDLAVEPPAPTVLQKLVDEKQGHVVSVGKIADIYANCGITKKVKATGLDALFDATIK----EMKDA--GD  294
sp|B5BAJ9|DEOB_SALPK (2)      AGNFQRTGNRHDLAVEPPAPTVLQKLVDEKQGHVVSVGKIADIYANCGITKKVKATGLDALFDATLK----EMKEA--GD  294
sp|C0Q7M5|DEOB_SALPC (2)      AGNFQRTGNRHDLAVEPPAPTVLQKLVDEKQGHVVSVGKIADIYANCGITKKVKATGLDALFDATLK----EMKEA--GD  294
sp|B5R9V1|DEOB_SALG2          AGNFQRTGNRHDLAVEPPAPTVLQKLVDEKQGHVVSVGKIADIYANCGITKKVKATGLDALFDATLK----EMKEA--GD  294
sp|B5FTC7|DEOB_SALDC          AGNFQRTGNRHDLAVEPPAPTVLQKLVDEKQGHVVSVGKIADIYANCGITKKVKATGLDALFDATLK----EMKEA--GD  294
sp|P63924|DEOB_SALTI (8)      AGNFQRTGNRHDLAVEPPAPTVLQKLVDEKQGHVVSVGKIADIYANCGITKKVKATGLDALFDATLK----EMKEA--GD  294
sp|C5BHJ4|DEOB_EDWI9          AGHFQRTGNRHDLAVEPPAPTMLKKLVDEKQGDVVSIGKIADIYANVGITKKVKATGIDALFDATLQ----EMRQA--GN  294
sp|Q6D990|DEOB_PECAS          PGNFERTGNRHDLAVEPPAPTILKKLVDEKGGEVVSVGKIADIYAQVGITKKVKATGIDALFDATLK----EMDSA--GD  294
sp|C6DKL9|DEOB_PECCP          PGHFERTGNRHDLAVEPPAPTILKKMVDEKGGEVVSVGKIADIYAQVGITKKVKATGIDALFDATLK----EMDSA--GD  294
sp|B4EWA2|DEOB_PROMH          PGNFSRTGNRHDLAVEPPAPTMLKKLVDEKQGHVVSIGKIADIYANVGITKKVKATGIDALFDATIE----EMKLA--GD  295
sp|Q7N931|DEOB_PHOLL          AGNFQRTGNRHDLAVEPPAPTILKKLVDEKNGEVVSIGKIADIYANVGITQKVKATGIDALFDATLV----EMEKA--GD  294
sp|A8G9H8|DEOB_SERP5          PGNFQRTGNRHDLAVEPPAPTVLKKLVDEKGGEVVSIGKIADIYANVGITKKVKATGIDALFDATLI----EMEKA--GD  294
sp|A1JJ99|DEOB_YERE8          PGNFQRTGNRHDLAVEPPAPTMLKKLVDEKGGEVVSIGKIADIYAHVGITQKVKATGLDALFDATIE----EMKKA--GD  294
sp|A9R047|DEOB_YERPG          PGHFQRTGNRHDLAVEPPAPTMLKKLVDEKGGEVVSIGKIADIYAQVGITQKVKATGLDALFDATIE----EMKKA--GD  294
sp|A4TQJ1|DEOB_YERPP (4)      PGHFQRTGNRHDLAVEPPAPTMLKKLVDEKGGEVVSIGKIADIYAQVGITQKVKATGLDALFDATIE----EMKKA--GD  294
sp|B1JL35|DEOB_YERPY (4)      PGHFQRTGNRHDLAVEPPAPTMLKKLVDEKGGEVVSIGKIADIYAQVGITQKVKATGLDALFDATIE----EMKKA--GD  294

Consensus                     N-TIVFTNFVDFDSSYGHRRDVAGYAAALEYFDXRLPELLALLKEDDLLILTADHGCDPTWPGTDHTREHIPVLAYGPGV  373
deoB_Bcereus_WP_098782964.1   FTGLSFLNLVDFDALFGHRRDPQGYGEALQEYDARLPEVFAKLKEDDLLLITADHGNDPIHPGTDHTREYVPLLAYSPSM  353
deoB_Rel606                   N-TIVFTNFVDFDSSWGHRRDVAGYAAGLELFDRRLPELMSLLRDDDILILTADHGCDPTWTGTDHTREHIPVLVYGPKV  373
sp|Q89A57|DEOB_BUCBP          N-TIVFVNFVDFDSLWGHRRDVSGYAKDLEWFDYNLPKLLKLVHNEDLLIITADHGCDPTWIGTDHTRENVPILIYQRSM  374
sp|Q8K936|DEOB_BUCAP          N-TVIFTNFVDFDSVWGHRRDVSGYAKGLEFFDSKLSEMIDLVKEGDLFILTADHGCDPTWKGTDHTRENVPILIYSPGE  375
sp|B8D866|DEOB_BUCAT          N-TIVFTNLVDFDSNWGHRRDVSGYAKGLELFDSRLSEIISLVQKNDLLILTADHGCDPTWIGTDHTRENVPVLIYSPGI  373
sp|P57607|DEOB_BUCAI          N-TIVFTNLVDFDSNWGHRRDVSGYAKGLELFDSRLSEIISLVQKNDLLILTADHGCDPTWIGTDHTRENVPVLIYSPGI  373
sp|B8D9W4|DEOB_BUCA5          N-TIVFTNLVDFDSNWGHRRDVSGYAKGLELFDSRLSEIISLVQKNDLLILTADHGCDPTWIGTDHTRENVPVLIYSPGI  373
sp|Q483R0|DEOB_COLP3          N-SLIFTNLVNFDQDFGHRRDAIGYAKELEALDVRIPELFHAMSAEDVLFLTADHGCDPTWPGTEHTREYVPIIAYHHQI  364
sp|Q3ICU9|DEOB_PSET1          H-SLIFTNLVDFDEMFGHRRNPVGYAKALKEFDDYLPTILNALKADDLLIITADHGCDPTFPGSEHTREYVPVIAYQPGM  364
sp|Q5QXT9|DEOB_IDILO          N-TLIFTNLVDFDTLYGHRRDIDGYARELETFDQWLPVIESAMTPDDVLVLTADHGCDPTWQGTDHTRELIPLLLSGQNV  363
sp|Q17YP1|DEOB_HELAH          N-SLIFTNFVHFDSDYGHRRDVIGYANALEYFDAYLKEILESLRENDLLILCADHGCDPSFKGTDHTREYIPVLMYHKDL  373
sp|Q9ZK37|DEOB_HELPJ          N-SLIFTNFVHFDSDYGHRRDVSGYANALEYFDARLKEVLDNLRENDLLILCADHGCDPSFKGTDHTREYIPVLFYHKDL  373
sp|B6JN18|DEOB_HELP2          N-SLIFTNFVHFDSDYGHRCDISGYANALEYFDTRLKEVLENLRENDLLILCADHGCDPSFKGTDHTREYIPVLFYHKDL  373
sp|P56195|DEOB_HELPY          N-SLIFTNFVHFDSDYGHRRDISGYANALEYFDARLKEVLENLRENDLLILCADHGCDPSFKGTDHTREYIPVLFYHKDL  373
sp|B5Z8H3|DEOB_HELPG          N-SLIFTNFVHFDSDYGHRRDISGYANALEYFDTRLKEVLENLKENDLLILCADHGCDPSFKGTDHTREYIPVLFYHKDL  373
sp|B2UUU2|DEOB_HELPS          N-SLIFTNFVHFDSDYGHRRDISGYANALEYFDARLKEVLENLRENDLLILCADHGCDPSFKGTDHTREYIPVLFYHKDL  373
sp|Q1CS87|DEOB_HELPH          N-SLIFTNFVHFDSDYGHRRDISGYANALEYFDARLKEILDNLKENDLLILCADHGCDPSFKGTDHTREYIPVLFYHKDL  373
sp|A4YPV1|DEOB_BRASO          G-GLLFANFIDFDTLYGHRRDVAGYASALEAFDARLPELLDALRDDDLLVITADHGCDPSWSGSDHTRERVPVLIVN-GR  369
sp|B8IJW7|DEOB_METNO          G-GLVFVNLVDFDTEYGHRRDVPGYAAALERFDARIPEIRAGLAPGDLCVITADHGNDPTWSGTDHTREQVPVLSFGPGL  374
sp|B0UPC1|DEOB_METS4          G-GLVFANLVDFDTEYGHRRDVPGYAAALERFDARLPEIRAALRPGDLCLVTADHGNDPTWTGTDHTREQVPVLAFGPGQ  372
sp|B1LYJ9|DEOB_METRJ          G-GLIFVNLVDFDTEHGHRRDVPGYAAELEAFDARIPEILAALAPGDLAVITADHGNDPTWTGTDHTREQVPVLAFGPGV  372
sp|B1ZIG8|DEOB_METPB          G-GFVFLNLVDFDTEHGHRRDVPGYAAELEAFDARIPEIHAVLRPGDLCVITADHGNDPTWTGTEHTREQVPVLAFGPGL  372
sp|A9W427|DEOB_METEP          G-GFVFLNLVDFDTEHGHRRDVPGYAAELEAFDRRLPEIQAVLKPGDLCVITADHGNDPTWTGTEHTREQVPVLAFGPGL  372
sp|B7KXI3|DEOB_METC4          G-GFVFLNLVDFDTEHGHRRDVPGYAAELEAFDARLPEIQAVLKPGDLCVITADHGNDPTWTGTEHTREQVPVLAFGPGL  372
sp|C3MBH5|DEOB_SINFN          G-DLVFTNFVDFDMLYGHRRDVPGYAAALEAFDARLPDLDRRLRPGDMVILTADHGCDPTWRGTDHTRERVPVLMFGPSL  372
sp|Q92T47|DEOB_RHIME          G-ALVFTNFVDFDMLYGHRRDVSGYAAALEAFDARLPDLDRRLKPGDMVILTADHGCDPTWRGTDHTRERVPVLMFGPTL  372
sp|A6UET6|DEOB_SINMW          G-ALIFTNFVDFDMLYGHRRDVAGYAAALEAFDARLPDLDRRLKPGDMVILTADHGCDPTWRGTDHTRERVPVLMFGPTL  372
sp|B9JYP7|DEOB_AGRVS          G-DLVFTNFVDFDMLYGHRRDVPGYAAALEAFDLWLPDVYRKLIPGDMVILTADHGCDPTWRGTDHTRERVPIMAFGPGI  372
sp|Q8UJ04|DEOB_AGRFC          G-DLVFTNFVDFDMLYGHRRDVAGYAAALEAFDARIPEIHRKMAPGDIALLTADHGCDPTWRGTDHTRERVPIMAFGPGI  372
sp|B9J6V9|DEOB_AGRRK          G-DLVFTNFVDFDMVYGHRRDVAGYAAALEAFDARLPEVHRKLKPGDLLILTADHGCDPTWRGTDHTRERVPIIAYELGI  372
sp|Q2KDR4|DEOB_RHIEC          G-DLVFTNFVDFDMNYGHRRDVPGYAAALEAFDARLPEVHKKLKPGDLVVLTADHGCDPTWRGTDHTRERVPIIAFGPGI  372
sp|B5ZXI4|DEOB_RHILW          G-DLVFTNFVDFDMVYGHRRDVPGYAAALEAFDARLAEVHKKLKPGDLVVLTADHGCDPTWRGTDHTRERVPVIAYGPGI  372
sp|Q1MMV6|DEOB_RHIL3          G-DLVFTNFVDFDMIYGHRRDVPGYAAALEAFDARLPDVHKKLKPGDLVVLTADHGCDPTWRGTDHTRERVPVIAYGPGI  372
sp|Q98BG5|DEOB_RHILO          G-DLVFANFVDFDTEFGHRRDVAGYAAALEAFDRRLPEAFARLRQGDLLILTADHGNDPTWRGTDHTRERIPVIGMGPGL  372
sp|Q11AV9|DEOB_CHESB          G-DLVFANFVDFDSLYGHRRDVAGYAAALEAFDRRLPEALGKLRAGDLLILTADHGCDPTWRGTDHTRECVPILGAGPGL  369
sp|A5IGR8|DEOB_LEGPC          G-SLVFTNFVDFDSSYGHRRDVAGYAHALEQFDARLPELEVLLQPNDMVFIAADHGCDPTFPGSDHTREHIPVLMFGPQV  372
sp|Q5X7B2|DEOB_LEGPA          G-SLVFTNFVDFDSSYGHRRDVAGYAHALEQFDVRLPELEALLQPDDMVFIAADHGCDPTFPGSDHTREHIPVLMFGPQV  372
sp|Q5WYR0|DEOB_LEGPL          G-SLVFTNFVDFDSSYGHRRDIAGYAHALEQFDARLPELEVLLQPNDMVFIAADHGCDPTFPGSDHTREHIPVLVFGPQV  372
sp|Q5ZXU2|DEOB_LEGPH          G-SLVFTNFVDFDSSYGHRRDIAGYAHALEQFDARLPELDALLQPDDMVFIAADHGCDPTFPGSDHTREHIPVLVFGPQV  372
sp|B2KBT7|DEOB_ELUMP          N-SIIFTNFVDFDMVWGHRRDFKGYAGGLEYFDSRLPELANILQEGDLVFITADHGCDPSYKGTDHTRENVPAIMFGKNV  363
sp|Q2SHN3|DEOB_HAHCH          D-TLIFTNLVDFDMLYGHRRDVTGYAKALEDFDAMLPDLLGAMMDDDLLILTADHGCDPTWPGSDHTREHIPILMYGKRA  376
sp|Q1QZC2|DEOB_CHRSD          R-TLVMTNFVDFDTLYGHRRDPDGYAEALEAFDRRLPEVLAKLRPDDLLILTADHGNDPTWTGTDHTREQVPVLASGAGL  364
sp|A1SYK3|DEOB_PSYIN          Q-TIIFTNFVNFDADFGHRRNLPGYAAALEYFDKRLPEMLALMTEDDLLVLTADHGCDPTWKGTDHTREHIPAIFYGHAV  380
sp|A6VM02|DEOB_ACTSZ          N-TIVFTNFVNFDADFGHRRDVTGYAKGLEYFDRRLPELLRLMKDDDLLIITADHGCDPTWQGSDHTREHIPVLMYGAQV  361
sp|C4LAY9|DEOB_TOLAT          Q-SIIFTNFVDFDSSYGHRRDVAGYAAALEYFDSRLPELLALLLPGDRVVLTADHGCDPIWTGTDHTREHVPVIFYGDTV  370
sp|A4SRU2|DEOB_AERS4          N-TIVFTNFVDFDSSYGHRRDVKGYADALEYFDSRLPELFEILQDGDVVVLTADHGCDPTWGGTDHTREYIPVLFFGKPV  368
sp|A0KPE2|DEOB_AERHH          Q-TIVFTNFVDFDSSYGHRRDVKGYADALEYFDSRLPELFELLEDGDVVVLTADHGCDPTWGGTDHTREYIPVLFYGKPV  368
```

```
sp|Q7NRT1|DEOB_CHRVO        KPAIIMANFVDFDSSYGHRRNTAGYAAALEEFDARLPEVMAALGDDDILILSADHGCDPTWPGTDHTREHIPVLCYGKQV  370
sp|Q086F8|DEOB_SHEFN        N-TIVFTNFVDFDSHYGHRRDTAGYAKALEYFDSRLPELLAILQPEDLVIFTADHGCDPTWVGTEHTRERVPVLAYGAGL  370
sp|Q12QG1|DEOB_SHEDO        R-TIVFTNFVDFDSHYGHRRDVAGYAKALEYFDERLPELLALLEQDDLLLLTADHGCDPTWPGSDHTRERVPVLALGAGL  370
sp|A1S476|DEOB_SHEAM        N-TIVFTNFVDFDSHYGHRRDVSGYAKALEYFDARLPELMALLEEGDLLLLTADHGCDPTWQGTDHTREFVPVLAYGAGL  370
sp|Q8EHK2|DEOB_SHEON        N-TIVFTNFVDFDSHYGHRRDVAGYAKGLEYFDARLPEMLALLDEDDLLILTADHGCDPTWQGTDHTREYVPVLAYGAGL  370
sp|A0KU09|DEOB_SHESA        N-TIVFTNFVDFDSHYGHRRDVAGYAKGLEYFDSRLPEMLALLDEDDLLILTADHGCDPTWQGTDHTREYVPVLAYGAGL  370
sp|Q0HLE8|DEOB_SHESM        N-TIVFTNFVDFDSHYGHRRDVAGYAKGLEYFDSRLPEMLALLDEDDLLILTADHGCDPTWQGTDHTREYVPVLAYGAGL  370
sp|Q0HXQ2|DEOB_SHESR        N-TIVFTNFVDFDSHYGHRRDVAGYAKGLEYFDSRLPEMLSLLDEDDLLILTADHGCDPTWQGTDHTREYVPVLAYGAGL  370
sp|A1RH89|DEOB_SHESW        N-TIVFTNFVDFDSHYGHRRDVAGYAKGLEYFDARLPEMLALLGEDDLLILTADHGCDPTWQGTDHTREYVPVLAYGAGL  370
sp|A4Y9A6|DEOB_SHEPC        N-TIVFTNFVDFDSHYGHRRDVAGYAKGLEYFDARLPEMLALLGEDDLLILTADHGCDPTWQGTDHTREYVPVLAYGAGL  370
sp|A6WRB6|DEOB_SHEB8        N-TIVFTNFVDFDSHYGHRRDVAGYAKGLEYFDARLPEMLALLGEDDLLILTADHGCDPTWQGTDHTREYVPVLAFGAGL  370
sp|A9KZ78|DEOB_SHEB9 (2)    N-TIVFTNFVDFDSHYGHRRDVAGYAKGLEYFDARLPEMLALLGEDDLLILTADHGCDPTWQGTDHTREYVPVLAFGAGL  370
sp|B8E6P6|DEOB_SHEB2        N-TIVFTNFVDFDSHYGHRRDVAGYAKGLEYFDARLPEMLALLGEDDLLILTADHGCDPTWQGTDHTREYVPVLAFGAGL  370
sp|B0TQ89|DEOB_SHEHH        N-TIVFTNFVDFDSHFGHRRDVAGYARSLEYFDSRLPEILALLDEDDFLLLTADHGCDPTWPGSDHTRERVPVLAYGAGL  370
sp|A8H726|DEOB_SHEPA        N-TIVFTNFVDFDSHYGHRRDVAGYARSLEYFDSRLPEILALLDEDDFLLLTADHGCDPTWPGSDHTRERVPVLAYGAGL  370
sp|A8FYQ7|DEOB_SHESH        N-TIVFTNFVDFDSHFGHRRDVAGYARSLEYFDSRLPELMALMGEDDLLLLTADHGCDPTWPGTDHTRERVPVLAYGAGL  370
sp|A3QGT1|DEOB_SHELP        R-TIVFTNFVDFDSHYGHRRDIAGYARALEYFDSRLPEMLALLGEEDLLLLTADHGCDPTWQGTDHTRERVPVLAYGAGL  370
sp|B1KRP6|DEOB_SHEWM        K-SIVFTNFVDFDSHYGHRRDIAGYAKALEYFDSRLPEIFEILGEDDLLLLTADHGCDPSWKGTDHTRERVPVLAYGAGL  370
sp|Q6LUH2|DEOB_PHOPR        N-SLVFTNFVDFDSAYGHRRNVAGYAAALEYFDKRLPEVLELLQEDDVLILTADHGCDPTWEGTDHTREHIPVIVTGPKI  372
sp|B6ENG5|DEOB_ALISL        N-TIVFTNFVDFDSAYGHRRDVAGYAAALEYFDGRLPEIMEMLQEDDILILTADHGCDPTWPGTDHTREHIPVLVYGHKV  372
sp|B5FAA0|DEOB_ALIFM        N-TIVFTNFVDFDSAYGHRRDVAGYAAALEYFDGRLPEIMDMLQEDDILILTADHGCDPTWPGTDHTREHIPVLVYGHKV  372
sp|Q5E7J5|DEOB_ALIF1        N-TIVFTNFVDFDSAYGHRRDVAGYAAALEYFDGRLPEIMDMLQEDDILILTADHGCDPTWPGTDHTREHIPVLVYGHKV  376
sp|C3LQC0|DEOB_VIBCM (2)    N-TIVFTNFVDFDSAYGHRRDVAGYAAALEYFDKRLPEVLALMQEDDILILTADHGCDPTWPGTDHTREHIPVLVYGKKV  372
sp|Q9KPL9|DEOB_VIBCH        N-TIVFTNFVDFDSAYGHRRDVAGYAAALEYFDKRLPEVLELMQEDDILILTADHGCDPTWPGTDHTREHIPVLVYGKKV  372
sp|A7MUW4|DEOB_VIBCB        N-TIVFTNFVDFDSAYGHRRDVAGYAAALEYFDGRINEVLEIMEEDDVLILTADHGCDPTWPGTDHTREHIPVIVYGKRV  372
sp|Q7MI40|DEOB_VIBVY (2)    N-TIVFTNFVDFDSAYGHRRDVAGYAAALEYFDGRIHEVMELMQEDDILILTADHGCDPTWPGTDHTREHIPVLVYGQKV  372
sp|Q87M24|DEOB_VIBPA        N-TIVFTNFVDFDSAYGHRRDVAGYAAALEYFDGRINEVLELMGEDDVLILTADHGCDPTWPGTDHTREHIPVLVYGQKV  372
sp|Q2NW04|DEOB_SODGM        N-TLVFTNFVDFDSAYGHRRDVAGYAAALELFDRRLPELMSRVQGNDILILTADHGCDPTWHGTDHTREHVPVLIYGPTV  373
sp|B2VH52|DEOB_ERWT9        N-SIVFTNFVDFDSAWGHRRDIPGYAAGLELFDRRLPELMALVKEGDILILTADHGCDPSWPGTEHTREHIPVLIFGPGV  373
sp|A7MGA8|DEOB_CROS8        N-TIVFTNFVDFDSSWGHRRDVAGYAAGLELFDRRLPELLELVGEDDIIIFTADHGCDPTWKGTDHTREHIPVLVYGPKV  373
sp|A4W6A0|DEOB_ENT38        K-TIVFTNFVDFDSSWGHRRDIAGYAAGLELFDRRLPELMELVGEDDILILTADHGCDPSWTGTDHTREHIPVLVYGPKV  373
sp|Q327L3|DEOB_SHIDS        N-TIVFTNFVDFDSSWGHRRDVAGYAAGLELFDRRLPELMSLLRDDDILILTADHGCDPTWTGTDHTREHIPVLVYGPKV  373
sp|Q1R260|DEOB_ECOUT (3)    N-TIVFTNFVDFDSSWGHRRDVAGYAAGLELFDRRLPELMSLLRDDDILILTADHGCDPTWTGTDHTREHIPVLVYGPKV  373
sp|B7UR11|DEOB_ECO27        N-TIVFTNFVDFDSSWGHRRDVAGYAAGLELFDRRLPELMSLLRDDDILILTADHGCDPTWTGTDHTREHIPVLVYGPKV  373
sp|Q3YU10|DEOB_SHISS        N-TIVFTNFVDFDSSWGHRRDVAGYAVGLELFDRRLPELMSLLRDDDILILTADHGCDPTWTGTDHTREHIPVLVYGPKV  373
sp|Q0SX28|DEOB_SHIF8 (22)   N-TIVFTNFVDFDSSWGHRRDVAGYAAGLELFDRRLPELMSLLRDDDILILTADHGCDPTWTGTDHTREHIPVLVYGPKV  373
sp|B5Y275|DEOB_KLEP3        E-TIVFTNFVDFDSSWGHRRDVAGYAAGLELFDRRLPELMELVGEDDILILTADHGCDPTWTGTDHTREHIPVLVYGPKV  373
sp|A9MRA5|DEOB_SALAR        K-TIVFTNFVDFDSSWGHRRDIAGYASGLELFDRRLPELMALVGEDDILILTADHGCDPSWTGTDHTREHIPVLIYGPKV  373
sp|B5BAJ9|DEOB_SALPK (2)    K-TIVFTNFVDFDSSWGHRRDIAGYAAGLELFDRRLPELMELVGEDDILILTADHGCDPSWTGTDHTREHIPVLIYGPKV  373
sp|C0Q7M5|DEOB_SALPC (2)    K-TIVFTNFVDFDSSWGHRRDIAGYAAGLELFDRRLPELMELVGEDDILILTADHGCDPSWTGTDHTREHIPVLIYGPKV  373
sp|B5R9V1|DEOB_SALG2        K-TIVFTNFVDFDSSWGHRRDIAGYAAGLELFDRRLPELMELVGEDDILILTADHGCDPSWTGTDHTREHIPVLIYGPKV  373
sp|B5FTC7|DEOB_SALDC        K-TIVFTNFVDFDSSWGHRRDIAGYAAGLELFDRRLPELMELVGEDDILILTADHGCDPSWTGTDHTREHIPVLIYGPKV  373
sp|P63924|DEOB_SALTI (8)    K-TIVFTNFVDFDSSWGHRRDIAGYAAGLELFDRRLPELMELVGEDDILILTADHGCDPSWTGTDHTREHIPVLIYGPKV  373
sp|C5BHJ4|DEOB_EDWI9        D-TIVFTNFVDFDSSYGHRRDVAGYAAALELFDRRLPEMLALVKEDDILILTADHGCDPTWHGSDHTREHIPVLVYGPKV  373
sp|Q6D990|DEOB_PECAS        N-TIVFTNFVDFDSAYGHRRDIPGYAAALELFDRRLPELMSRVTGDDILILTADHGCDPSWHGTDHTRENVPVLIYGPKV  373
sp|C6DKL9|DEOB_PECCP        N-TIVFTNFVDFDSAYGHRRDIPGYAAALELFDRRLPEMLSRVKGDDILILTADHGCDPSWHGTDHTRENVPVLIYGPNV  373
sp|B4EWA2|DEOB_PROMH        N-TIVFTNFVDFDSSYGHRRDVVGYGEALELFDRRLPELMELVKEDDILILTADHGCDPTWQGSDHTREHIPVLVYGPKV  374
sp|Q7N931|DEOB_PHOLL        N-TIVFTNFVDFDSSYGHRRDVPGYAAALELFDRRLPEMLKRVKNDDILILTADHGCDPTWSGTDHTREHIPVLIYGPKV  373
sp|A8G9H8|DEOB_SERP5        N-TIVFTNFVDFDSSYGHRRDVAGYAAALELFDRRLPELLKLVKDEDIIIFTADHGCDPTWPGTDHTREHIPVLVYGPKV  373
sp|A1JJ99|DEOB_YERE8        N-TIVFTNFVDFDSSYGHRRDVAGYAAALELFDRRLPELMALVKEDDILLLTADHGCDPTWPGTDHTREHIPVLVYGPKV  373
sp|A9R047|DEOB_YERPG        N-TIVFTNFVDFDSSYGHRRDVAGYAAALELFDRRLPELMALIKEDDILILTADHGCDPTWPGTDHTREHIPVLVYGPKV  373
sp|A4TQJ1|DEOB_YERPP (4)    N-TIVFTNFVDFDSSYGHRRDVAGYAAALELFDRRLPELMALIKEDDILILTADHGCDPTWPGTDHTREHIPVLVYGPKV  373
sp|B1JL35|DEOB_YERPY (4)    N-TIVFTNFVDFDSSYGHRRDVAGYAAALELFDRRLPELMALVKEDDILILTADHGCDPTWPGTDHTREHIPVLVYGPKV  373

Consensus                   KPG--SLGRRET----FADIGQTIASYFGL-SPMDYGKSFLNFKGQPXSSHASTTGD  423
deoB_Bcereus_WP_098782964.1 KEGGQELPLRQT----FADIGATVAENFGV-KMPEYGKSFLNELKK-----------  394
deoB_Rel606                 KPG--SLGHRET----FADIGQTLAKYFGT-SDMEYGKAMF----------------  407
sp|Q89A57|DEOB_BUCBP        ESK--NFGYRET----FSDIGQTLAKYFNL-STMSYGTSIF----------------  408
sp|Q8K936|DEOB_BUCAP        EIK--FLGHRDT----FSDIGQTIAKYFSL-SDMKYGKSMF----------------  409
sp|B8D866|DEOB_BUCAT        KKN--FLGHRKT----FADIGQTIAKYFLL-TDMSYGQNML----------------  407
sp|P57607|DEOB_BUCAI        KKN--FLGHRKT----FADIGQTIAKYFLL-TDMSYGQNML----------------  407
sp|B8D9W4|DEOB_BUCA5        KKN--FLGHRKT----FADIGQTIAKYFLL-TDMSYGQNML----------------  407
sp|Q483R0|DEOB_COLP3        DSV--NLGNRKT----FADLGQSVAELFNV-EAMDYGTSFLSEIYSK----------  404
sp|Q3ICU9|DEOB_PSET1        TDI--PLGERNS----FADIGQTLAQWFNL-PALEYGDGFIDKLTTSK---------  405
sp|Q5QXT9|DEOB_IDILO        QAG--NRGKRHS----FADLGQTLCRLFDL-PAMEEGKAIKLS--------------  399
sp|Q17YP1|DEOB_HELAH        QPT--FLGRSET----FADIGQSIAHFLGL-SPLDYGKNLLNFKGQP----------  413
sp|Q9ZK37|DEOB_HELPJ        QPA--FLGKSET----FADIGQSIAYFLGL-SPLDYGKNLLNFKGQP----------  413
sp|B6JN18|DEOB_HELP2        QPA--FLGKSDS----FADIGQSIAYFLGL-SPLDYGKNLLNFKGQP----------  413
sp|P56195|DEOB_HELPY        QPA--FLGKSES----FADIGQSIAHFLGL-SPLDYGKNLLNFKGQP----------  413
sp|B5Z8H3|DEOB_HELPG        QPA--FLGKSES----FADIGQSIAYFLGL-SPLDYGKNLKFKGQP-----------  413
sp|B2UUU2|DEOB_HELPS        QPA--FLGKSES----FADIGQSIAYFLGL-SPLDYGKNLLNFKGQS----------  413
sp|Q1CS87|DEOB_HELPH        QPA--FLGKSES----FADIGQSIAYFLGL-SPLDYGKNLLNFKGQP----------  413
sp|A4YPV1|DEOB_BRASO        SAG--SIGARTS----FADTGATIAQHLQL-PPTRHGNSFWPNGA------------  407
sp|B8IJW7|DEOB_METNO        RPG--PLGRRDT----FADIGAALAAHLGL-PP-SCGRSWL----------------  407
sp|B0UPC1|DEOB_METS4        RPG--PIGRREG----LADIGATVAAHLGL--ALPAGRSWL----------------  405
sp|B1LYJ9|DEOB_METRJ        AAR--PIGRRET----FADIGATVAAHLGL-AWDGAGTPFL----------------  406
sp|B1ZIG8|DEOB_METPB        ATG--AIGRRET----FADIGASVAAHLGL-PPLGAGRAWW----------------  406
sp|A9W427|DEOB_METEP        APR--ALGRRES----FADMGASVAAHLGL-PPLGAGQAWW----------------  406
sp|B7KXI3|DEOB_METC4        TPR--ALGRRES----FADMGASVAAHLGL-PPLGAGQAW-----------------  405
sp|C3MBH5|DEOB_SINFN        RSR--SFGIANS----FAHIGETAARHLGI-APGPHGRSLL----------------  406
sp|Q92T47|DEOB_RHIME        RSR--SFGIADS----FAHIGETVARHLGI-GVGPHGRSLI----------------  406
sp|A6UET6|DEOB_SINMW        RSR--SVGIVGS----FAHIGETVASHLGI-DPGPHGRSLI----------------  406
sp|B9JYP7|DEOB_AGRVS        RAR--SIGIRDT----YADIGETIAAHLGI-APGRHGMSFL----------------  406
sp|Q8UJ04|DEOB_AGRFC        RSR--DVGIRSS----YADIGESIAHHLGI-EAGSHGRSFI----------------  406
sp|B9J6V9|DEOB_AGRRK        RSR--PIGIRQT----YADIGETVAHHLGI-AAGPHGRSFL----------------  406
sp|Q2KDR4|DEOB_RHIEC        RSR--SIGVRRT----YADIGESIARHLGI-PAGPHGRSFL----------------  406
sp|B5ZXI4|DEOB_RHILW        RSR--SIGVRRS----YADIGESIARHLGI-PAGPHGRSFL----------------  406
sp|Q1MMV6|DEOB_RHIL3        RSR--SIGVRRG----YADIGESIARHLGI-PAGPHGRSFL----------------  406
sp|Q98BG5|DEOB_RHILO        AGG--DIGLRTT----FADIGETVAEHLGL-APGRHGTSFHAMIGGHA---------  413
sp|Q11AV9|DEOB_CHESB        AKG--SIGRRRS----YADIGETIASHLAL-PAGRHGVSFLNALQHA----------  409
sp|A5IGR8|DEOB_LEGPC        NSK--FIGRRDC----FADIGQSIAEHLQLSSPLAHGVSFL----------------  407
sp|Q5X7B2|DEOB_LEGPA        NSK--FIGRRDC----FADIGQSIAEHLQLSSPLTHGVSFL----------------  407
sp|Q5WYR0|DEOB_LEGPL        NSK--FIGRRDC----FADIGQSIAEHLQLSSPLTHGVSFL----------------  407
sp|Q5ZXU2|DEOB_LEGPH        NSK--FIGRRDC----FADIGQSIAEHLQLSSPLAHGVSFL----------------  407
sp|B2KBT7|DEOB_ELUMP        KNG--FIGGRET----YSDLGQTVAEYLGI-TKLNNGTSFL----------------  397
sp|Q2SHN3|DEOB_HAHCH        PHG--FIGARKT----FADIGQTLAEYFQL-DRLDFGESFLVADK------------  414
sp|Q1QZC2|DEOB_CHRSD        APR--PLGHDDGHLVTFADIGQSLATHFGL-SPMAHGSDFLDVAPTGPSSHASTTGD  418
sp|A1SYK3|DEOB_PSYIN        KPG--PIGLRET----FADIGQSIADFHKL-PALAYGKSIFS---------------  415
sp|A6VM02|DEOB_ACTSZ        PAR--FLGARET----FADIGQTVAKYLGV-SPMEYGTAII----------------  395
sp|C4LAY9|DEOB_TOLAT        QPQ--DLGMRDT----FADIGQTIAAYHGL-PVLDYGSNCLPEHH------------  408
sp|A4SRU2|DEOB_AERS4        KAG--SVGRRET----FADIGQSIAAYHGL-PKLEYGTSFL----------------  402
sp|A0KPE2|DEOB_AERHH        KAG--SVGRRET----FADIGQSIAAYHGL-PKLAYGTSFL----------------  402
sp|Q7NRT1|DEOB_CHRVO        KAG--PIGERAT----FADIGQSVAAHLGL-PRMDYGTSFLD---------------  405
sp|Q086F8|DEOB_SHEFN        SAG--SLGRRKS----FADIGQSIASYFKL-EPMNYGESFIN---------------  405
sp|Q12QG1|DEOB_SHEDO        AAG--SLGLRNS----FADMGQSIASYFEL-EPMEYGESFIQ---------------  405
sp|A1S476|DEOB_SHEAM        EAG--SLGKRGS----FADIGQSIASYFGL-AAMEYGESFMPAR-------------  407
sp|Q8EHK2|DEOB_SHEON        KAG--SLGRRNS----FADIGQSIASYFKL-EPMEYGESFI----------------  404
sp|A0KU09|DEOB_SHESA        KAG--SLGRRNS----FADIGQSIASYFKL-EPMEYGESFI----------------  404
sp|Q0HLE8|DEOB_SHESM        KAG--SLGRRNS----FADIGQSIASYFKL-EPMEYGESFI----------------  404
sp|Q0HXQ2|DEOB_SHESR        KAG--SLGRRNS----FADIGQSIASYFKL-EPMEYGESFI----------------  404
sp|A1RH89|DEOB_SHESW        KAG--SLGRRNS----FADIGQSIASHFKL-EPMAYGESFI----------------  404
sp|A4Y9A6|DEOB_SHEPC        KAG--SLGRRNS----FADIGQSIASHFKL-EPMAYGESFI----------------  404
sp|A6WRB6|DEOB_SHEB8        KAG--SLGRRKS----FADIGQSIASHFKL-EPMAYGESFL----------------  404
sp|A9KZ78|DEOB_SHEB9 (2)    KAG--SLGRRKS----FADIGQSIASHFKL-EPMAYGESFL----------------  404
sp|B8E6P6|DEOB_SHEB2        KAG--SLGRRKS----FADIGQSIASHFKL-EPMAYGESFL----------------  404
sp|B0TQ89|DEOB_SHEHH        EPG--SLGLRNS----FADMGQSIASYHKL-EPMEYGESFVRS--------------  406
sp|A8H726|DEOB_SHEPA        EAG--SLGLRSS----FADMGQSIASYFKL-EPMEYGESFIR---------------  405
sp|A8FYQ7|DEOB_SHESH        EAG--SLGRRKS----FADMGQSIASYFKL-EPMEYGESFIK---------------  405
sp|A3QGT1|DEOB_SHELP        APG--SLGRRNS----FADIGQSIASYFGL-EPMEYGESFVA---------------  405
sp|B1KRP6|DEOB_SHEWM        KAG--SLGRRNS----FADIGQSIASYFKL-EPMEYGESFIK---------------  405
sp|Q6LUH2|DEOB_PHOPR        PAG--SLGRRET----FADIGQSLAEYYGT-SDMEYGKSFL----------------  406
sp|B6ENG5|DEOB_ALISL        PAG--SLGLRDT----FADIGQTLAEYFEI-SDMEYGKSFL----------------  406
sp|B5FAA0|DEOB_ALIFM        PAG--SLGRRDT----FADIGQTLAEYFET-SDMEYGKSFL----------------  406
sp|Q5E7J5|DEOB_ALIF1        PAG--SLGRRDT----FADIGQTLAEYFET-SDMEYGKSFL----------------  410
sp|C3LQC0|DEOB_VIBCM (2)    APG--SLGRRDT----FADIGQTLASYFGT-SPMDYGKNFL----------------  406
sp|Q9KPL9|DEOB_VIBCH        APG--SLGRRDT----FADIGQTLASYFGT-SPMDYGKNFL----------------  406
sp|A7MUW4|DEOB_VIBCB        PAG--SLGRRDS----FADIGQTLATYFGT-SPMDYGKNFL----------------  406
sp|Q7MI40|DEOB_VIBVY (2)    PAG--SLGRRET----FADIGQTLASYFGT-SPMDYGKNFL----------------  406
sp|Q87M24|DEOB_VIBPA        PAG--SLGRRET----FADIGQTLASYFGT-SPMDYGKNFL----------------  406
sp|Q2NW04|DEOB_SODGM        NPG--FYGHRTT----FADIGQTVARYFGL-SPMDYGKAIM----------------  407
sp|B2VH52|DEOB_ERWT9        RPG--DYGYRDT----FADIGQTLAHYFGL-SPMAYGKPFF----------------  407
sp|A7MGA8|DEOB_CROS8        KPG--SLGHRDT----FADIGQTVAKYFGL-SDMEYGKALF----------------  407
sp|A4W6A0|DEOB_ENT38        KPG--SLGHRET----FADIGQTIASYFGT-SPMDYGKNML----------------  407
sp|Q327L3|DEOB_SHIDS        KPG--SLGHRET----FADIGQTLAKYFGT-SDMEYGKAMF----------------  407
sp|Q1R260|DEOB_ECOUT (3)    KPG--SLGHRET----FADIGQTLAKYFGT-SDMEYGKAMF----------------  407
sp|B7UR11|DEOB_ECO27        KPG--SLGHRET----FADIGQTLAKYFGT-SDMEYGKAMF----------------  407
sp|Q3YU10|DEOB_SHISS        KPG--SLGHRET----FADIGQTLAKYFGT-SDMEYGKAMF----------------  407
sp|Q0SX28|DEOB_SHIF8 (22)   KPG--SLGHRET----FADIGQTLAKYFGT-SDMEYGKAMF----------------  407
sp|B5Y275|DEOB_KLEP3        KPG--SLGHRET----FADIGQTIAKYFGT-SDMEYGKAMF----------------  407
sp|A9MRA5|DEOB_SALAR        KPG--SLGHRET----FADIGQTLASYFGT-SPMDYGKNML----------------  407
sp|B5BAJ9|DEOB_SALPK (2)    KPG--SLGHRET----FADIGQTLATYFGT-SPMDYGKNML----------------  407
sp|C0Q7M5|DEOB_SALPC (2)    KPG--SLGHRET----FADIGQTLATYFGT-SPMDYGKNML----------------  407
sp|B5R9V1|DEOB_SALG2        KPG--SLGHRET----FADIGQTLATYFGT-SPMDYGKNML----------------  407
sp|B5FTC7|DEOB_SALDC        KPG--SLGHRET----FADIGQTLATYFGT-SPMDYGKNML----------------  407
sp|P63924|DEOB_SALTI (8)    KPG--SLGHRET----FADIGQTLATYFGT-SPMDYGKNML----------------  407
sp|C5BHJ4|DEOB_EDWI9        KPG--SLGERDT----FADIGQTVARYFGL-SPMAYGKPMF----------------  407
sp|Q6D990|DEOB_PECAS        KPG--SYGHRET----FADIGQTVAAYFGL-SPMDYGKSIL----------------  407
sp|C6DKL9|DEOB_PECCP        KPG--SYGHRET----FADIGQTVAAYFGL-SPMDYGKSIL----------------  407
sp|B4EWA2|DEOB_PROMH        KPG--SLGHRET----FADIGQTVVKYFGL-SPVEYGKAMF----------------  408
sp|Q7N931|DEOB_PHOLL        QPG--SLGHRET----FADIGQTVAKYFEL-SPMEYGKSML----------------  407
sp|A8G9H8|DEOB_SERP5        KPG--SLGHRET----FADIGQTVASYFGV-SPMDYGKSMF----------------  407
sp|A1JJ99|DEOB_YERE8        KPG--SLGHRET----FADIGQTVAKYFDL-SPMDYGKNML----------------  407
sp|A9R047|DEOB_YERPG        KPG--SLGHRET----FADIGQTVAAYFGL-SPMDYGKNML----------------  407
```

```
sp|A4TQJ1|DEOB_YERPP (4)    KPG--SLGHRET----FADIGQTVAAYFGL-SPMDYGKNML----------------                407
sp|B1JL35|DEOB_YERPY (4)    KPG--SLGHRET----FADIGQTVAAYFGL-SPMDYGKNML----------------                407
```
